# Human myelocyte and metamyelocyte-stage neutrophils suppress tumor immunity and promote cancer progression

**DOI:** 10.1101/2025.06.14.659663

**Authors:** Wei Liu, Tao Shi, Chun Lu, Keying Che, Zijian Zhang, Yuting Luo, Daniel Hirschhorn, Hanbing Wang, Shaorui Liu, Yan Wang, Shuang Liu, Haiqiao Sun, Jun Lu, Yuan Liu, Dongquan Shi, Shuai Ding, Heping Xu, Liaoxun Lu, Jianming Xu, Jun Xin, Yinming Liang, Taha Merghoub, Jia Wei, Yan Li

## Abstract

Tumor-infiltrating neutrophils (TINs) are highly heterogeneous and mostly immunosuppressive in the tumor immune microenvironment (TIME). Current biomarkers of TINs and treatment strategies targeting TINs have not yielded optimal responses in patients across cancer types. Here, we separated human and mouse neutrophils into three developmental stages, including promyelocyte (PM), myelocyte & metamyelocyte (MC & MM), and banded & segmented (BD & SC) neutrophils. Based on this separation, we observed the predominance of human but not mouse MC & MM-stage neutrophils in bone marrow (BM), which exhibit potent immunosuppressive and tumor-promoting properties. MCs & MMs also occupy the majority of TINs among patients with 17 cancer types. Moreover, through the creation of a NCG-*Gfi1*^-/-^ human immune system (HIS) mouse model, which supports efficient reconstitution of human TIN, we found a significant increase of BM MCs & MMs in tumor-bearing mice. By comparing the scRNA-seq analysis results of human neutrophils from both BM and tumors, we found CD63 and Galectin-3 distinguish MC & MM from neutrophil populations in cancer patients. Furthermore, we proposed a strategy with Flt3L to specifically induce the trans-differentiation of MCs & MMs into monocytic cells, and trigger tumor control in NCG-*Gfi1*^-/-^ HIS mice. Thus, our findings establish an essential role of human MC & MM-stage neutrophils in promoting cancer progression, and suggest their potential as targets for developing potential biomarkers and immunotherapies for cancer.

## INTRODUCTION

Neutrophils represent the most abundant immune cell population in human peripheral blood (PB) and bone marrow (BM), involved in both physiological and pathological processes^1,2^. Tumor-infiltrating neutrophils (TINs), which accumulate in a wide range of cancer types, have recently been reported to be rather heterogeneous and can exhibit both pro-tumor and anti-tumor roles^3–5^. In most cancers, TINs are found to be immunosuppressive and facilitate cancer progression via multiple means such as promotion of angiogenesis or inhibition of T cell activation^6–10^. Meanwhile, the accumulation of anti-tumorigenic neutrophils was also reported to be essential for successful T cell immunotherapies in patients^11–14^. Despite these recent efforts, the biomarkers currently reported for TINs remain limited, and effective treatment strategies targeting TINs are still lacking. This underscores the necessity to undertake more in-depth neutrophil studies in cancer.

The classification of neutrophil development based on histological staining and microscopy has existed for decades, in which neutrophils undergo promyelocyte (PM), myelocyte (MC), metamyelocyte (MM), band (BD), and segmented (SC) stages^15^. Using mass cytometry (CyTOF) and single-cell-based multi-omics sequencing, recent studies have delineated and categorized the developmental stages of human neutrophils based on specific marker expressions^16,17^. Dinh et al. divided human BM neutrophils into N1-N5 stages^16^, while Evrard et al. defined neutrophil development process from neutrophil progenitor (proNeu), neutrophil precursor (preNeu), immature neutrophils to mature neutrophils^17^. Neutrophil development has been associated with their functional roles in several disease models^18,19^. However, the dynamics and impact of neutrophil development in cancer patients remain largely unknown, which hinders the understanding of TIN heterogeneity and the development of potential TIN-targeted therapies.

Current research on TINs is primarily based on syngeneic mouse models^3–5^, which cannot accurately replicate the features and dynamics of neutrophils in human tumors, therefore limiting the translational value of their findings. Human neutrophils have a short lifespan, making it rather difficult to explore the role and function of human neutrophils in preclinical mouse models^20,21^. Recent studies by Ito and Flavell groups reported two humanized mouse models with human granulocyte colony-stimulating factor (G-CSF) knock-in and/or mouse G-CSFR knock-out, which facilitate the differentiation and reconstitution of human neutrophils in the mouse BM and organ tissues^22,23^. However, these two gene knockout/in strategies are not fully effective for eliminating mouse neutrophils in vivo, as 20-30% of circulating and 70% of BM mouse neutrophils remain, thus limiting the peripheral reconstitution of human neutrophils. As such, a humanized mouse model optimized for circulating human neutrophils is still lacking.

Here, we separated human neutrophils into three developmental stages based on reported gating strategies^15,17,24^. We found that MC & MM-stage neutrophils, which constitute the majority of neutrophils in human BM and tumors across 17 cancer types, exhibit immunosuppressive and tumor-promoting properties. By the creation of NCG- *Gfil*^-/-^ HIS mouse model which eliminates mouse neutrophil and enables efficient peripheral reconstitution of human neutrophils, we found the increase of BM MCs & MMs under tumor-bearing conditions. With scRNA-seq analysis of human BM and intratumoral neutrophils across 17 cancer types, we found CD63 and Galectin-3 distinguish MC & MM from neutrophil populations in cancer patients, and Fms-like tyrosine kinase 3 ligand (Flt3L) as a potential approach transdifferentiating MCs & MM neutrophils into monocytic cells for cancer therapy. These findings pave the way in the exploration of neutrophil development in cancer prevention, monitoring, and treatment.

## RESULTS

### MC & MM-stage neutrophils in human bone marrow are naturally immunosuppressive

We analyzed human BM and PB samples after lysing red blood cells from non-tumor donors by flow cytometry and found the expression of LOX1 and ARG1, previously reported as an immunosuppressive marker in neutrophils^25^, is markedly upregulated in BM neutrophils compared to those in PB of non-tumor donors (Fig. 1a; Supplementary information, Fig. S1a). Expression of immunosuppression related genes (*ARG1*, *LOX1*, *FATP2*, *NOX2*, *S100A8* and *S100A9*)^26,27^ is also significantly increased in BM neutrophils (Fig. 1b). Also, T cell proliferation was significantly suppressed when co-cultured with BM neutrophils (E506^-^CD45^+^SSC^hi^CD66b^+^) compared to those co-cultured with PB neutrophils or in the control group (Fig. 1c).

**Fig. 1.**
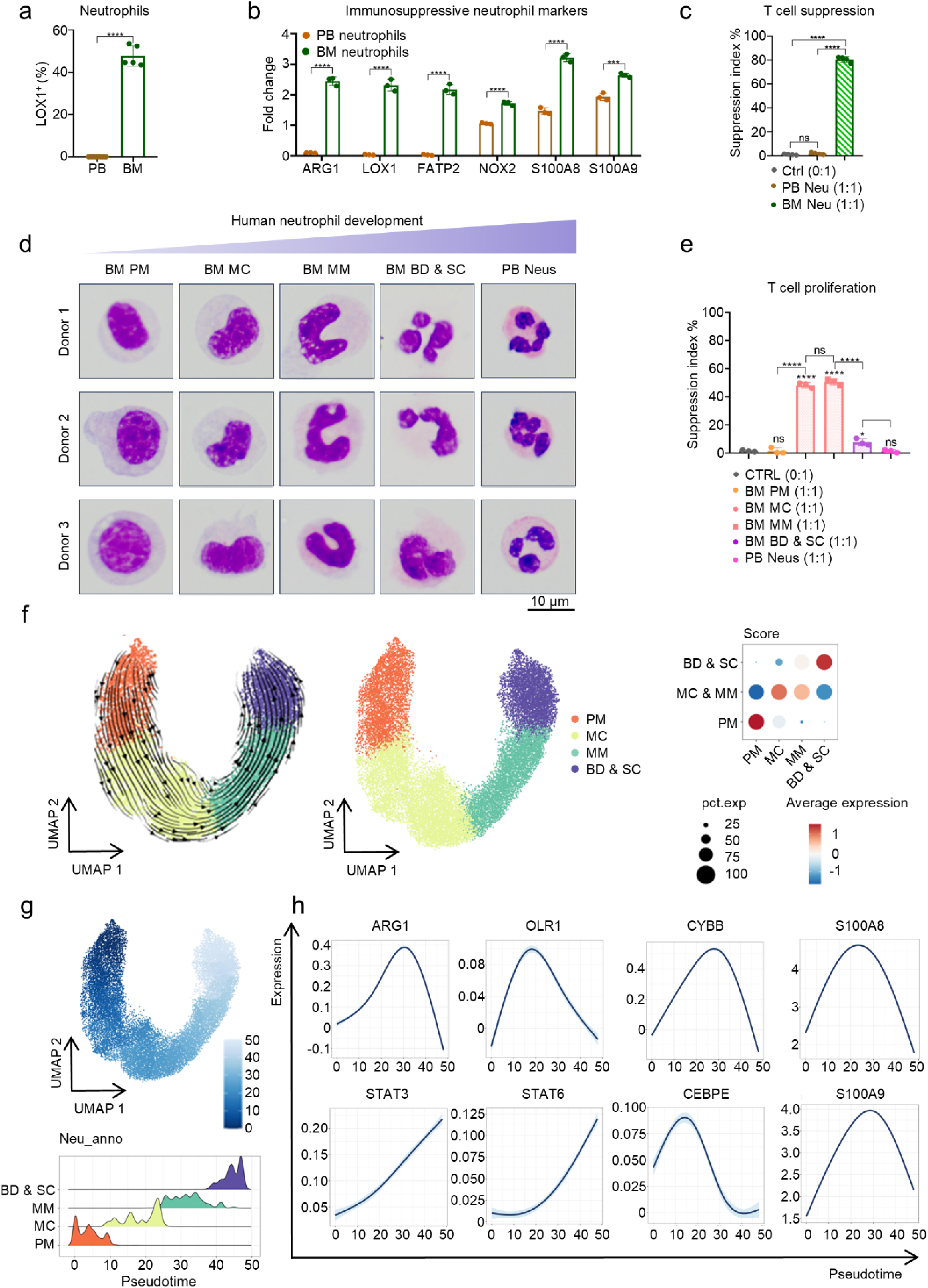
Characterization of human bone marrow neutrophils. **a** BM neutrophils from non-tumor donors were collected for subsequent analysis, and the proportion of LOX1^+^ / neutrophils (CD11b^+^CD66b^+^) in PB (*n* = 16) and BM (*n* = 5) and the proportion of ARG1^+^ / neutrophils (CD11b^+^CD66b^+^) in PB (*n* = 4) and BM (*n* = 4) of non-tumor donors was determined by flow cytometry. **b** mRNA expression of immunosuppressive genes in neutrophils from PB and BM of non-tumor donors (*n* = 3) analyzed by q-PCR. Reference gene: human *GAPDH*; endogenous control: human total BM cells. **c** Suppression index of human T cell proliferation co-cultured with neutrophils sorted from BM or PB of non-tumor donors (*n* = 4), and the neutrophils : T cells ratio is 1 : 1. **d** Giemsa staining of BM PM∼BD & SC-stage neutrophils and PB neutrophils of non-tumor donors (*n* = 3). **e** T cell suppression assay of BM PM∼BD & SC-stage neutrophils and PB neutrophils of non-tumor donors (*n* = 3). **f** We analyzed the scRNA-seq dataset of human BM neutrophils, and the scTour analysis and the score of human BM neutrophils at different developmental stages was shown. **g** Pseudotime analysis of human BM neutrophils. **h** Expression levels of immunosuppressive markers of human BM neutrophils at different stages. All data are mean ± SD and were analyzed by two-tailed, unpaired Student’s t-test (\**P* < 0.05, \*\**P* < 0.01, \*\*\**P* < 0.001, \*\*\*\**P* < 0.0001).

Given the immunosuppressive properties of human BM neutrophil, we sought to further characterize their developmental stages. We sorted human BM neutrophils at various stages^15^, and results of Giemsa staining showed a progressive change in nuclear morphology during human BM neutrophil development, transitioning from circular (PM, CD66b^+^CD11b^-^) to horseshoe-like (MC, CD66b^+^CD11b^+^CD10^-^CD16^-^; MM, CD66b^+^CD11b^+^CD10^-^CD16^lo^), band and segmented (BD & SC, CD66b^+^CD11b^+^CD10^lo/+^CD16^+^) shapes (Fig. 1d). Importantly, the T cell suppression assay demonstrated that MC & MM neutrophils exhibit the strongest immunosuppressive properties, whereas PM or BD & SC-stage neutrophils do not suppress T cell proliferation (Fig. 1e). We next aimed to analyzed the single-cell RNA-seq dataset (from GSE130756 and the website https://usegalaxy.eu/published/history?id-008000b0c3d8bb03) of human BM neutrophils. However, we found that markers routinely used for neutrophil identification in flow cytometry, such as CD11b (encoded by *ITGAM*), CD66b (encoded by *CEACAM8*), CD10 (encoded by *MME*), and CD16 (encoded by *FCGR3A*), exhibit minimal expression in single-cell transcriptomic datasets, rendering them unsuitable for neutrophil subpopulation clustering in transcriptomic analyses (Supplementary information, Fig. S1b). To address this, we chose to sort neutrophils based on their differentiation stage and then perform bulk RNA-seq. This approach allowed us to achieve a much higher sequencing depth of neutrophils compared to scRNA-seq, enabling us to detect low-abundance transcripts and subtle changes in gene expression that might be missed in single-cell datasets. We performed bulk RNA-seq on monocytes, PM, MC & MM, and BD & SC neutrophils sorted from human BM (as described above), with the aim to establish gene sets specific for human neutrophils at various developmental stages (Supplementary information, Table S1, Score_genes.xlsx). The Principal Component Analysis (PCA) analysis showed that these 4 populations were clearly separated (Supplementary information, Fig. S1c). We then analyzed the differential expressed genes (DEGs) of each myeloid cell population. After overlapping neutrophil DEGs (identified by us) with those previously reported by Calzetti et al. and Grassi et al.^28,29^, 671 genes were identified as human neutrophil-specific genes (Supplementary information, Fig. S1d). These 671 genes were then divided into three categories, which were specifically highly expressed at the PM / MC & MM / BD & SC stages, respectively (Supplementary information, Table S2, DEGs_overlap.xlsx). Next, we applied PM, MC & MM and BD & SC scores (calculated based on the expression of DEGs specific for each developmental stage) to annotate human neutrophils from the scRNA-seq data, and defined PM to BD & SC stages using scTour analysis and Neutrophil scores (Fig. 1f). Pseudotime analysis confirmed that these populations followed a continuous differentiation trajectory from PM to BD & SC stage (Fig. 1g). We also examined the expression of polymorphonuclear myeloid-derived suppressor cell (PMN MDSC)-related markers^26,27^ at different neutrophil stages and found that key PMN MDSC-related markers, such as *ARG1, OLR1* (*LOX1*) and *CYBB*, were predominantly expressed in MC & MM neutrophils (Fig. 1h). To strengthen the generalizability of our findings, we also analyzed another recently published scRNA-seq dataset of human BM neutrophils^18^. And we found that the PM to BD & SC stages could still be clearly defined on the data of Montaldo et al using Neutrophil scores analysis (Supplementary information, Fig. S1e). Pseudotime analysis estimated by scTour and monocle3 further confirmed that these populations followed a continuous differentiation trajectory from PM to BD & SC (Supplementary information, Fig. S1f). The PMN MDSC-related markers are also predominantly highly expressed during the MC & MM stages (Supplementary information, Fig. S1f). Taken together, these data demonstrated the unique immunosuppressive property of MC & MM neutrophils during development in BM.

Recent papers have generated distinct nomenclature for human neutrophil developmental staging, such as N1-N5; Pro Neu1-Mature Neu^16,17^. Dinh et al. initially performed mass cytometry (CyTOF) analysis on CD45^+^ cells from healthy human BM using a 35-marker panel, followed by UMAP cluster analysis and cumulative distribution function (CDF) curve optimization to resolve five distinct neutrophil subsets (N1∼N5)^16^. Also, by integrating scRNA-seq datasets of mouse granulocyte-monocyte progenitors (GMPs), Kwok et al. identified a distinct subpopulation highly expressing neutrophil-specific genes (e.g., *Gfi1*, *S100a8*). Subsequent InfinityFlow surface marker screening revealed unique immunophenotypic signatures, and in vitro differentiation assays demonstrated exclusive neutrophil commitment (no monocytic differentiation under high M-CSF conditions), leading to the designation of this population as Pro Neu1. Monocle pseudotime analysis and in vivo transfer experiments further delineated a hierarchical maturation trajectory: mouse Pro Neu1, Pro Neu2, Pre Neu, immature and mature neutrophils^17^. And we next tried to map the classical PM-BD & SC neutrophils to these populations to facilitate the identification of MM & MC equivalent populations. We conducted flow cytometry analysis of human BM neutrophils according to three reported human neutrophil gating strategies^15,17,24^(Supplementary information, Fig. S2a). The same BM samples analyzed by three different gating strategies were visualized on t-distributed Stochastic Neighbor Embedding (t-SNE) plots, and we found overlaps of neutrophil clusters across different t-SNE plots (PM overlaps with N1, Pro Neu1 & Pro Neu2; MC & MM overlap with N2 & N3, Pre Neu & Immature Neu; BD & SC overlap with N4 & N5, Mature Neu) (Supplementary information, Fig. S2b, c). Thus, human N2 & N3 and Pre Neu & Immature Neu likely to suppress T cells as MM & MC neutrophils.

### Human MC & MM-stage neutrophils are long-lived and tumor-promoting

Next, we compared the longevity of human neutrophil at various stages, and their immunosuppressive function via co-culture with T cells. We sorted human PB neutrophils and BM neutrophil at various stages (PM: CD66b^+^CD11b^-^; MC & MM: CD66b^+^CD11b^+^CD10^-^CD16^-/lo^; BD & SC: CD66b^+^CD11b^+^CD10^lo/+^CD16^+^; PB Neus: SSC^hi^CD66b^+^), and analyze their survival time ex vivo. We observed that MC & MM neutrophils have the longest survival time, with majority of MC & MM neutrophils still alive on Day 2 (Fig. 2a). We also documented the apoptosis of CD45^+^CD66b^+^ cell populations during this process, and the results demonstrated that MC & MM neutrophils exhibited greater apoptosis resistance (Supplementary information, Fig. S3a, b). Moreover, we found that the proliferation of human T cells was only suppressed when co-cultured with MC & MM neutrophils across all three timepoints (Fig. 2b). In addition, we assessed the proliferative potential of neutrophils at various stage by detecting Ki67 expression levels. The results revealed that the proliferative potential of neutrophils rapidly diminished with developmental maturation, and low Ki67 expression was observed in MC & MM, BD & SC, and PB Neu (Supplementary information, Fig. S3c, d). Therefore, these results suggested that in addition to the expression of PMN MDSC-related genes, the long lifespan also contributes to the immunosuppressive property of human MC & MM neutrophils. To further examine whether BM MCs & MMs promote tumor progression in vivo, we constructed the NOD/ShiLtJGpt-Prkdc^em26Cd52^Il2rg^em26Cd22^/Gpt (NCG) human immune system (HIS) mouse model, which supports *de novo* human T cell engraftment and is widely applied as a preclinical model for tumor immunotherapy^30–32^. MC & MM- or BD & SC-stage neutrophils, sorted from human BM (as described above), were intravenously (i.v.) transferred into A375 (human melanoma cell line)-bearing NCG HIS mice on Day 13 and Day 21 following tumor inoculation (Fig. 2c). Consequently, tumor burden increased significantly in mice transferred with MC & MM neutrophils (Fig. 2d, e). Also, we observed that HIS mice transferred with human MCs & MMs have decreased human T cells in the PB and tumor tissues compared to other two groups (Fig. 2f). And qPCR analysis of bulk tumor tissues revealed that MC & MM neutrophil transfer significantly upregulated T cell exhaustion-associated markers (Supplementary information, Fig. S3e). Collectively, these results indicate that MC & MM-stage neutrophils, which predominate in human BM, are long-lived and tumor-promoting.

**Fig. 2.**
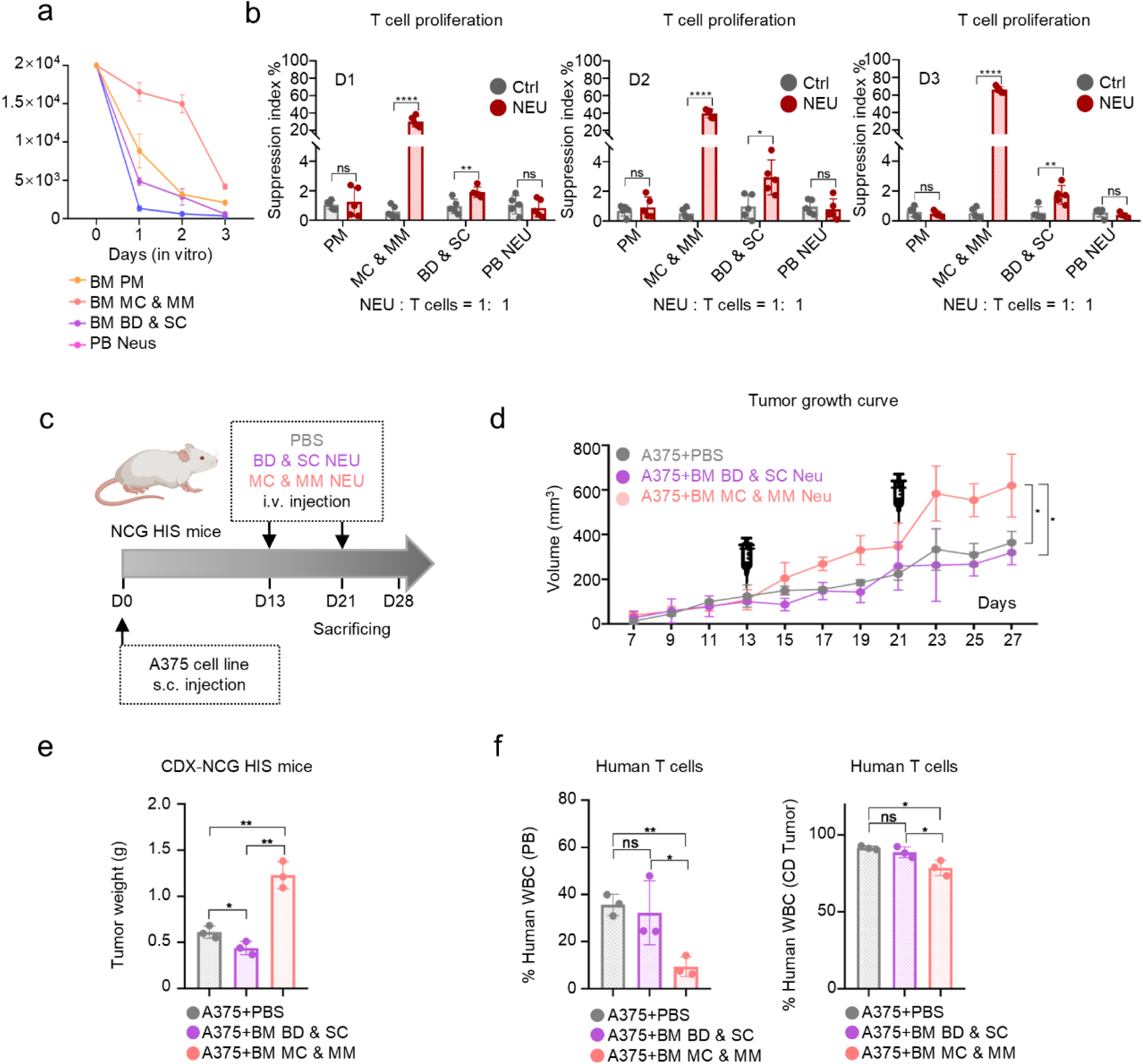
MC & MM-stage neutrophils in human bone marrow are long-lived and tumor-promoting. **a** Number of alive human neutrophils at various developmental stages ex vivo (*n* = 5). **b** Suppression index of human T cell proliferation co-cultured with human PB neutrophils or BM neutrophils at different developmental stages at the ratio of 1:1 (neutrophil : T cell). The timepoints of the assay include Day1, Day2, and Day3. **c** Schematic of NCG HIS mice (s.c. implantation of A375 cells) with human BM MC & MM or BD & SC-stage neutrophil transfer (2.5×10^6^ neutrophils i.v. injection per dose at Day 13 and 21). **d** Tumor growth of NCG HIS mice with human BM MC & MM or BD & SC-stage neutrophil transfer (*n* = 3). **e** Tumor weight of NCG HIS mice with human BM MC & MM or BD & SC-stage neutrophil transfer (*n* = 3). **f** Proportion of human T cells in the PB or tumor of HIS mice from each group (*n* = 3). All data are mean ± SD and were analyzed by two-tailed, unpaired Student’s t-test (\**P* < 0.05, \*\**P* < 0.01, \*\*\**P* < 0.001, \*\*\*\**P* < 0.0001).

### The predominance and immunosuppressive property of MC & MM-stage neutrophils in BM are specific to human neutrophil development

We next explored whether the predominance and immunosuppressive property of MC & MM neutrophils could also be observed in mouse BM neutrophils. We performed flow cytometry and scRNA-seq analysis on BM cells isolated from 12-week-old C57BL/6J mice, and the results showed that neutrophils are also the dominant immune cell population in mouse BM (Supplementary information, Fig. S4a, b). We divided mouse neutrophil populations into G1∼G5 by scTour and pseudotime analysis (Supplementary information, Fig. S4c, d), and then we observed that gene markers associated with immunosuppression are not enriched in G1 ∼G5 clusters (Supplementary information, Fig. S4e, f). We also separated mouse BM neutrophils into the four stages through flow cytometry (PM: Lin^-^Ly6G^-^CD117^lo^; MC: Lin^-^Ly6G^lo^CD117^lo^; MM: Lin^-^ Ly6G^lo/+^CD117^lo/-^; BD & SC: Lin^-^Ly6G^+^CD117^-^) (Supplementary information, Fig. S5a). Different from human samples, we found neutrophils were predominantly in BD & SC stage in the PB, spleen, or BM of mice (Supplementary information, Fig. S5b). Meanwhile, results from the T cell suppression assay demonstrated that all the subsets of mouse neutrophils, including PM, MC, MM, BD & SC and PB neutrophils, did not affect mouse T cell proliferation (Supplementary information, Fig. S5c). Therefore, these results suggest that mouse neutrophils (including PM, MC, MM and BD & SC) are predominant in BM, but not inherently immunosuppressive under physiological conditions. However, we found that while neutrophils from the BM and spleen of WT mice do not suppress T cell proliferation, neutrophils sorted from mouse tumor tissue displayed strong suppressive function (Supplementary information, Fig. S6), indicating that the acquisition of immunosuppressive properties in mouse neutrophils is independent of their developmental stages.

As human neutrophils have been shown to develop in BM of HIS mice^22,23^, we next evaluated whether human MC & MM-stage neutrophils developed in mouse BM environment could maintain their predominance and immunosuppression. We constructed the HIS mouse model and detected the proportion, maturity, and immunosuppression of human neutrophils in the bone marrow of HIS mice at 10 weeks. As shown in Supplementary information, Fig. S7a, b, we found a SSC^hi^ population, which mostly consisted of human MC & MM-stage neutrophils, present only in the BM of NCG HIS mice and non-tumor human donors. Non-neutrophilic myeloid cells (Lin^-^CD66b^-^CD11b^+^) and neutrophils (Lin^-^SSC^hi^CD66b^+^) were sorted from the BM of NCG HIS mice (Supplementary information, Fig. S7c). Consistent with our findings in human BM neutrophils, compared to non-neutrophilic myeloid cells, we observed that sorted human neutrophils from NCG HIS mice suppress T cell proliferation and have significantly increased immunosuppression-related gene expression (Supplementary information, Fig. S7d-f). We then sorted neutrophils at MC & MM, BD & SC stages of development from NCG HIS mice, and validated the dominance of MC & MM-stage neutrophils in BM and their capability to inhibit T cell proliferation (Supplementary information, Fig. S7g, h). Thus, results comparing mouse and human BM neutrophil development and function supported that the predominance and immunosuppressive properties of MC & MM neutrophils in BM are specific to human neutrophil development.

### MC & MM-stage neutrophils constitute the majority of TINs across cancer types

As human MC & MM-stage neutrophils demonstrated immunosuppressive and tumor-promoting property in BM, we continued to analyze the distribution and function of MC & MM neutrophils in PB and tumor of cancer patients. We first found a significant increase of MC & MM neutrophils and decrease of BD & SC neutrophils in the PB of gastric cancer (GC) patients (Fig. 3a). A high prevalence of MC & MM neutrophils (about 80%) was also observed within the primary and peri-tumor tissues collected from GC patients (Fig. 3b), which significantly inhibited T cell proliferation after in vitro co-culture (Fig. 3c). In addition, the predominance of MC & MM neutrophils was further validated in tumor tissues of pancreatic ductal adenocarcinoma (PDAC), bladder cancer (BLCA) and melanoma patients (Fig. 3d).

**Fig. 3.**
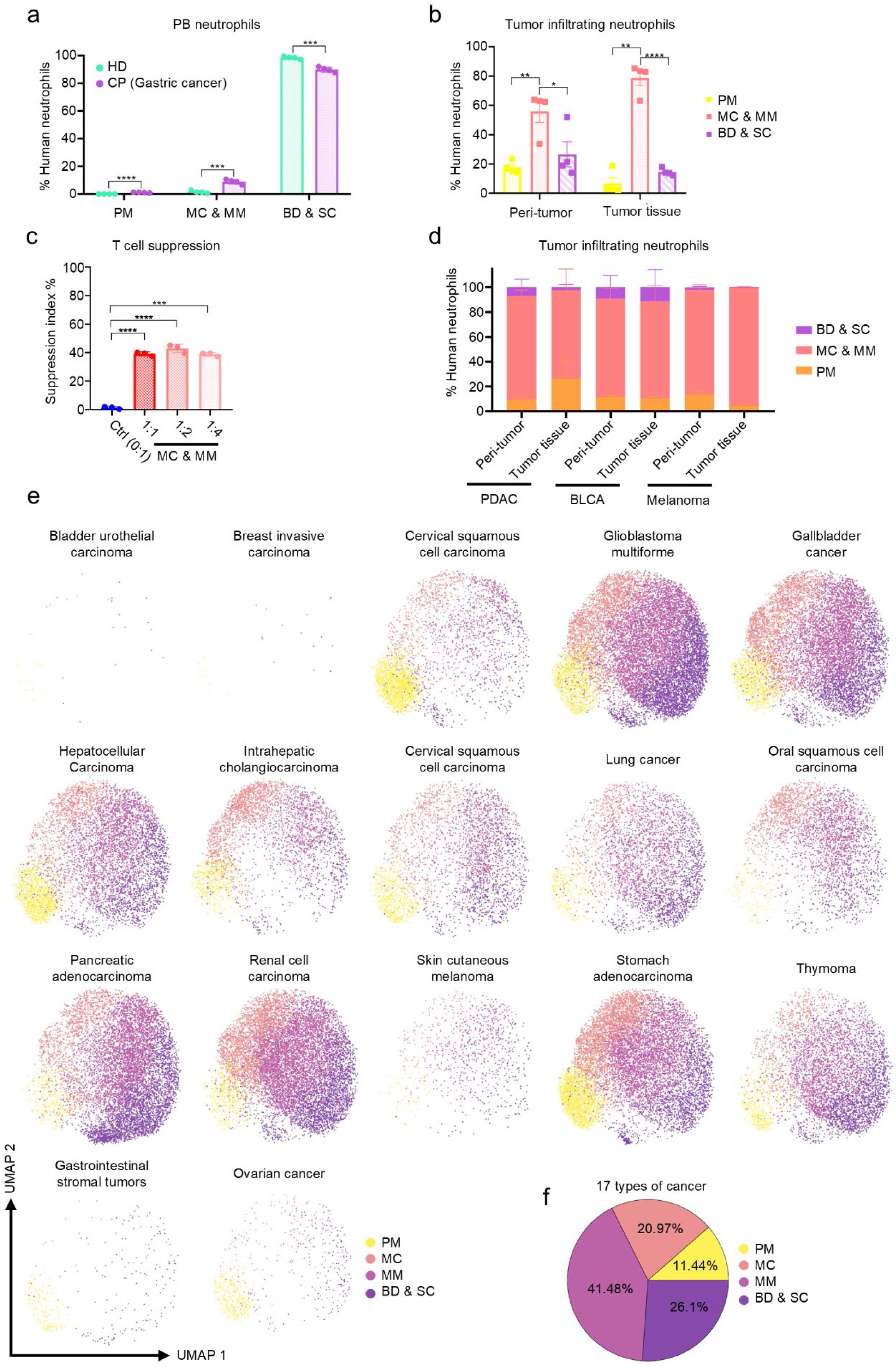
MC & MM neutrophils constitute the majority of TINs across tumor types. **a** Proportions of human neutrophils at different developmental stages (PM: CD66b^+^CD11b^-^), MC & MM: CD66b^+^CD11b^+^CD10^-^CD16^-/lo^), and BD & SC: CD66b^+^CD11b^+^CD10^lo/+^CD16^+^) in PB of healthy donors (HD, *n* = 6) and gastric cancer patients (CP, *n* = 4). **b** Proportions of neutrophils at different developmental stages in peri-tumor and tumor tissues of GC patients determined by flow cytometry (*n* = 4). **c** Suppression index of human T cell proliferation co-cultured with MC & MM neutrophils (CD66b^+^CD11b^-^CD10^-^CD16^-/lo^) sorted from tumors of GC patients at different E:T ratios (*n* = 3). **d** Proportions of neutrophils at different developmental stages in peri-tumor and tumor tissues of PDAC, BLCA and melanoma patients determined by flow cytometry (*n* = 2 for each cancer type). **e** UMAP clustering of TINs at different developmental stages in each cancer type from reported sc-RNAseq datasets of 17 cancer types^14^. 17 cancer types include bladder urothelial carcinoma (BLCA), breast invasive carcinoma (BRCA), cervical squamous cell carcinoma and endocervical adenocarcinoma (CESC), colon adenocarcinoma (COAD), gallbladder cancer (GBC), gastrointestinal stromal tumor (GIST), hepatocellular carcinoma (HCC), intrahepatic cholangiocarcinoma (ICC), lung cancer (LC), lower grade glioma (LGG), oral squamous cell carcinoma (OSCC), ovarian serous cystadenocarcinoma (OV), pancreatic adenocarcinoma (PAAD), renal cell carcinoma (RCC), skin cutaneous melanoma (SKCM), stomach adenocarcinoma (STAD), thymoma (THYM). **f** Proportions of TINs at different developmental stages in 17 types of tumors. All data are mean ± SD and were analyzed by two-tailed, unpaired Student’s t-test (\**P* < 0.05, \*\**P* < 0.01, \*\*\**P* < 0.001, \*\*\*\**P* < 0.0001).

We next aimed to extend our findings to other types of cancer with the analysis of external scRNA-seq datasets. We applied neutrophil scores we generated (Supplementary information, Fig. S1c, d) to annotate TINs into 4 populations (PM, MC, MM and BD & SC) with the scRNA-seq data from 17 cancer types^14^, and confirmed the predominance of MC & MM in TINs across majority of tumor types (Fig. 3e, f; Supplementary information, Fig. S8a-c; Supplementary information, Table S3).

To investigate whether the presence of tumor alters the composition of mouse neutrophils in mice and favor the accumulation of MC & MM neutrophils like in human settings, we established a syngeneic GC mouse model via subcutaneously (s.c.) injection of mouse forestomach carcinoma (MFC) cells. As shown in Supplementary information, Fig. S9a-e, we found that the proportion of MC & MM neutrophils in the BM was increased, while BD & SC rather than MC & MM neutrophils were increased in the PB of tumor-bearing mice. However, unlike human, the proportion of BD & SC neutrophils was relatively high among TINs in CDX and spontaneous tumor models (Supplementary information, Fig. S9f, g), supporting the necessity to develop a suitable humanized mouse model to overcome the impact of species differences.

### BM MC & MM-stage neutrophil increases in response to tumor stimulation in human and NCG-*Gfi1*^-/-^HIS mice

The predominance of MC & MM-stage neutrophils in healthy human BM and tumors prompted us to further investigated whether the composition of BM neutrophils changes under the influence of tumor cells. Current research of neutrophils in the TIME primarily relies on syngeneic mouse models^5^. However, given the difference of neutrophil composition between mouse and human described above, optimized HIS mouse models for human TIN research in vivo are needed, which should accurately resemble the dynamics of neutrophil distribution in cancer patients. Although the NCG HIS mice support the development of human neutrophils in the BM, the peripheral reconstitution of human neutrophils is extremely low (3.49 ± 1.40%, Fig. 4a-d), making this model impossible to interrogate the dynamics of human peripheral neutrophils. We hypothesized that the poor engraftment of peripheral human neutrophils is due to the competition with intact mouse neutrophil compartment for cytokines cross-species active from mouse to human in NCG mice. To test this hypothesis, we depleted mouse neutrophils by anti-Ly6G antibodies (Supplementary information, Fig. S10a), and observed the elevated serum concentrations of mouse G-CSF, which can cross-react on human cells, accompanied by a significant increase of human neutrophils in PB and BM of NCG HIS mice (Supplementary information, Fig. S10b, c). Therefore, we employed CRISPR-Cas9 technology to knock out the Growth factor independent 1 transcriptional repressor gene (*Gfi1*, reported as a key transcription factor in neutrophil development^33,34^) in NOD mice and crossed to NCG background to block mouse neutrophil development (Supplementary information, Fig. S11a, b). As shown in Supplementary information, Fig. S11c, d, numbers of mouse WBCs and hematopoietic progenitor cells were dramatically reduced in NCG-*Gfi1*^-/-^ mice compared to NCG and NCG-*Gfi1*^+/-^mice. Importantly, NCG-*Gfi1*^-/-^ mice demonstrated a phenotype of severe neutropenia (Supplementary information, Fig. S11e-f). Consequently, after humanization of NCG-*Gfi1*^-/-^ mice, significant increases of human WBCs and neutrophils were observed in the PB, BM, and SP of NCG-*Gfi1*^-/-^ HIS mice (15 ± 1.21%, Fig. 4a-d). We also performed Giemsa staining on sorted human neutrophils at various developmental stages from the BM and PB of NCG-*Gfi1^-/-^* HIS mice. Human neutrophils differentiated in NCG-*Gfi1^-/-^* HIS mice still underwent progressive morphological transitions of their nuclei during maturation, from circular to horseshoe-shaped, banded, and ultimately segmented forms (Supplementary information, Fig. S12a), though not as highly segmented as those in human (Fig. 1d). To evaluate the functionality of circulating human neutrophils developed in NCG-*Gfi1^-/-^* HIS mice, we performed the transwell migration assay of human neutrophils sorted from PB of NCG-*Gfi1*^-/-^ HIS mice. Sorted neutrophils were cultured with serum collected from NCG-*Gfi1*^-/-^ non-HIS mice hydrodynamically injected with human CXCL8 plasmid. We observed that the number of neutrophils migrated into the CXCL8-enriched medium was significantly higher than in the control group (Supplementary information, Fig. S12b), demonstrating the chemotactic function of human neutrophils in NCG-*Gfi1*^-/-^ HIS mice. We also sorted human neutrophils from PB of NCG-*Gfi1*^-/-^ HIS mice and cultured them with 100 nM Phorbol 12-myristate 13-acetate (PMA) for 2 h. Giemsa staining and qPCR results showed that these human neutrophils formed neutrophil extracellular traps (NETs), and the expressions of PAD4, MPO and PKCβ were significantly up-regulated after PMA treatment (Supplementary information, Fig. S12c). We also analyzed the reconstitution of human T, B and monocytes in NCG-*Gfi1*^-/-^ HIS mice, and observed an obvious increase of monocytes in PB and spleen of NCG-*Gfi1^-/-^* HIS mice compared to NCG HIS mice (Supplementary information, Fig. S12d). In addition, we observed the reduced proportions of human T and B cells observed in the PB of NCG-*Gfi1^-/-^* HIS mice (Supplementary information, Fig. S12d), probably due to the concomitant increase of human monocytes and neutrophils. Therefore, based on all these observations, we generated the NCG-*Gfi1*^-/-^ HIS mice with high levels of human peripheral neutrophils.

**Fig. 4.**
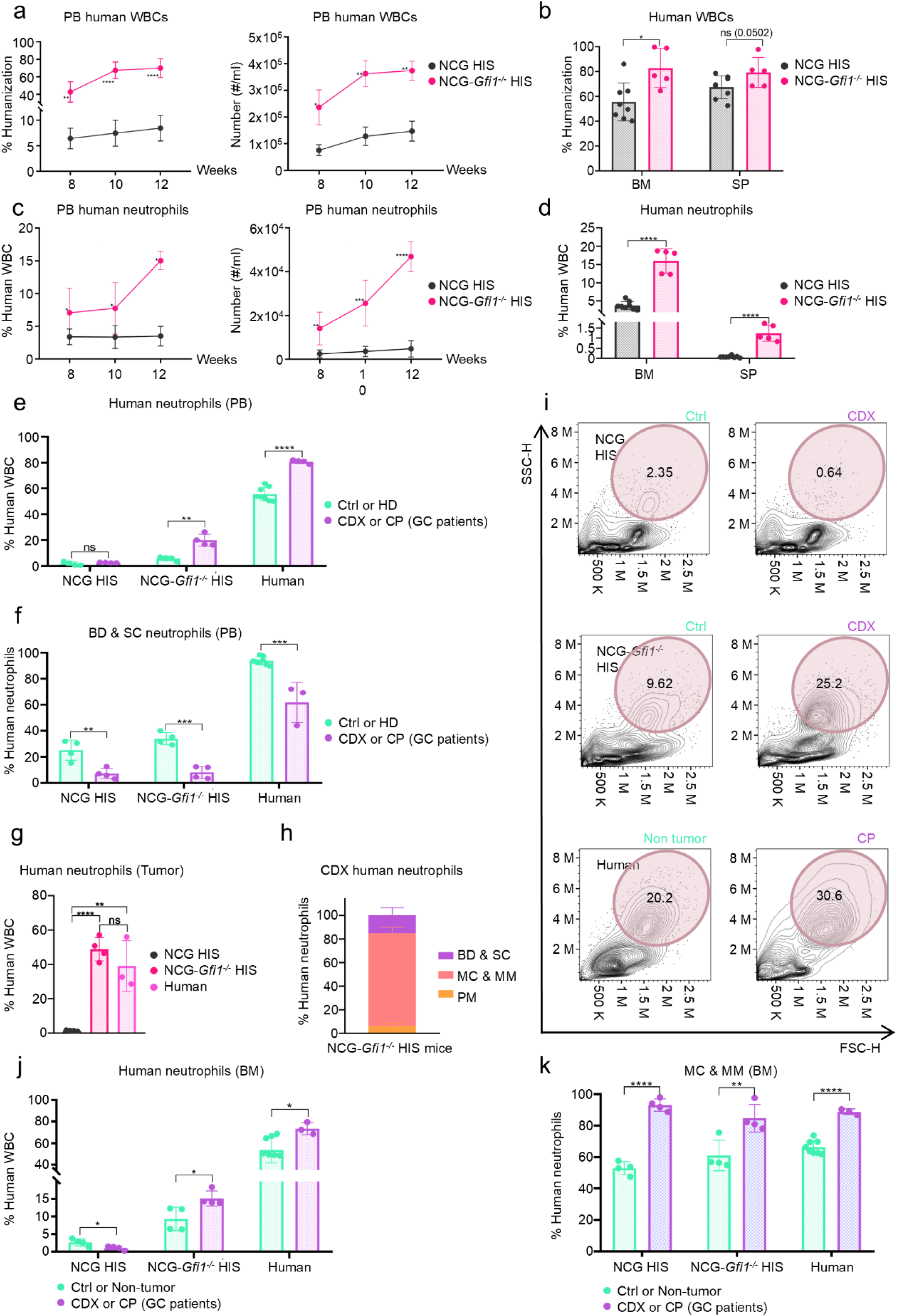
The proportion of MC & MM neutrophils is increased in the bone marrow of cancer patients and tumor-bearing NCG-*Gfi1*^-/-^ HIS mice. **a** The humanization ratio (human immune cell / total immune cells) and number of human WBCs in the PB of NCG (*n* = 8) or NCG-*Gfi1*^-/-^ HIS mice (*n* = 5) at different timepoints. **b** The humanization ratio in the BM and SP of NCG or NCG-*Gfi1*^-/-^ HIS mice. **c** Proportions and number of human neutrophils among WBCs in the PB of NCG or NCG-*Gfi1*^-/-^ HIS mice at different timepoints. **d** Proportions of human neutrophils among WBCs in BM and SP of NCG or NCG-*Gfi1*^-/-^ HIS mice. **e** Proportions of human neutrophils in the PB of NCG HIS mice, NCG-*Gfi1*^-/-^ HIS mice (Ctrl, *n* = 4; CDX, *n* = 4), and humans (HD *n* = 8; CP, gastric cancer patients, *n* = 4). **f** Proportions of human BD & SC neutrophils in the PB of NCG HIS mice, NCG-*Gfi1*^-/-^ HIS mice (Ctrl, *n* = 4; CDX, *n* = 4), and humans (HD, *n* = 8; CP, *n* = 3). **g** Proportions of human neutrophils in the tumor tissues of NCG HIS mice, NCG-*Gfi1*^-/-^ HIS mice (*n* = 4), and cancer patients (*n* = 3). **h** Proportions of human neutrophils at different developmental stages in the tumor tissues of NCG-*Gfi1*^-/-^ HIS mice (*n* = 4). **i** Representative flow cytometry plots of BM cells in NCG or NCG-*Gfi1*^-/-^ HIS mice (with or without tumor), Non-tumor donors, and cancer patients. **j** Proportions of human neutrophils in the BM of NCG HIS mice, NCG-*Gfi1*^-/-^ HIS mice (Ctrl, *n* = 4; CDX, *n* = 4), and humans (non-tumor, *n* = 8; CP, *n* = 3). **k** Proportions of human MC & MM neutrophils in the BM of NCG HIS mice, NCG-*Gfi1*^-/-^ HIS mice (Ctrl, *n* = 4; CDX, *n* = 4), and humans (Non-tumor, *n* = 8; CP, *n* = 3). All data are mean ± SD and were analyzed by two-tailed, unpaired Student’s t-test (\**P* < 0.05, \*\**P* < 0.01, \*\*\**P* < 0.001, \*\*\*\**P* < 0.0001).

With the development of NCG-*Gfi1*^-/-^ HIS mice, we were able to analyze the composition of TIN and BM neutrophils simultaneously in the melanoma A375 cell derived xenograft (CDX) HIS mice. As shown in Fig. 4e, f, the presence of tumors resulted in a significant increase of human total neutrophils, and decrease of BD & SC neutrophils in the PB of NCG-*Gfi1*^-/-^ HIS mice and gastric cancer patients, but not in NCG HIS mice due to the lack of peripheral neutrophils. Also, like in gastric cancer patients, human neutrophils comprised about half of WBCs in the TIME of the CDX-NCG-*Gfi1*^-/-^ HIS mice, while there was nearly no human TIN in CDX-NCG HIS mice (Fig. 4g), probably due to the competition of tumor-derived cytokines with mouse neutrophils. Consistently, MC & MM neutrophils consisted of almost 80% of TINs in NCG-*Gfi1*^-/-^ HIS mice, similar to the composition in cancer patients shown in Figure 2 (Fig. 4h). Of note, we found that human SSC^hi^ neutrophils increase in the BM (Fig. 4i, j) of CDX-NCG-*Gfi1*^-/-^ HIS mice and cancer patients, in contrast to the decrease of human BM neutrophils in CDX-NCG HIS mice. Moreover, significantly increased proportion of MC & MM neutrophils (about 80% among total human neutrophils) in the BM of CDX-HIS mice and cancer patients were observed (Fig. 4k). Together, these results demonstrated the similarity between human and NCG-*Gfi1*^-/-^ HIS mice, and suggested that tumor derived signals expand human MC & MM neutrophils in BM remotely.

### CD63 and Galectin-3 distinguish MC & MM from neutrophil populations in cancer patients

Considering the increase of human MC & MM-stage neutrophils in both BM and tumors of cancer patients, we aimed to identify markers that can precisely characterize this neutrophil subset to develop targeted therapy. Human WBCs (E506^-^ mCD45^-^ hCD45^+^) from the BM of NCG HIS and NCG-*Gfi1*^-/-^ HIS mice were sorted for scRNA-seq and UMAP clustering plots were shown (Supplementary information, Fig. S13a, b). We observed a higher proportion of human neutrophils in the BM of NCG-*Gfi1*^-/-^ HIS mice compared to NCG HIS mice (Fig. 5a). Based on the neutrophil scores we generated above, we divided human neutrophil subsets into PM, MC, MM and BD & SC stages in the BM of NCG HIS and NCG-*Gfi1*^-/-^ HIS mice (Supplementary information, Fig. S13c, d). By conducting the pseudotime analysis, we found consistent gene expression profile between neutrophils in the BM of NCG-*Gfi1*^-/-^ HIS mice and human BM (Supplementary information, Fig. S13e, f). Together with previous the classification of neutrophil clusters in 17 types of human cancers by scoring (Supplementary information, Fig. S8b), we identified the differential expression genes (DEGs) of MC & MM neutrophils in human tumors (302 genes), human BM (203 genes), NCG HIS mice BM (212 genes) and NCG-*Gfi1*^-/-^ HIS mice BM (439 genes), and then we overlapped the DEGs of MC & MM neutrophils identified from the four scRNA-seq datasets. Four genes encoding intracellular proteins (*CTSD*, *FLNA*, *FGR*, and *GRN*) and four genes encoding cell surface markers (*FCER1G*, *CD63*, *LGALS3*, and *CYBB*), were found to be shared in all four datasets (Fig. 5b).

**Fig. 5.**
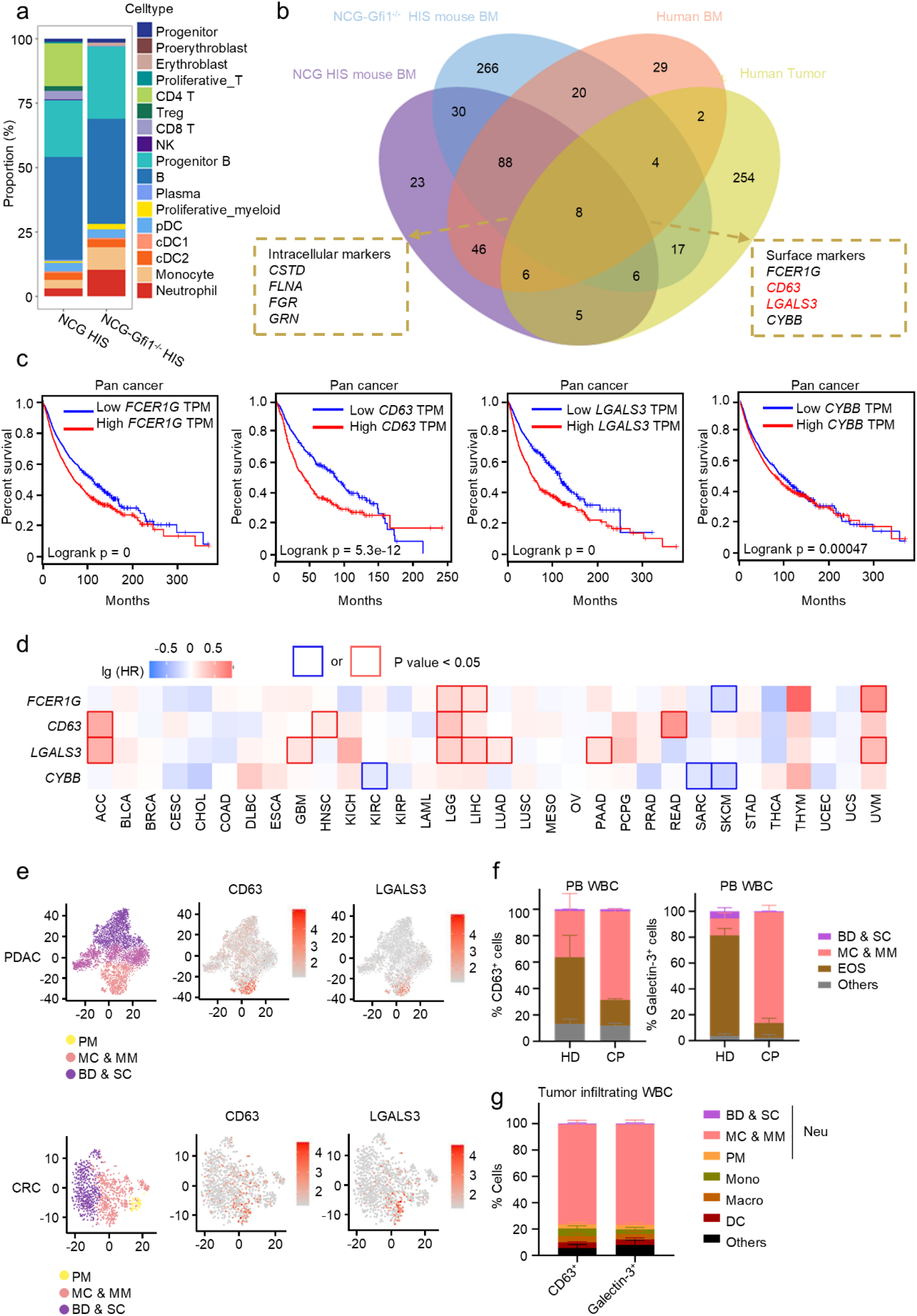
CD63 and Galectin-3 distinguish MC & MM from neutrophil populations in cancer patients. **a** The proportions of immune cell subsets in the BM of NCG HIS mice and NCG-*Gfil*^-/-^ HIS mice based on scRNA-seq and UMAP clustering. **b** Venn diagram depicting DEGs of human MC & MM neutrophils from the BM of NCG HIS mice and NCG-*Gfil*^-/-^ HIS mice and other reported datasets (including human BM ^77,78^ and 17 types of tumors ^14^). **c** Survival analysis of patients with 32 cancer types (pan-cancer) from TCGA with low or high *FCER1G*, *CD63*, *LGALS3*, or *CYBB* expression. **d** Heatmap depicting the association between gene marker expression and hazard ratio (HR) of patients with 32 cancer types from TCGA. Color scales represent the log (HR) value across cancer types. **e** *CD63* and *LGALS3* expression in TINs at different developmental stages from reported sc-RNAseq datasets of pancreatic and colorectal cancer ^35,36^. **f** Proportions of eosinophils (EOS), BD & SC and MC & MM neutrophils in PB CD63^+^ or Galectin-3^+^ (encoded by *LGALS3*) cells of healthy donors (HD, *n* = 4) and gastric cancer patients (CP, *n* = 4) determined by flow cytometry. **g** Composition of CD63^+^ or Galectin-3^+^ cells in tumor tissues of gastric cancer patients (*n* = 5). All data are mean ± SD and were analyzed by two-tailed, unpaired Student’s t-test. Survival curves were analyzed by Log-rank test (\**P* < 0.05, \*\**P* < 0.01, \*\*\**P* < 0.001, \*\*\*\**P* < 0.0001).

We investigated further on the value of 4 cell surface genes as a biomarker for MC & MM neutrophils. All 4 markers were significantly and positively associated with worse survival outcomes of patients across cancer types (32 cancer types included in pan-cancer from TCGA) (Fig. 5c; Supplementary information, Fig. S14). By conducting the univariate Cox analysis among 32 cancer types from TCGA databases, we found *CD63* and *LGALS3* gene expressions were frequently associated with higher risk of patient mortality (HR > 1, Fig. 5d), while the expression of *FCER1G* and *CYBB* did not show a consistently significant correlation across different cancer types (Fig. 5d). Subsequently, we focused on *CD63* and *LGALS3*, and validated their expression on MC & MM neutrophils among TINs of PDAC and CRC tumors from two additional reported datasets^35,36^ (Fig. 5e). Of note, we found almost no CD63 or Galectin-3 (encoded by *LGALS3*) expression on the neutrophils from PB of healthy donors, and we did not detect PM population from both healthy donors and cancer patients (Supplementary information, Fig. S15a). However, CD63 and Galectin-3 were expressed in neutrophils from the bone marrow of healthy donors and were predominantly enriched in MC & MM neutrophils, a finding consistent with our scRNA-seq results (Supplementary information, Fig. S15b). On the contrary, we confirmed that CD63 and Galectin-3 were highly expressed on human neutrophils from PB of GC patients, and MC & MM neutrophils represent the predominant CD63^+^ and Galectin-3^+^ populations in PB of cancer patients (Supplementary information, Fig. S15c-e). And MC & MM neutrophils, rather than eosinophils (CD11b^+^Siglec8^+^) or BD & SC neutrophils, are the dominant cell subsets to express CD63 and Galectin-3 in the PB of GC patients (Fig. 5f). Moreover, we accessed the scRNA-seq data of patient PDAC tumors (GSE155698) to identify genes that are highly expressed in CD63^+^LGALS3^+^ neutrophils and combined them with TCGA pan-cancer survival data to analyze the correlation between the CD63^+^LGALS3^+^ neutrophil-associated gene signature (quantified as a composite score) and patient prognosis (Supplementary information, Fig. S15f, g). The results demonstrated that elevated expression levels of the CD63^+^LGALS3^+^ neutrophil-associated gene signature is significant associated with worse patient survival (Supplementary information, Fig. S15h). We also collected tumor tissues from gastric cancer patients, and flow cytometry analysis showed that tumor-infiltrating neutrophils (SSC^hi^CD66b^+^), monocytes (CD11b^+^CD14^+^), macrophages (HLA-DR^+^CD163^+^), and DCs (HLA-DR^+^CD11c^+^) all expressed CD63 and Galectin-3 (Supplementary information, Fig. S15i). The expression levels of CD63 and Galectin-3 were highest in tumor-infiltrating neutrophils and monocytes, followed by macrophages, with DCs showing the lowest expression (Supplementary information, Fig. S15j). More importantly, we found that tumor-infiltrating CD63^+^ or Galectin-3^+^ cells consist of primarily MC & MM neutrophils (Fig. 5g).

### Flt3L transdifferentiates MC & MM-stage neutrophils into monocytic cells

Despite the discovery of surface markers for human MC & MM neutrophils, the shared expression of CD63 and Galectin-3 on eosinophils and macrophages prompted us to develop an alternative targeting strategy for cancer treatment. By analyzing scRNA-seq data from both tumors of cancer patients and BM of our NCG-*Gfi1*^-/-^ HIS mice, we noticed that human neutrophils and monocytes were clustered closely in t-SNE plots and have significantly correlated gene expression profile (Fig. 6a-c; Supplementary information, Fig. S16a, b), indicating similar gene expression patterns between these two populations. Considering that neutrophils at different developmental stages do not contain CD11b^+^CD14^+^ monocyte-like subpopulations (Supplementary information, Fig. S16c), we hypothesized that MC & MM neutrophils might respond to cytokines that drive monocytic differentiation. Considering the very low proportion of PM, we then sorted human MC & MM and BD & SC neutrophils from NCG-*Gfi1*^-/-^ HIS mice and cultured them with either G-CSF (reported as a key cytokine in granulopoiesis^37^) or Flt3L (reported as a key cytokine in monocytic differentiation^38^) ex vivo (Fig. 6d). Interestingly, Flt3L, instead of G-CSF, led to a significantly increased expression of monocytic markers (CD11b^+^CD14^+^) and decreased expression of neutrophil markers (SSC^hi^CD66b^+^) on cultured MC & MM neutrophils (Fig. 6e, f). Conversely, markers of BD & SC neutrophils remained unchanged after co-cultured with G-CSF or Flt3L (Fig. 6e, f). Moreover, considering that CD54⁻ neutrophils have been previously reported to have trans-differentiation potential^38^, we detected and observed that CD54⁻ neutrophils are primarily composed of MC & MM neutrophils in human BM (Supplementary information, Fig. S16d). We also sorted human BM neutrophils at each stage and separately evaluated their trans-differentiation potential under Flt3L treatment, and the results indicated that BD & SC neutrophils could not be transdifferentiated into monocytic cells by Flt3L (Fig. 6g, h; Supplementary information, Fig. S16e). We also conducted several analyses to compare monocytes, MC & MM neutrophils, and monocytes that are transdifferentiated from MC & MM neutrophils by Flt3L (Fig. 6i). Results of T cell suppression assay showed that the suppression of T cell proliferation was reversed after co-culture with Flt3L-educated MC & MM neutrophils (Fig. 6j), indicating that MCs & MMs lose their immunosuppressive properties after trans-differentiation into CD11b^+^CD14^+^ monocytic population. Giemsa staining results showed similar cell morphology (irregular cell membrane shape) between monocytes and monocytic cells (transdifferentiated from MC & MM neutrophils) (Fig. 6k). In addition, we found that CD11b^+^CD14^+^ monocytic population transdifferentiated from MCs & MMs lost the expression of immunosuppression-related gene markers such as *ARG1* and *LOX1* and acquired monocyte/macrophage marker *CD68* and *CD163* (Fig. 6l).

**Fig. 6.**
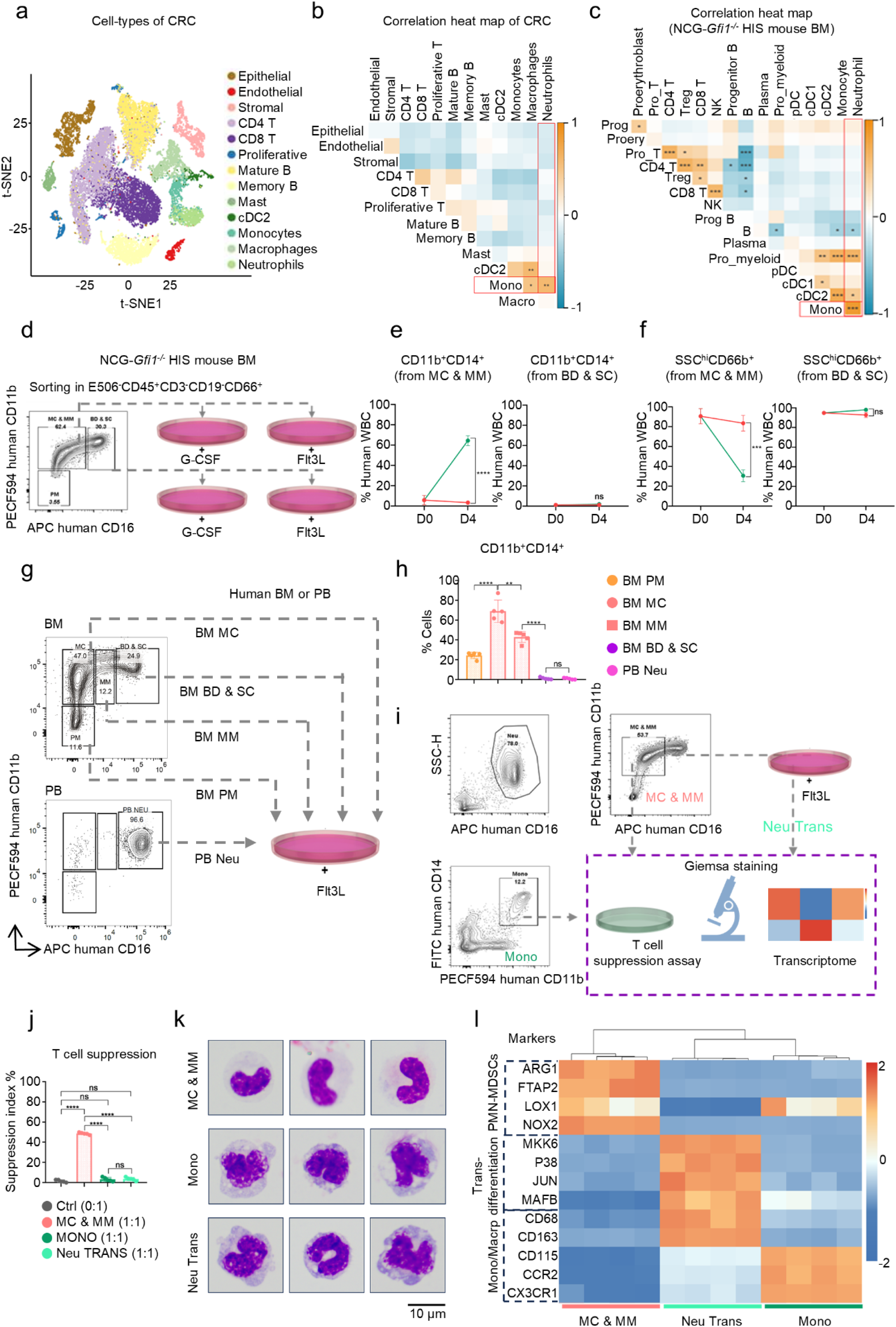
Flt3L transdifferentiates MC & MM-stage neutrophils into monocytic cells. **a** t-SNE clustering of cell types in the tumors of CRC patients^36^. **b** Heatmap depicting the correlation of cell types in the tumors of CRC patients. **c** Heatmap depicting the correlation of immune cell clusters from scRNA-seq data of NCG HIS and NCG-*Gfi1*^-/-^ HIS mice. **d** Schematic of MC & MM or BD & SC neutrophils sorted from BM of NCG-*Gfi1*^-/-^ HIS mice were cultured with G-CSF or Flt3L. **e** After culture for 4 days, proportions of CD11b^+^CD14^+^ cells among human WBCs were determined by flow cytometry (*n* = 3). **f** After culture for 4 days, proportions of SSC^hi^CD66b^+^ cells among human WBCs were determined by flow cytometry (*n* = 3). **g** Schematic of neutrophils at various stages sorted from BM and PB of non-tumor donors were cultured with Flt3L. **h** Proportions of CD11b^+^CD14^+^ cells in each group were determined by flow cytometry (*n* = 5). **i** Schematic of sorted monocytes, MC & MM, and CD11b^+^CD14^+^ population transdifferentiated from MC & MM for T cell suppression, giemsa staining and transcriptome detection. **j** Suppression index of human T cell proliferation co-cultured with monocytes, MC & MM, and CD11b^+^CD14^+^ population transdifferentiated from MC & MM (*n* = 5). **k** Giemsa staining of monocytes, MC & MM, and monocytes transdifferentiated from MC & MM by Flt3L. **l** Heatmap of PMN-MDSC, trans-differentiation, monocytes/macrophages and DC-related gene markers in monocytes, MC & MM, and CD11b^+^CD14^+^ population transdifferentiated from MC & MM (*n* = 4). All data are mean ± SD and were analyzed by two-tailed, unpaired Student’s t-test (\**P* < 0.05, \*\**P* < 0.01, \*\*\**P* < 0.001, \*\*\*\**P* < 0.0001).

### Flt3L-induced trans-differentiation of MC & MM-stage neutrophils control tumor growth in NCG-*Gfi1*^-/-^ HIS model

To further evaluate the potential of Flt3L mediated trans-differentiation as a MM & MC targeting immunotherapy in vivo, we overexpressed human G-CSF or Flt3L in HIS mice via hydrodynamic injection of cytokine encoding plasmids (which has been used by us as a stable method to overexpress proteins in mouse liver^39^) (Fig. 7a). Consistent with ex vivo culture, Flt3L overexpression led to an evident reduction of SSC^hi^ cell population (mostly MC & MM neutrophils) in the BM (Fig. 7b). Thus, both ex vivo co-culture and in vivo results support our hypothesis that human MC & MM neutrophils hold the potential to transdifferentiate into monocytic cell via Flt3L. Thus, all these ex vivo and in vivo results supported that human MC & MM neutrophils can be transdifferentiated into the monocytic population by Flt3L, which loses their immunosuppressive function.

**Fig. 7.**
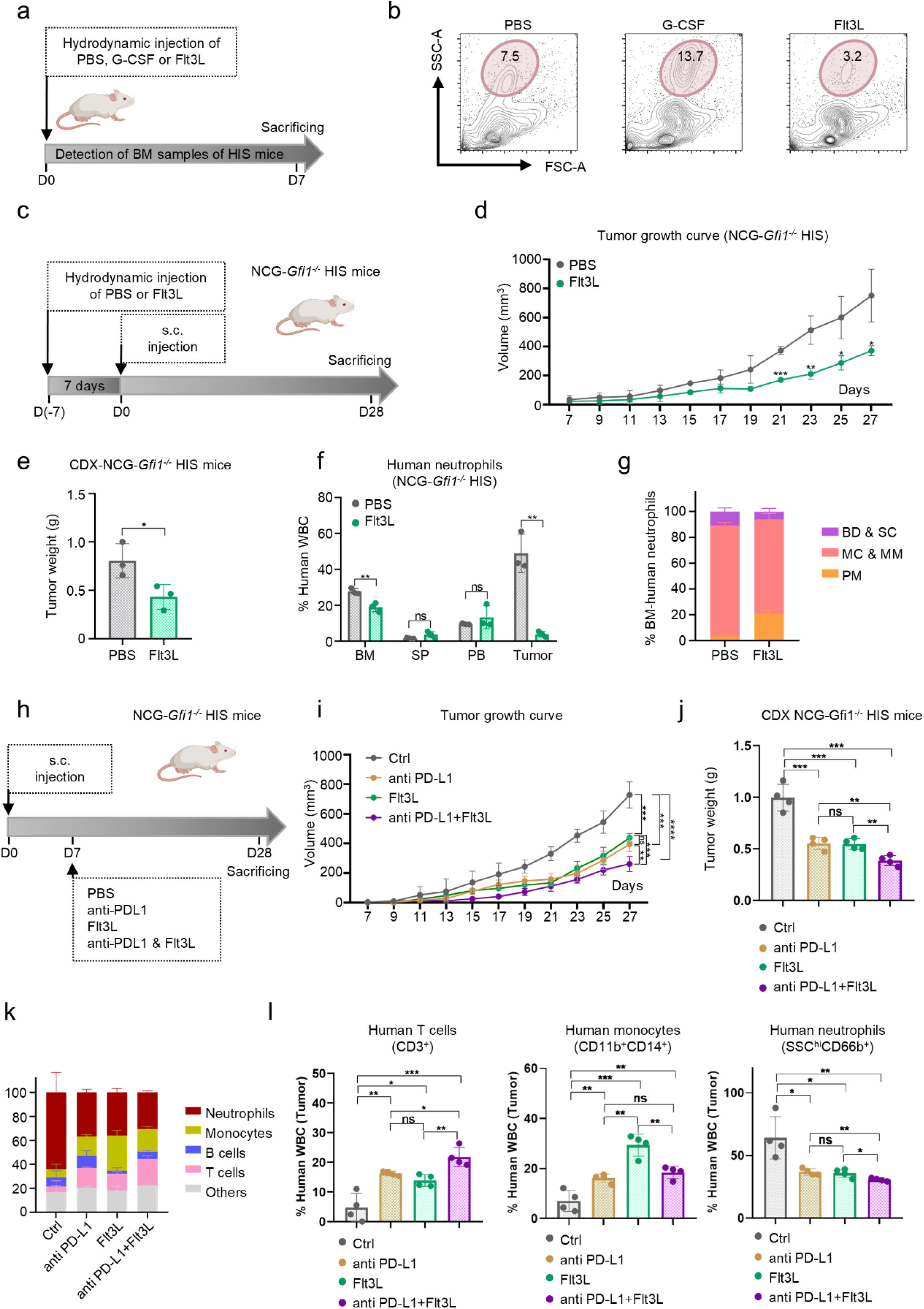
Flt3L trans-differentiation treatment controls tumor growth in NCG-*Gfi1*^-/-^ HIS model. **a** Schematic of NCG-*Gfi1*^-/-^ HIS mice were hydrodynamically i.v. injection of pLIVE-G-CSF or Flt3L plasmids. **b** representative flow cytometry plots of BM cells 7 days after the injection were shown. **c** As depicted in the schematic, A375 xenograft NCG-*Gfi1*^-/-^ HIS model was established, and pLIVE-Flt3L plasmid was injected i.v. hydrodynamically. **d** Tumor growth of NCG-*Gfi1*^-/-^ HIS mice with PBS or Flt3L treatment (PBS, *n* = 3; Flt3L, *n* = 3). **e** Tumor weight of NCG-Gfi1^-/-^ HIS mice with PBS or Flt3L treatment (PBS, *n* = 3; Flt3L, *n* = 3) **f** Proportions of neutrophils (CD11b^+^CD66b^+^) in the BM, SP, PB and tumors of NCG-*Gfi1*^-/-^HIS mice with or without Flt3L treatment (*n* = 3). **g** Proportions of human neutrophils at different developmental stages in the BM of NCG-*Gfi1*^-/-^HIS mice with or without Flt3L treatment (*n* = 3). **h** As depicted in the schematic, A375 xenograft NCG-*Gfi1*^-/-^ HIS model was established, and pLIVE-Flt3L plasmid was injected i.v. hydrodynamically while the anti-PD-L1 antibodies were injected i.p. (150 µg/dose, once every three days). **i** Tumor growth of NCG-*Gfi1*^-/-^ HIS mice with PBS, Flt3L and/or anti-PD-L1 treatment (*n* = 4). **j** Tumor weight of NCG-Gfi1^-/-^ HIS mice with PBS, Flt3L and/or anti-PD-L1 treatment (*n* = 4). **k** Proportion of different immune cell populations in the tumor of NCG-*Gfi1*^-/-^ HIS from each group (*n* = 4). **l** The proportion of human T cells (CD3^+^), monocytes (CD11b^+^CD14^+^) and neutrophils (SSC^hi^CD66b^+^) in the tumor of NCG-*Gfi1*^-/-^ HIS from each group (*n* = 4). All data are mean ± SD and were analyzed by two-tailed, unpaired Student’s t-test (\**P* < 0.05, \*\**P* < 0.01, \*\*\**P* < 0.001, \*\*\*\**P* < 0.0001).

We next evaluated the effect of Flt3L for tumor control in NCG-*Gfi1*^-/-^ HIS mice (Fig. 7c). We first pre-treated NCG-*Gfi1^-/-^* HIS mice with Flt3L hydrodynamic injection and one-week later s.c. inoculated A375 tumor cells. The tumor volume and tumor weight were significantly decreased in A375-inoculated NCG-*Gfi1*^-/-^ HIS mice after human Flt3L overexpression (Fig. 7d, e). Notably, the downregulation of BM neutrophils and the dramatical reduction of TINs were still observed after 35 days of Flt3L plasmid injection in NCG-*Gfi1*^-/-^ HIS mice (Fig. 7f). And the proportion of MC & MM neutrophils in the BM of NCG-*Gfi1*^-/-^ HIS mice was significantly reduced after Flt3L treatment (Fig. 7g), to the levels in the BM of non-tumor bearing NCG-*Gfi1*^-/-^ HIS mice (Fig. 4k). By contrast, we found that Flt3L overexpression failed to induce a tumor-control effect in A375-challenged NCG HIS mice (Supplementary information, Fig. S17), suggesting that Flt3L executes anti-tumor effect through altering the number and composition of neutrophils. After observing that Flt3L treatment exerts prophylactic efficacy in controlling tumor growth of the NCG-*Gif1^-/-^* HIS mice, we further evaluated the therapeutic potential of this trans-differentiation strategy. We found that Flt3L trans-differentiation strategy enhanced the anti-tumor effect of anti-PD-L1 therapy in A375-challenged NCG-*Gfi1*^-/-^ HIS mice (Fig. 7h-j). And we observed increased intratumoral T cell proportion, along with increased monocyte proportion and decreased neutrophil proportion (particularly a significant decline in MC & MM neutrophils in PB) in the tumor, BM, PB of mice receiving combination treatment (Fig. 7k, l; Supplementary information, Fig. S18). Together, our data demonstrate that trans-differentiation of MC & MM neutrophils into monocytic cells with Flt3L represents a potential strategy for enhanced cancer immunotherapy.

## DISCUSSION

In the current study, we separated the developmental stages of human neutrophils based on reported gating strategies^15,17,24^, and observed that MC & MM-stage neutrophils are the predominant cluster of neutrophils in human BM and TINs across 17 cancer types. Also, human MC & MM neutrophils exhibit robust immunosuppressive capabilities to inhibit T cell proliferation ex vivo and promote tumor growth in vivo. With the development of NCG-*Gfil*^-/-^ HIS mice, we found that BM MC & MM neutrophils drastically increased and potentially contributed to the circulating neutrophils and TINs. Moreover, by analyzing DEGs of human BM and tumor MC & MM neutrophils from scRNA-seq data, we identified CD63 and LGALS3 as surface markers of MC & MM neutrophils in pan-cancer patients. Finally, we noticed the plasticity of MC & MM neutrophils and discovered Flt3L as a potential approach to transdifferentiate MC & MM neutrophils into monocytic cells, which leads to tumor control effects in NCG-*Gfil*^-/-^ HIS mice (Fig. 8).

**Fig. 8.**
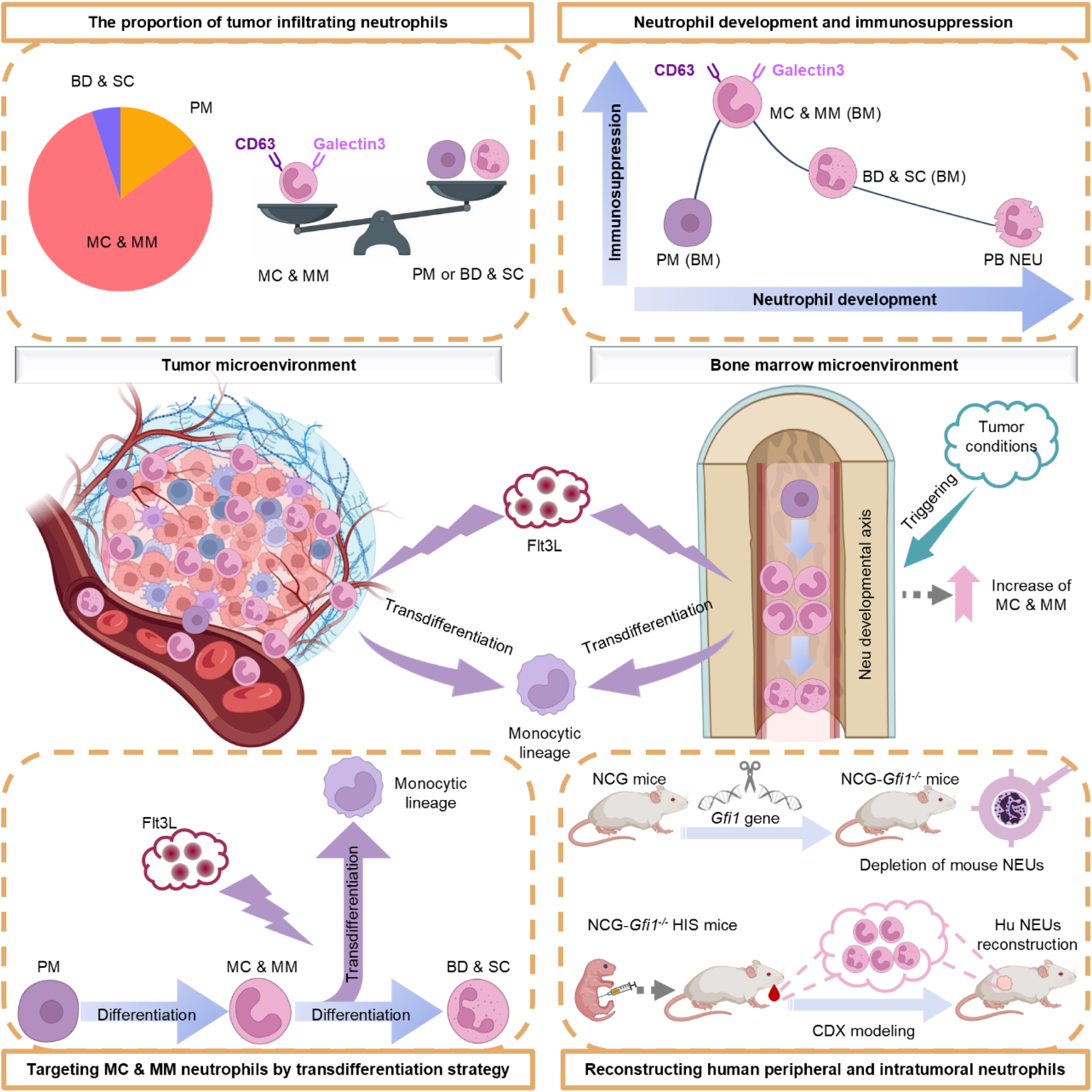
A schematic diagram illustrating dynamics of human MC & MM-stage neutrophils across physiological and tumor conditions. Human MC & MM-stage neutrophils, predominant in BM and constituting most TINs within the TIME across cancer types, exhibit immunosuppressive and tumor-promoting properties. Tumor-driven expansion of these cells occurs in BM and periphery of cancer patients and NCG-*Gfil^-/-^*HIS mice, where CD63 and Galectin-3 distinguish them from other neutrophils, while their transdifferentiation into monocytic cells via Flt3L retards tumor growth.

Current studies explore pro-tumor neutrophils through comprehensive analysis of the snapshots of TINs in one or several cancer models or patients, aiming to identify specific TIN clusters with pro-tumor properties^6–9,25,40–42^. For instance, a study by Zhang group identified CCL4^+^ and PDL1^+^ TANs, which recruit macrophages and suppress T cell cytotoxicity to induce immune suppression in the tumors of live cancer patients^8^. Another study by Ng group reported that in pancreatic cancer models, dcTRAIL-R1^+^ TINs show a prolonged lifespan and pro-angiogenic properties to fuel tumor growth^9^. However, considering the heterogeneity of neutrophils, the probability to identify a conserved protumor TIN subset would decrease with the increase of cancer types. In our study, instead of focusing on the clustering of TINs, we explored the function of neutrophils from a developmental perspective, and viewed TINs and BM neutrophils as a developmentally related continuum. We separated the developmental stages of human neutrophils into PM (CD66b^+^CD11b^-^), MC & MM (CD66b^+^CD11b^+^CD10^-^CD16^-/lo^), and BD & SC (CD66b^+^CD11b^+^CD10^lo/+^CD16^+^), and observed a clear predominance of MC & MM in human BM. Notably, MC & MM constitute the majority of TINs within the TIME of patients across 17 cancer types (including gastric, bladder, pancreatic, lung, breast, colorectal cancer, melanoma, etc.), which is also confirmed in the tumor-bearing NCG-*Gfil*^-/-^ HIS mouse model. Moreover, we explored the function of human MC & MM, and found that MC & MM neutrophils from either healthy donors or cancer patients exhibited robust immunosuppressive and tumor-promoting capabilities. Therefore, we propose to interrogate the function of neutrophil under the conserved developmental trajectory, and uncover the protumor role of human MC & MM neutrophils in a widespread of cancer types. Also, considering their dominance in BM and immunosuppressive properties, it is of value to explore the impact of MC & MM neutrophils in other pathological conditions such as autoimmune or inflammatory diseases.

Neutrophil studies rely on syngeneic mouse models as human surrogate ^12,13,43–47^, but the non-negligible species discrepancy may hinder the clinical translation of therapies targeting TINs into cancer patients^3,4,48^. Unlike human settings, our results showed a compositional difference that neutrophils in mouse BM and tumors are mostly in BD & SC stage, which is also consistent with previous observations of mouse TINs^49,50^. Our results further demonstrated that mouse neutrophils can acquire immunosuppressive and pro-tumorigenic properties within the tumor microenvironment, independent of their developmental stages. Thus, it is of necessity to develop animal models which recapitulate the clinical observations, especially for the composition of human TIN. Previous studies have reported HIS mouse models with G-CSF/G-CSFR gene modifications to support human neutrophil reconstruction, but the incomplete removal of mouse neutrophils leads to limited improvement of peripheral neutrophil levels ^22,23^. In this study, by inducing severe congenital neutropenia through *Gfi1* knockout in NCG mice and HIS construction^39,51^, we efficiently reconstitute peripheral and intratumoral human neutrophils, which precisely replicates the immune composition found in tumors of patients. Thus, NCG-*Gfi1*^-/-^ HIS model enables us to investigate human neutrophil composition and function in multiple tissues from one individual beyond tumor. We observed further increase of human MC & MM neutrophils in BM of CDX NCG-*Gfi1*^-/-^ HIS mice, and postulated that tumor cells remotely expand and recruit MC & MM neutrophils in the BM, possibly via secreted proteins.

As BM serves as the potential source of MC & MM neutrophils in the tumor microenvironment (TME), it is necessary to target both BM and intratumoral neutrophils for the development of optimal diagnostic and therapeutic strategies. In this study, *CD63* and *LGALS3* were identified from overlapped DEGs of MC & MM neutrophils from scRNA-seq datasets of HIS BM, human BM and 17 types of tumors. High *CD63* and *LGALS3* expressions in tumor are also associated with worse survival outcomes in patients from 32 cancer types in TCGA database analyzed by GEPIA tool. Tetraspanin CD63 is the main component of extracellular vesicles^52^, whose expression on tumors has been reported to have regulatory role in cancer malignancy^53–56^. LAGLS3 (human gene encoding Galectin-3), which belongs to the galectin family, has also been reported to promote treatment resistance in both solid and hematologic cancers^57–60^. Additional scRNA-seq and flow cytometry data from human PB further demonstrated relatively restricted expression of these two markers on MC & MM neutrophils. Therefore, CD63 and LGALS3 hold promise as diagnostic biomarkers across cancer types and show advantages over known markers of human MC & MM neutrophils (CD10, CD16, etc.)^3,61–64^. Although MC & MM-neutrophils are the main composition of CD63⁺ or Galectin-3⁺ cell populations, the co-expression of these markers in eosinophils and monocytes may compromises their potential to be therapeutic targets specific for MC & MM-neutrophils. This limitation highlights the value our Flt3L-driven trans-differentiation therapy as more specific strategy targeting MC & MM. Also, analyzing the association of CD63 and LGALS3 expression with patient survival using GEPIA tool has some limitation, as this method does not consider the possible bias caused by treatment or other confounders (sex, age, etc.). As such, the therapeutic potential and safety profile of these two markers need to be investigated in the future studies.

Recent studies have shed light on the importance of anti-tumorigenic neutrophil for the successful T cell-based cancer immunotherapies^12–14,65^, provoking the thoughts of transforming the protumor neutrophils into antitumor neutrophils^12,13,66^. In our study, we observed unexpected similarity between human neutrophils and monocytes in both BM and patient tumor samples, and boldly hypothesized the possibility of trans-differentiation from MC & MM neutrophils to monocytic cells. Notably, we found Flt3L induced trans-differentiation of MC & MM neutrophils (SSC^hi^CD66b^+^CD14^-^) into monocytic lineages (SSC^low^CD66b^-^CD14^+^), but did not alter the state of BD & SC neutrophils. Coincidentally, a study by Mastio et al. reported the presence of monocyte-like precursors of granulocytes in mouse BM, which expanded and differentiated into polymorphonuclear myeloid-derived suppressor cells under tumor-bearing conditions^67^. And Strobl et al. suggested the feasibility to in vitro convert human band-stage neutrophils into monocytic cells via MKK6-p38MAPK inflammatory signaling^38^. Thus, trans-differentiation between MC & MM neutrophils and monocytic cells could be bidirectional and conserved between mouse and human. Meanwhile, considering the abundance of neutrophils in human PB and their short lifespan^1,68^, we postulate that human hematopoiesis might have evolved to accumulate a large number of MC & MM neutrophils as a reservoir in the BM. For this reason, compared to mice, cancer patient could be more influenced by MC & MM neutrophils, particularly those from the BM. In this study, we found that overexpression of Flt3L efficiently reduced TIN infiltration and the proportion of MC & MM neutrophils in BM, which brought tumor-control effects in NCG-*Gfi1*^-/-^ HIS mice. Therefore, in addition to converting TIN into anti-tumor neutrophils, we propose an alternative strategy with Flt3L to specifically transdifferentiate both BM and intratumoral MC & MM neutrophils into monocytic cells for the treatment of cancer patients. Although, there is one study reporting the role of Flt3L on promoting the differentiation of promonocytes into mature monocytes^38^, to our knowledge, we are the first to demonstrate that Flt3L can also induce the trans-differentiation of MC & MM neutrophils into monocytes, thereby reversing the neutrophil-caused immunosuppression in the TME. Notably, based on our bulk RNA-seq analyses, the PD-L1 (encoded by *CD274*) expression in human neutrophils at various stages are very low, indicating that neutrophils are not the primary source of PD-L1 expression. However, tumor cells and other myeloid cell populations such as macrophage are reported to have high PD-L1 expression, which causes immune evasion^69,70^. Therefore, we also evaluated the treatment effect of Flt3L trans-differentiation strategy combined with PD-L1 blockade in the mouse tumor model, and found enhanced tumor-control effect compared to single treatment. Overall, our preclinical results using NCG-*Gfi1^-/-^* HIS mice suggest that Flt3L trans-differentiation strategy is a promising new treatment approach targeting neutrophil developmental stage for cancer control, as previous neutrophil-targeting therapies ignore the impact of neutrophil developmental stage in the TME.

In summary, our work identifies human MC & MM-stage neutrophils, which are immunosuppressive and predominate among TINs across tumor types, as the key contributor to cancer progression. We also highlight the broad prospects for future studies to analyze and target the developmental stages of neutrophils for cancer prevention, monitoring, and treatment.

## MATERIALS AND METHODS

### Mice

All mice were housed in specific pathogen-free conditions in animal facilities certified by an Institutional Animal Care and Use Committee (IACUC) at the Model Animal Research Center in Nanjing University (AP# LY-01). We obtained several mouse strains from Gempharmatech, NOD Prkdc^scid^ Il2Rγc^-/-^ (NCG, Cat No. T001475) and C57BL/6JGpt (Cat No. N000013). The NCG-*Gfi1*^-/-^ strain was originally constructed by our team. In brief, we performed mouse *Gfi1* gene knockout on the basis of NOD mice. Three sgRNAs were designed for the sixth zinc finger domain sequence of the mouse *Gfi1* gene: sgRNA1 (AGCTTTGACTGTAAGATCTGTTGG), sgRNA2 (TGCTCATTCACTCGGACACCCGG), and sgRNA3 (GGGCCGGGATAGGGACAGTCATG). Three sgRNAs and Cas9 mRNA were co injected into mouse fertilized egg cells to obtain *Gfi1*^-/-^ fertilized egg cells, and transfer these *Gfi1*^-/-^ fertilized egg cells to pseudo pregnant female mice to obtain F0 *Gfi1*^-/-^ mice. Afterwards, we co bred NOD-*Gfi1*^-/-^ mice with NCG mice to gradually obtain NCG-*Gfi1*^-/-^ mice. HIS mice were generated in NCG or NCG-*Gfi1*^-/-^ hosts as previously described ^39^. Intrahepatic injection of 5×10^4^ CD34^+^CD38^-^ human fetal liver hematopoietic stem cells (HSCs) into newborn pups (4∼6 days old) was performed. After several weeks, 50 µL of blood was drawn from each mouse to check human cell reconstitution. Subsequent experiments used NCG or NCG-*Gfi1*^-/-^ HIS mice with human CD45^+^ cells in the blood larger than 1×10^5^/mL.

### Xenograft and syngeneic cancer mouse models

For the establishment of xenograft human melanoma mouse model, NCG HIS or NCG-*Gfi1*^-/-^ HIS mice were injected subcutaneously with 100 µL DPBS containing 5×10^6^ A375 cells. For the establishment of syngeneic GC mouse model, MFC cells (1 × 10^6^ per animal) were injected subcutaneously (s.c.) into 6–8-week-old sex-matched 615-line mice to form solid tumors. Tumor volume was evaluated using the equation volume = (xy^2^)/2, where x and y are the lengths of the major and minor axes, respectively.

### Neutrophil transfer in vivo

We conducted human BM neutrophil transfusion experiments on the 13th and 21st days after CDX modeling. We sorted MC & MM (CD66b^+^CD11b^-^CD10^-^CD16^-/lo^) and BD & SC (CD66b^+^CD11b^-^CD10^lo/+^CD16^+^) neutrophils from human BM by flow cytometry and injected these cells into CDX-NCG HIS mice through the tail vein. The injection dose for each mouse is 2.5×10^6^ MC & MM or BD & SC neutrophils.

### Plasmids and reagents

Human plasmids were obtained from Origene company, and their product information is Flt3L (Cat#: RC222242) and G-CSF (Cat#: RC217237). Reagents and cytokines for cell culture and tissue digestion were Flt3L (Sinobiological, Cat#:10315-H07B), G-CSF (Sinobiological, Cat#:10007-H01H), DMEM (Biological Industries, Cat#: 06-1055-57-1ACS), RPMI 1640 (Biological Industries, Cat#: 01-100-1ACS), fetal bovine serum (FBS) (Gibco, Cat#: 10099141), penicillin/streptomycin (P/S) (Gibco, Cat#: 15140122), Dulbecco’s phosphate buffered saline (Biological Industries) (Bioind, Cat#: 02-023-1ACS), collagenase mix (SIGMA, Cat#: 9001-12-1), DNase I (SIGMA, Cat#: DN25), Ficoll (GE Healthcare, Cat#: 17544203), Red Blood Cell Lysing Buffer Hybri-Max (SIGMA, Cat#: R7757-100ML), Atezolizumab (anti PD-L1, MedChemExpress, Cat#: HY-P9904) and Lysing Solution 10× Concentrate (BD, Cat#: 349202).

### Human samples

The BM samples of the control group (non-tumor) were collected from arthritis patients aged 50∼70 years old, who were also in the same age range of tumor patients. The HSCs we used were sorted from fetal livers. Both tumor-related samples (Ethics Protocol #2021-324-03, #2009022) and fetal liver samples (Ethics Protocol #2021-488-01/02) were obtained from Nanjing Drum Tower Hospital. The study protocol was reviewed and approved by the Ethics Committee of Nanjing Drum Tower Hospital in accordance with the Declaration of Helsinki. Because the patients in the public databases could not be identified, the analysis and reporting of the data in our study were exempt from review by the Ethics Committee of Nanjing Drum Tower Hospital. The requirement for written informed consent to participate was waived. The sampling process was certified by the ethics committee and all donors signed informed consent. We prepared a sterile 10 cm petri dish and added culture medium (DMEM with 1% P/S and 2% FBS). We cut off the gallbladder and cut the fetal liver into small pieces of 1∼2 mm in a petri dish, transferred them to a centrifuge tube, and added the preheated digestion solution (DMEM with DNase I (5 mg/100 mL), collagenase (50 mg/100 mL), 1% P/S and 2% FBS) maintained at 37 °C for 30 min. Then, the samples were filtered through a 70 µm sieve, the filtrate was collected and centrifuged (500 g, 4 °C, 10 min), and the supernatant was discarded. We pipetted the pelleted cells with culture medium and enriched the leukocytes with Ficoll. Human CD34^+^ cells were purified by a Direct CD34 Progenitor Cell Isolation Kit and subsequently phenotyped for CD38 expression. After paying attention to the aseptic operation, all steps were carried out in a biosafety cabinet. We collected fresh human BM cells, centrifuged them (500 g, 4 °C, 5min), discarded the supernatant, and resuspended them in FACS buffer to remove the oil layer in BM cells. Then, after lysing red blood cells, we manipulated the leukocytes in BM cells for flow cytometry and cell sorting. Peripheral blood of healthy donors or gastric cancer patients was collected through an anticoagulation tube containing heparin and EDTA. Since the half-life of neutrophils in peripheral blood is very short, we should operate on ice and complete the experiment within 12 h. We contacted the surgical department and asked them to take human gastric, pancreatic and bladder cancer and melanoma tissues.

### Tissues processing for flow cytometric analysis and sorting

Staining for flow cytometry and cell sorting was performed on ice in FACS buffer (DPBS with 2% FBS, 1% P/S, and 2 mM EDTA). Human gastric tumors or CDX tumors were minced and treated with RPMI 1640 with collagenase (0.75 mg/mL) and DNase I (50 μg/mL), with rigorous agitation (150 rpm, 37 °C, 1 h). Spleen and BM cells can be directly triturated to obtain a single cell suspension. Tissue cells to be tested are first stained with e506 for 15 min to label live or dead cells. Then, the cells were stained with antibody mix for 35 min. Peripheral blood should be lysed red blood cells after antibody staining. Before testing on the flow cytometer, the cell suspension was passed through a 70 µm sieve to prevent clogging. Flow cytometric analysis was carried out by an LSRFortessa (BD Biosciences) and Novocyte Penteon (Agilent), and cell sorting was performed on a FACS AriaIII (BD Biosciences). Flow Cytometry Standard (FCS) files were analyzed by FlowJo. Antibody product information is summarized in Supplementary information, Table S4.

### T cell suppression assay

Human T cells or mouse T cells were labeled with 1 µM CSFE (Invitrogen, Cat #: 65-0850), and labeled T cells were placed into each well in a U-bottom 96-well plate and mixed with several types of myeloid cells sorted from the bone marrow of C57BL/6J mice, HIS mice or humans (the ratio is labeled in each assay among figures). Human T-cell cultures were stimulated for 96 h with anti-CD3/CD28 Dynabeads (Gibco, Cat #: 11131D) at a cell to bead ratio of four to one. Mouse T cells were stimulated with antibodies against CD3e (5 ng/wells, eBioscience, Cat#: 14-0031) and CD28 (2 µg/mL, eBioscience, Cat#: 16-0281). The suppression index is calculated as 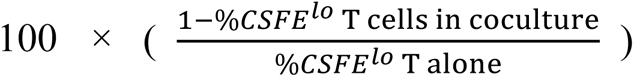.

### RNA extraction and qPCR

We used TRIzol (Invitrogen, Cat #: 15596026) to lyse the cells to release the RNA and then extracted and purified the RNA with chloroform, isopropanol and 75% ethanol. The concentration of RNA was measured by Nanodrop. Quantitative RT–PCR was performed using the ABI Prism Step-One system with Reagent Kit from Yeasen Biotechnology (Cat#: 11141-B,11141-C and 11201ES03). Primer sequences are summarized in Supplementary information, Table S5.

### Bone marrow dissociation and preparation

The mice were sacrificed, the femur and tibia were dissected, and the muscles and periosteum on the outer surface of the femur and tibia were removed. We triturated directly to obtain BM single cell suspension. The single-cell suspension was then centrifuged at 500 g for 5 min at 4 ℃. Next, we resuspended the cells in 1 mL of red blood lysis buffer for 15 min in room temperature to remove the red blood cells. Then we added 10 mL of FACS buffer to terminate red blood lysis, and the solution was centrifuged at 500 g for 5 min at 4 ℃ and resuspended in FACS buffer. Then, flow cytometry analysis and sorting were performed according to the above method.

### Bulk RNA sequencing

Cells were sorted using FACSAria III cell sorter, and RNA was extracted according to the above method. The following antibodies were used: BV510-conjugated anti-mouse CD45, BV605-conjugated anti-human CD45, FITC-conjugated anti-human CD14, APC-conjugated anti-human CD66b, PECF594-conjugated anti-human CD11b, APC-Cy7-conjugated anti-human CD16 and Fixable Viability Dye eFluor 506. The libraries and sequencing were conducted by Novogene (Beijing, China) and ANNOROAD (Beijing, China). Libraries were constructed using NEBNext Ultra RNA Library Prep Kit for Illumina, and samples were sequenced on the Illumina NovaSeq 6000 (Illumina, USA) sequencer with 150-bp paired-end reads.

### Bulk RNA sequencing analyses

The sequencing data were filtered with fastp (v0.23.4) to remove reads with sequencing adaptors^71^, low-quality reads (reads with Q_phred_ < = 20 bases accounted for more than 50 % of the total reads length), and reads with ‘N’ bases (where ‘N’ represents undetermined base information). Clean reads were stored in fastq format. The clean reads were aligned to the reference genome of Homo sapiens (UCSC, hg38) using hisat2 (v2.2.1)^72^. Aligned reads were converted and then sorted by samtools (v1.6)^73^. Gene counts were obtained using featureCounts (v2.0.2) on GENCODE v33 annotations^74^. Differential expression analysis were carried out using DESeq2 (v1.20.0) with default parameters. Genes with fold change > 1.5 and Benjamini-Hochberg false discovery rate < 0.05 were considered significantly differentially expressed^75^. Transcripts per million (TPM) was selected as the gene expression value for RNA sequencing, and then gene expression TPMs were log2-transformed. Principal component analysis (PCA) of Monocytes/Neu Pro/Immature Neu/Mature Neu samples were performed by using PCA tools in Hiplot Pro (https://hiplot.com.cn/), a comprehensive web service for biomedical data analysis and visualization.

### Changes in gene expression during neutrophil development

In order to obtain the markers of different developmental stages of neutrophils, we intersected the significantly high expression genes of neutrophils at various stages obtained from our RNA-seq data of PM, MC, MM, and BS & SC Neu samples with the gene sets obtained from published data^28,29^. Specifically, significantly high expression genes identified from our bulk RNA-seq analyses were intersected with gene groups (g4-g10) and gene clusters (r1-r7) obtained from Calzetti F et al. and Grassi L et al ^28,29^. The Venn diagram was drawn using the online website (https://jvenn.toulouse.inrae.fr/app/index.html)^76^. Finally, we obtained 671 genes that were significantly highly expressed in PM, MC, MM, and BS & SC Neu, respectively.

### Single cell collection and sequencing

Based on FACS sorting, human CD45^+^ living cells from 8 HIS mice were isolated for sequencing. The following antibodies were used: BV510-conjugated anti-mouse CD45, BV605-conjugated anti-human CD45 and Fixable Viability Dye eFluor 506. Libraries were constructed using 10X Genomics 3’ CellPlex Multiplexing Solution according to the manufacturer’s instructions. Each sample was uniquely labeled using custom antibody-based hashtag oligos (HTOs). And then labeled cells were pooled prior to loading onto a 10X Genomics chip. Gene expression (GEX) library and HTOs library were sequenced on the Illumina NovaSeq 6000 to depth of 30,000 reads and 5,000 reads.

### scRNA-seq data access

The single-cell gene expression data for PDAC, CRC and human BM (BM1) were collected from the Gene Expression Omnibus database (10X Genomics data, PDAC: GSE155698; CRC: GSE188711; BM1: GSE130756)^35,36,77^. We also downloaded another scRNA-seq h5ad file of human BM (BD Rhapsody data, BM2) from the website https://usegalaxy.eu/published/history?id=008000b0c3d8bb03^78^. In addition, a Seurat object containing neutrophils from 17 types of cancer was downloaded from the website http://www.pancancer.cn/neu/^14^.

### Single-cell RNA-seq data processing

The Cell Ranger toolkit (v6.1.1, 10X Genomics) was used for read processing, alignment with the hg38 transcriptome, and unique molecular identifier (UMI) counting/collapsing. Main downstream analyses were run with Seurat (v4.1.1) package in R (v4.1.3)^79,80^. In all the datasets, genes detected in fewer than 3 cells were filtered out prior to further analyses. For HIS mice BM data, HTOs were used for demultiplexing, filtering and clustering, and singlets were extracted from the HTOs library for downstream analyses. The initial filtration of HIS mice BM and human BM (BM1 and BM2) cells used the “percent.mt < 10” and “nFeature_RNA > 180” thresholds. For tumor scRNA-seq data, cells with fewer than 180 genes detected or with percent.mt higher than 30% were filtered. Then the UMI count matrix was standardized and centralized, the highly variable genes were calculated, dimension reduction and the clustering were performed. DoubletFinder (v2.0.3) R package was used to detect doublets in each sample^81^. Finally, 21625 cells in HIS mice BM, 16791 cells in human BM1, 11549 cells in human BM2, 76319 cells in 17 types of cancer, 43009 cells in PDAC and 26905 cells in CRC were retained for downstream analysis. Since the samples of human BM1 and 17 types of cacer come from different datasets and patient individuals, this may introduce technical and biological batch effects in downstream analysis. Therefore, based on PCA, the RunHarmony function of harmony (v1.0) R package was used to correct the batch effect^82^. For PDAC and CRC, since they contained different patient individuals from the same dataset, we used the anchor-based CCA integration method to obtain better integration results.

### Cell type annotation

All clusters were annotated according to the distinct expression patterns of the following reported cell markers^83,84^: Hematopoietic stem/progenitor cells (*SPINK2/CD34*), Erythrocytes and precursors (*TFRC/ALAS2/GYPA/BPGM/HBB/HBG1/HBG2/HBA1/HBA2*), Epithelial cells (*EPCAM*), Endothelial cells (*PECAM1/CLDN5*), Stromal cell (*COL1A2*), Immune cells (*PTPRC*), T cells (*CD3D/CD3E/CD3G*), NK cells (*NKG7/KLRD1*), B cells (*CD79A/CD79B/CD19*), Plasma cells (*MZB1/SSR4*), Classic dendritic cells (*HLA-DQA1/HLA-DQB1/HLA-DMA/HLA-DRB5/CST3/LYZ/HLA-DMB/FCER1A*), Plasmacytoid dendritic cells (*IRF8/IRF7/LILRA4/TCF4/ITM2C/MZB1/UGCG/SERPINF1*), Monocytes and macrophages (*CD68/CD163/CD14/CSF1R/FCGR2A*), Mast cells (*HPGD/TPSAB1/TPSB2/MS4A2/GATA2*) and neutrophils (*S100A8/S100A9/S100A12/CSF3R/FCGR3B/CXCR2/G0S2*). Neutrophils were identified for downstream analysis and performed the same processing steps and further clustering. We used markers of different developmental stages of neutrophils obtained from the RNA-seq dataset and previous reports^28,29^, such as *ELANE/MPO/PRTN3/CTSG/CTSC/PSS57/AZU1/SLPI/BPI* (Promyelocyte, PM), *CAMP/LTF/LCN2/MMP8/CAECAM8/CEBPE* (Myelocyte, MC), *MMP9/ORM1/PADI4/ S100A12/IFIT1/CD177* (Metamyelocyte, MM), *CXCR2/CSF3R/FCGR3B/MME/SELL* (Band & Segmented, BD&SC), to annotate the main neutrophil clusters.

### Score according to cell markers

We performed bulk RNA-seq on monocytes, PM, MC & MM, and BD & SC neutrophils sorted from human BM (as described above), with the aim to establish gene sets specific for human neutrophils at various developmental stages. Significantly high expression genes (fold change > 1.5 and Benjamini-Hochberg false discovery rate < 0.05) identified from our bulk RNA-seq analyses were intersected with gene groups (g4-g10) and gene clusters (r1-r7) obtained from Calzetti et al. and Grassi et al to ensure the robustness of the marker genes^28,29^. Through this process, we identified 671 genes that were significantly highly expressed in Neu Pro, Immature Neu, and Mature Neu, respectively (Supplementary information, Table S1, Score_genes.xlsx). To evaluate the gene expression patterns of neutrophil subsets in the BM of human, NCG HIS, NCG-*Gfi1^-/-^*HIS mice and TIN subsets of 17 types of cancer, genes with significantly high expression of PM, MC&MM, BD&SCin bulk RNA-seq data were taken as reference. The ref. score was calculated using “AddModuleScore” function in Seurat.

### Identification of differentially expressed genes (DEGs)

Differentially expressed genes (DEGs) expressed between PM, MC, MM and BD&SC neutrophil clusters were identified using the Wilcoxon method implemented by “FindMarkers” or “FindAllMarkers” functions in Seurat. Genes with log2 fold change more than 0.25 and adjusted p value less than 0.05 were considered as DEGs (Supplementary information, Table S2, DEGs_overlap.xlsx). We intersected the significantly highly expressed genes of MC and MM neutrophil clusters in *Gfi1*^-/-^ HIS mice BM, NCG HIS mice BM, human BM and human tumors, and obtained 8 genes (*FCER1G*, *CD63*, *LGALS3*, *CYBB*, *CTSD*, *FGR*, *GRN*, *FLNA*).

### Pearson correlation analysis

In order to reveal the similarity between neutrophil and monocyte clusters in HIS mice BM, PDAC and CRC, their Pearson correlation coefficients were calculated in the CCA integrated PCA space. Each cell cluster was represented by the centroid of the first 20 principal components of all cells belonging to that cluster. Then Pearson correlation coefficients were calculated using these centroids by R package psych (v2.4.3). The correlation coefficients matrix heatmap was plotted using the R package corrplot (v0.92).

### Pseudo-time trajectory analysis

In order to infer developmental dynamics from scRNAseq data of BM, we first sampled cells. If the number of cells was more than 20,000, 20,000 cells were randomly selected. If the number of cells is less than 20,000 cells, there is no need to sample. Pseudotime trajectories were inferred by scTour^85^, a deep learning-based framework for single-cell trajectory analysis. A neural network model was trained to learn the underlying dynamics of the data using the train_model function. Trajectory plots were visualized using scTour’s built-in plotting functions. To validate and refine the pseudotime trajectories inferred by scTour, we employed Monocle3^86^, where cells were ordered along a pseudotemporal axis based on their transcriptional similarity. A trajectory graph was constructed using the learn_graph function, and the root of the trajectory was manually selected by identifying the PM clusters. To visualize cell distribution and gene expression trends along pseudotime, we used ggplot2 package. Cell density along pseudotime was plotted using geom_density_ridges function from the ggridges package, highlighting the distribution of cells across pseudotime for different clusters or cell types. Gene expression trends along pseudotime were visualized using natural cubic splines (splines::ns()) to fit smoothed curves. A sequence of pseudotime values was generated, and the predict function was used to calculate fitted values and their 95% confidence intervals (lower and upper bounds). These values were plotted as smoothed curves with shaded confidence intervals, representing the uncertainty in the gene expression trends.

### Mortality risk and survival analysis

The survival analysis of 32 types of cancer was conducted by using the GEPIA2 (http://gepia2.cancer-pku.cn/#index). In GEPIA, the “0” time point typically represents the time of initial diagnosis, rather than the first treatment, surgery, or remission. GEPIA utilizes publicly available data from The Cancer Genome Atlas (TCGA) and the Genotype-Tissue Expression (GTEx) project, where survival analysis is based on the time from diagnosis to an event (e.g., death or disease progression). GEPIA’s survival analysis does not explicitly adjust for confounders such as sex, age, or other clinical variables. Specific gene expression levels were subjected to univariate Cox regression analysis. The prognostic impacts of gene expression level were displayed by the heatmap of mortality risk. We defined the top 50% and bottom 50% of patients with specific gene or signature expression in the TCGA/GTEx cohort as overexpression and low expression groups. Kapla-Meier survival analysis was further used to identify the survival difference between these two groups. The log-rank test was used to analyze survival rate.

### Data analysis and statistics

Data for all experiments were analyzed with Prism 8 software (GraphPad). Biological replicates and graph presentation in each figure are shown as the mean ± SD mentioned in the figure legends. Statistical significance was assessed by paired or unpaired, Student’s t test (\**P* < 0.05, \*\**P* < 0.01, \*\*\**P* < 0.001, \*\*\*\**P* < 0.0001) to compare the means of two groups. For sequencing data, R (v4.1.3) was used for statistical analysis. Wilcoxon rank-sum tests were used for continuous variables. Adjusted P-value was obtained through Bonferroni’s or Benjamini-Hochberg’s correction. Adjusted *P* < 0.05 was considered to be statistically significant.

## DATA AVAILABILITY

This study did not generate new materials. The raw sequence data reported in this paper have been deposited in the Genome Sequence Archive in National Genomics Data Center, China National Center for Bioinformation / Beijing Institute of Genomics, Chinese Academy of Sciences (GSA: HRA011772) that are publicly accessible at https://ngdc.cncb.ac.cn/gsa. The scRNA-seq and bulk RNA-seq data are also available from the corresponding author on request.

## ACKNOWLEDGMENTS

This work was funded by grants from the National Science and Technology Major Program (2023ZD0500400), the Fundamental Research Funds for the Central Universities (2024300408, XJ2024003602), the National Natural Science Foundation of China (32471000, U24A20378, 82373263, 82403835), the National Key Research and Development Program of China (2023YFC2506400), the Jiangsu Provincial Science and Technology Plan Special Fund (BK20232018) and the Funding of the Major Program of Shenzhen Bay Laboratory.

## AUTHOR CONTRIBUTIONS

Y.Li. and W.L. designed the study. Y.Li., J.W., T.M., and Y.Liang. supervised the study and revised the manuscript. W.L. performed the experiments and analyzed the data. with the assistance from T.S., C.L., K.C., Z.Z., Y.Luo., D.H., H.W., Shaorui.L., Y.W., Shuang.L., H.S., J.L., Y.Liu, D.S., S.D., H.X., L.L., J.X. and Jun X. T.S. and W.L. drafted the manuscript. The clinical samples were provided by S.D., Y.Luo. and Y.Liu. The bioinformatic analyses were conducted by C.L. NCG-*Gfi1^-/-^* mice were created by Y.Li. W.L. L.L., J.X., Jun X., and Y.Liang. All authors read and approved the manuscript.

---

**Supplementary information, Fig. S1.**
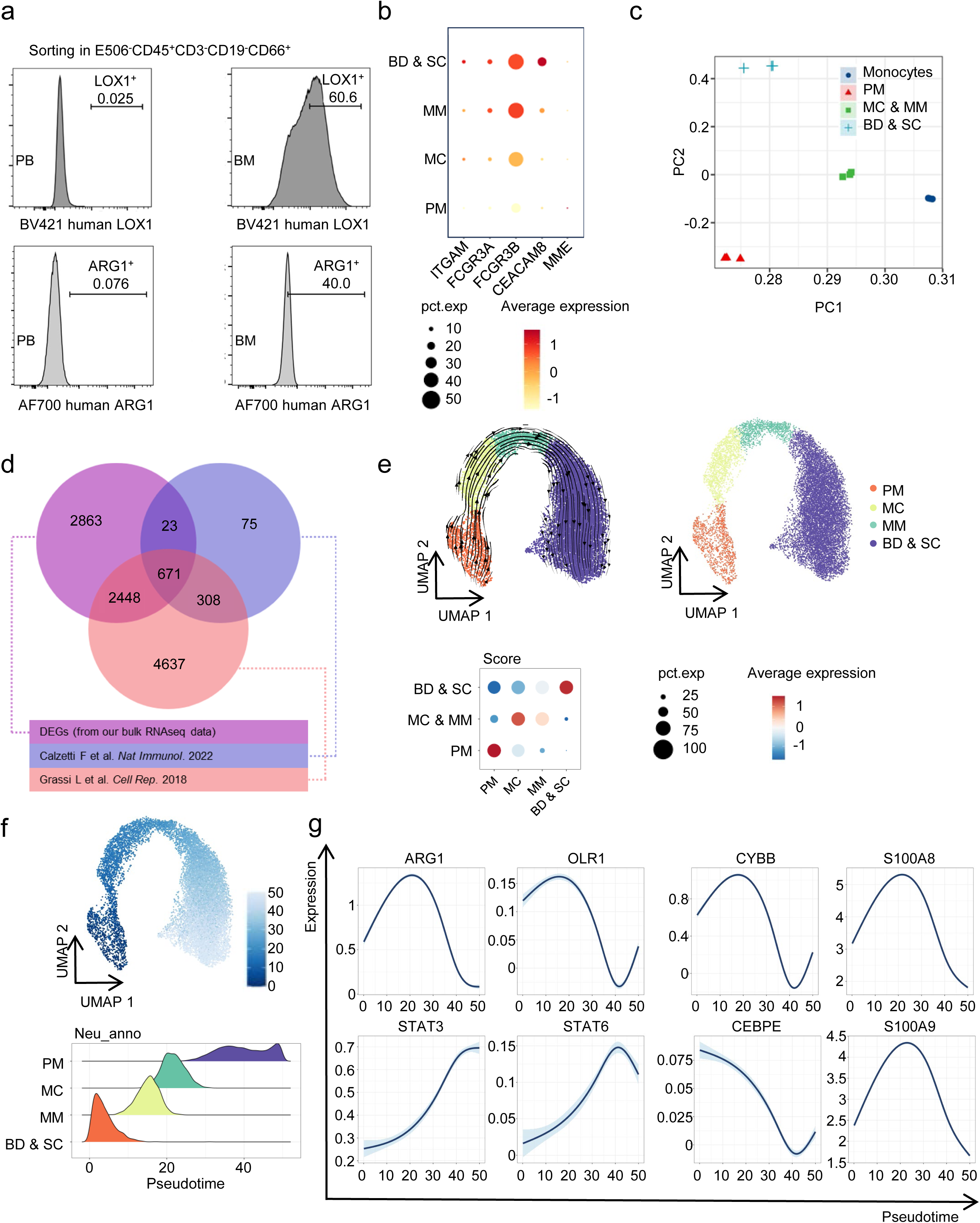
Flow cytometry and scRNA-seq of human BM neutrophils, related to Fig. 1. **a** Flow cytometry gating strategies for LOX1^+^ & ARG1^+^ neutrophils. Pseudotime **b** Dotplot illustrating the expression analysis or sorting marker levels of neutrophils at various stages within single-cell RNAseq dataset. **c** Bulk RNA-seq of different myeloid cell populations (monocytes: Lin^-^CD66b^-^CD11b^+^CD14^+^, PM: Lin^-^CD66b^+^CD11b^-^, MC & MM: Lin^-^ CD66b^+^CD11b^+^CD10^-^CD16^-/lo^, BD & SC: Lin^-^CD66b^+^CD11b^+^CD10^lo/+^CD16^+^) sorted from BM of non-tumor donors (*n* = 3) was performed, and PCA analysis was shown. **d** Venn diagram depicting DEGs of human neutrophils from us and two reported datasets ^28,29^. **e** We analyzed the scRNA-seq dataset of human BM neutrophils (https://www.ebi.ac.uk/biostudies/arrayexpress/studies/E-MTAB-11188), and the scTour analysis and the score of human BM neutrophils at different developmental stages was shown. **f** Pseudotime analysis of human BM neutrophils. **g** Expression levels of immunosuppressive markers of human BM neutrophils at different stages.

**Supplementary information, Fig. S2.**
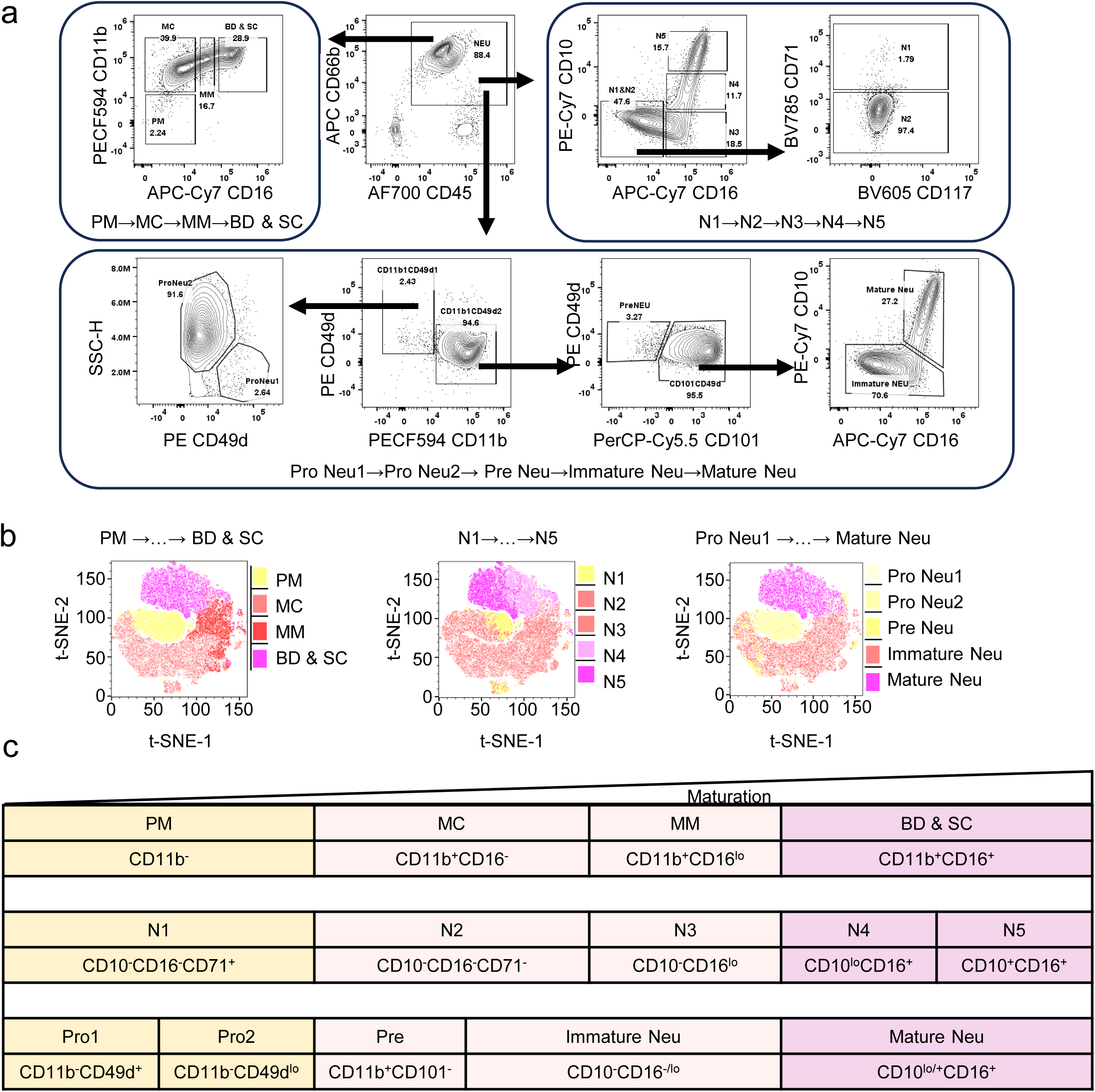
Flow cytometry gating strategy for human BM neutrophil development, related to Figure 1. **a** Flow cytometry gating strategies of human neutrophils based on three reported nomenclatures of neutrophil development in the BM of non-tumor donors. All plots gated on Lin (eFluor 506, CD3, CD19, CD56) negative. **b** BM neutrophils from non-tumor donors were collected for flow cytometry analysis, and TSNE clustering and mapping of human BM neutrophils with three distinct nomenclatures15-17 were shown. TSNE panel: FITC-CD15, PerCP-Cy5.5-CD101, PE-CD49d, PE-Cy7-CD10, PECF594-CD11b, APC-CD66b, AF700-CD45, APC-Cy7-CD16, BV421-CD34, BV510-Lin, BV605-CD117, BV785-CD71. **c** Classification of human neutrophil developmental stages with three nomenclatures.

**Supplementary information, Fig. S3.**
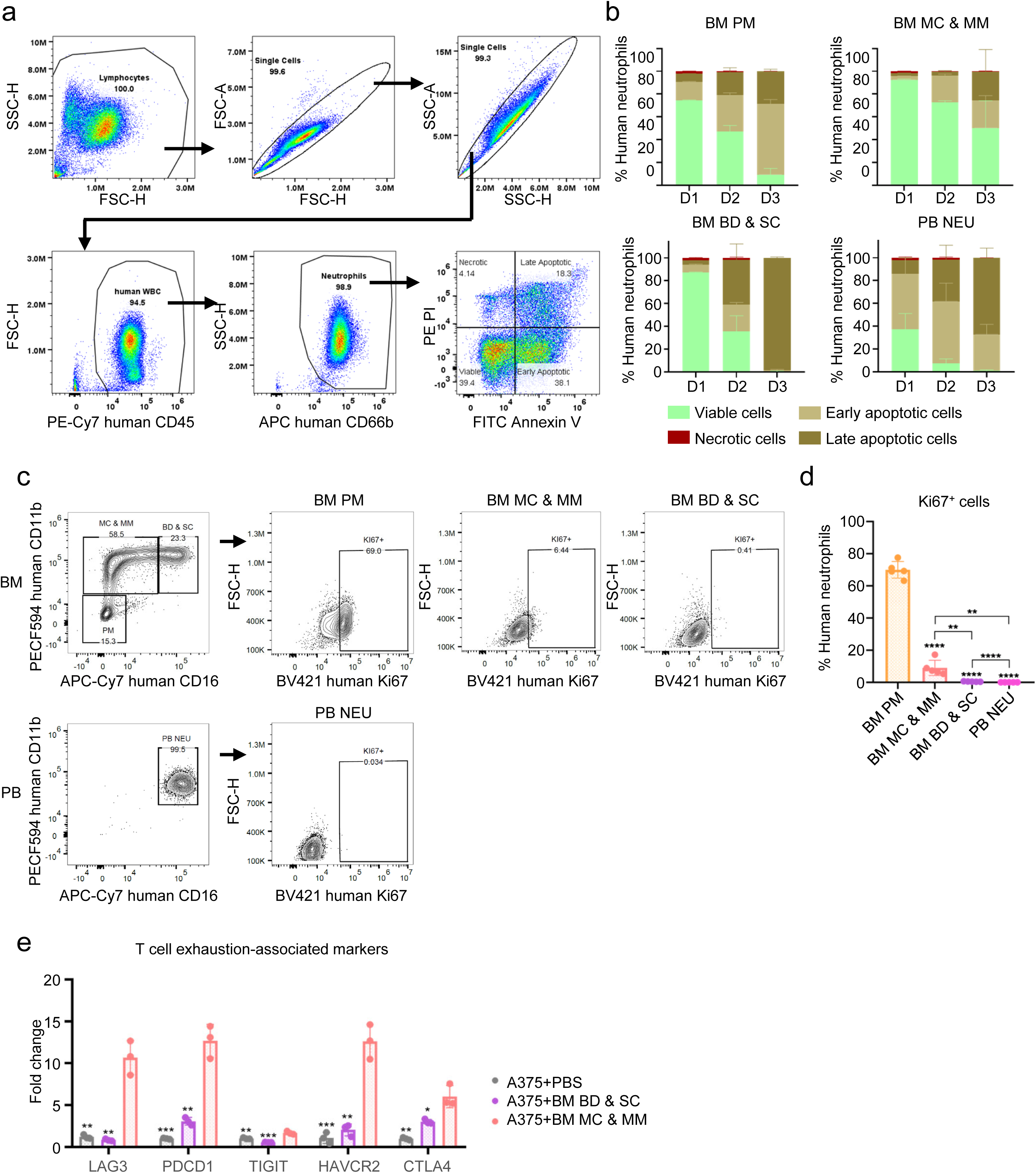
MC & MM neutrophils exhibit low Ki67 expression and enhanced resistance to apoptosis, related to Figure 2. **a** Flow cytometry gating strategies for apoptosis detection. **b** Proportions of viable, early apoptotic, late apoptotic and necrotic cells in each group were determined by flow cytometry (*n* = 5). **c** Flow cytometry gating strategies for Ki67^+^ neutrophils. **d** Proportions of Ki67^+^ cells in each group were determined by flow cytometry (*n* = 5). **e** mRNA expression of T cell exhaustion-associated markers of bulk tumor tissues analyzed by q-PCR (*n* = 3). Reference gene: human GAPDH; endogenous control: A375 + PBS group. All data are mean ± SD and were analyzed by two-tailed, unpaired Student’s t-test (\**P* < 0.05, \*\**P* < 0.01, \*\*\**P* < 0.001, \*\*\*\**P* < 0.0001).

**Supplementary information, Fig. S4.**
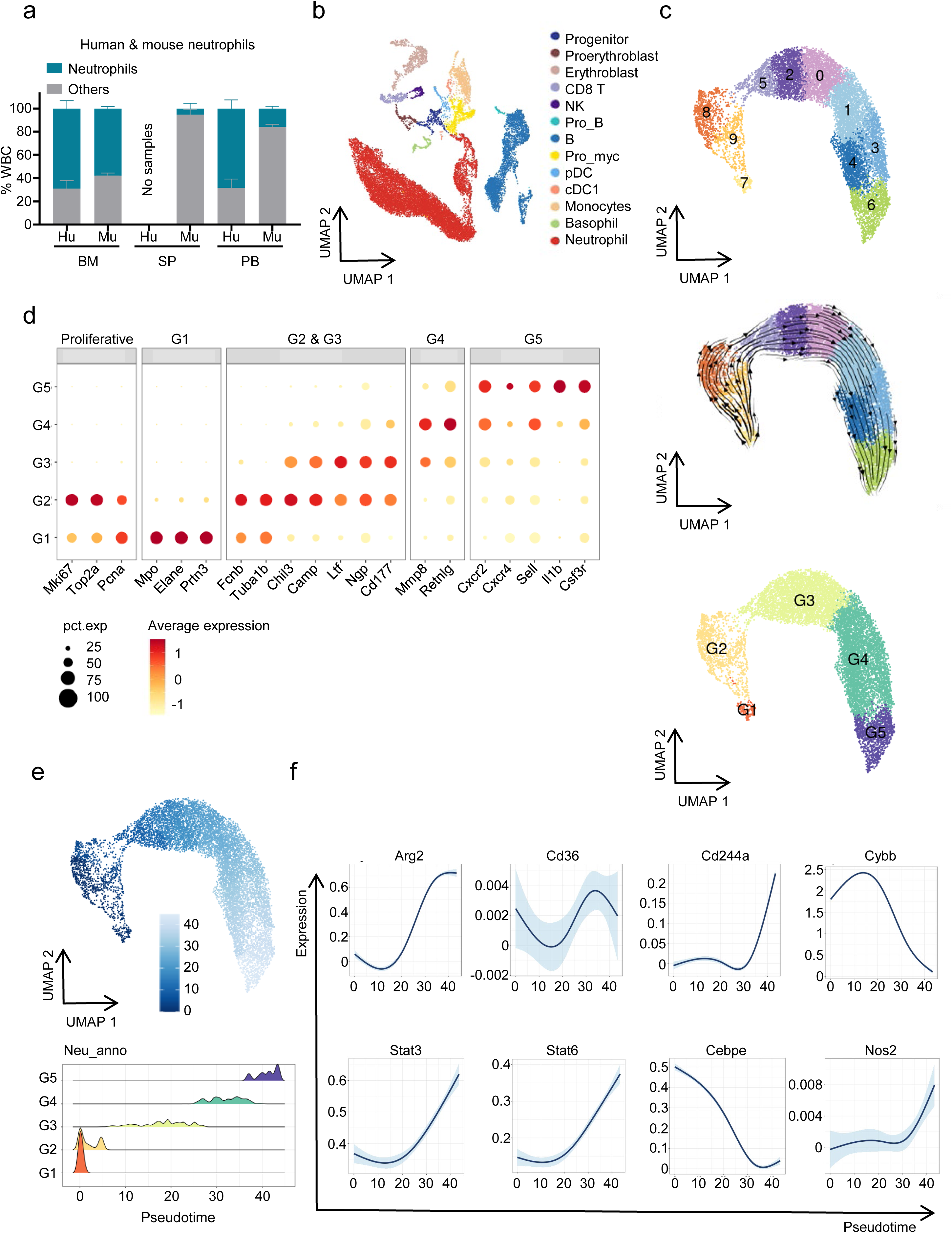
Characterization of mouse neutrophils at different developmental stages, related to Figure 1. **a** Proportions of neutrophils among WBCs in BM, SP and PB of C57BL/6J mice (*n* = 4) and human (*n* = 18). **b** UMAP plot and proportions of mouse immune cells in the BM of C57BL/6J mice. **c** scTour analysis and neutrophil scores of mouse BM neutrophils. **d** Dotplot of neutrophil markers in mouse at various stages from scRNA-seq data. **e** Pseudotime analysis of mouse BM neutrophils. **f** Expression levels of immunosuppression-related gen markers in mouse BM neutrophils.

**Supplementary information, Fig. S5.**
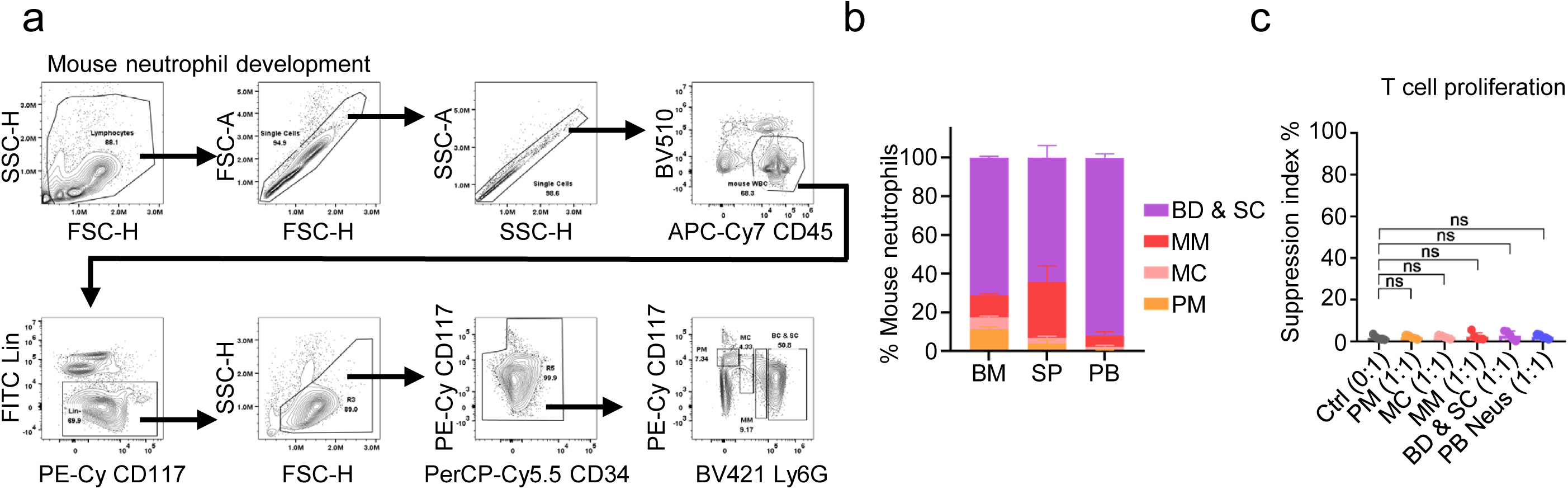
Developing neutrophils in healthy mice do not inherently possess immunosuppressive properties, related to Figure 2. **a** Flow cytometry gating strategies for mouse neutrophils at different developmental stages. **b** Proportions of mouse neutrophils at different developmental stages (*n* = 5). **c** Suppression index of mouse T cells after co-cultured with mouse neutrophils in different stages (*n* = 5). All data are mean ± SD and were analyzed by two-tailed, unpaired Student’s t-test (\**P* < 0.05, \*\**P* < 0.01, \*\*\**P* < 0.001, \*\*\*\**P* < 0.0001).

**Supplementary information, Fig. S6.**
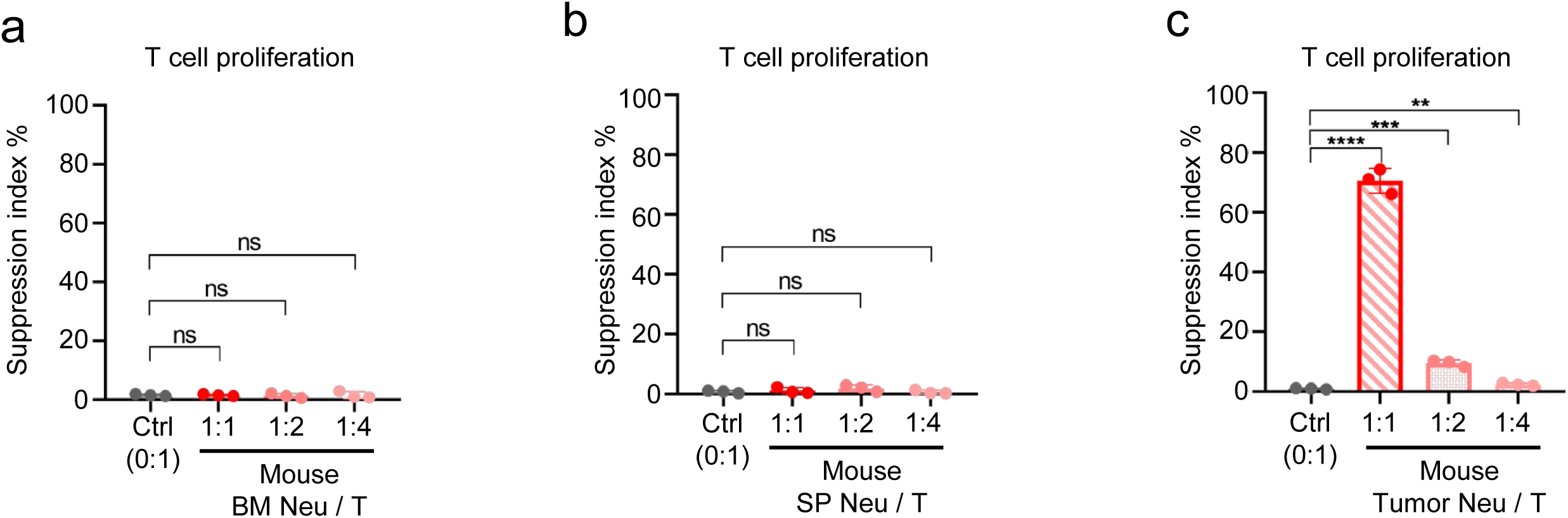
Mouse BM neutrophils have no effect on T cell proliferation, related to Figure 2. **a** Suppression index of mouse T cell proliferation co-cultured with mouse neutrophils sorted from BM of C57BL/6J mice (*n* = 3). **b** Suppression index of mouse T cell proliferation co-cultured with mouse neutrophils sorted from SP of C57BL/6J mice (*n* = 3). **c** Suppression index of mouse T cell proliferation co-cultured with mouse neutrophils sorted from prostate tumor of B6/JGpt-Pten^em1Cflox^/Gpt mice (*n* = 3). All data are mean ± SD and were analyzed by two-tailed, unpaired Student’s t-test (\**P* < 0.05, \*\**P* < 0.01, \*\*\**P* < 0.001, \*\*\*\**P* < 0.0001).

**Supplementary information, Fig. S7.**
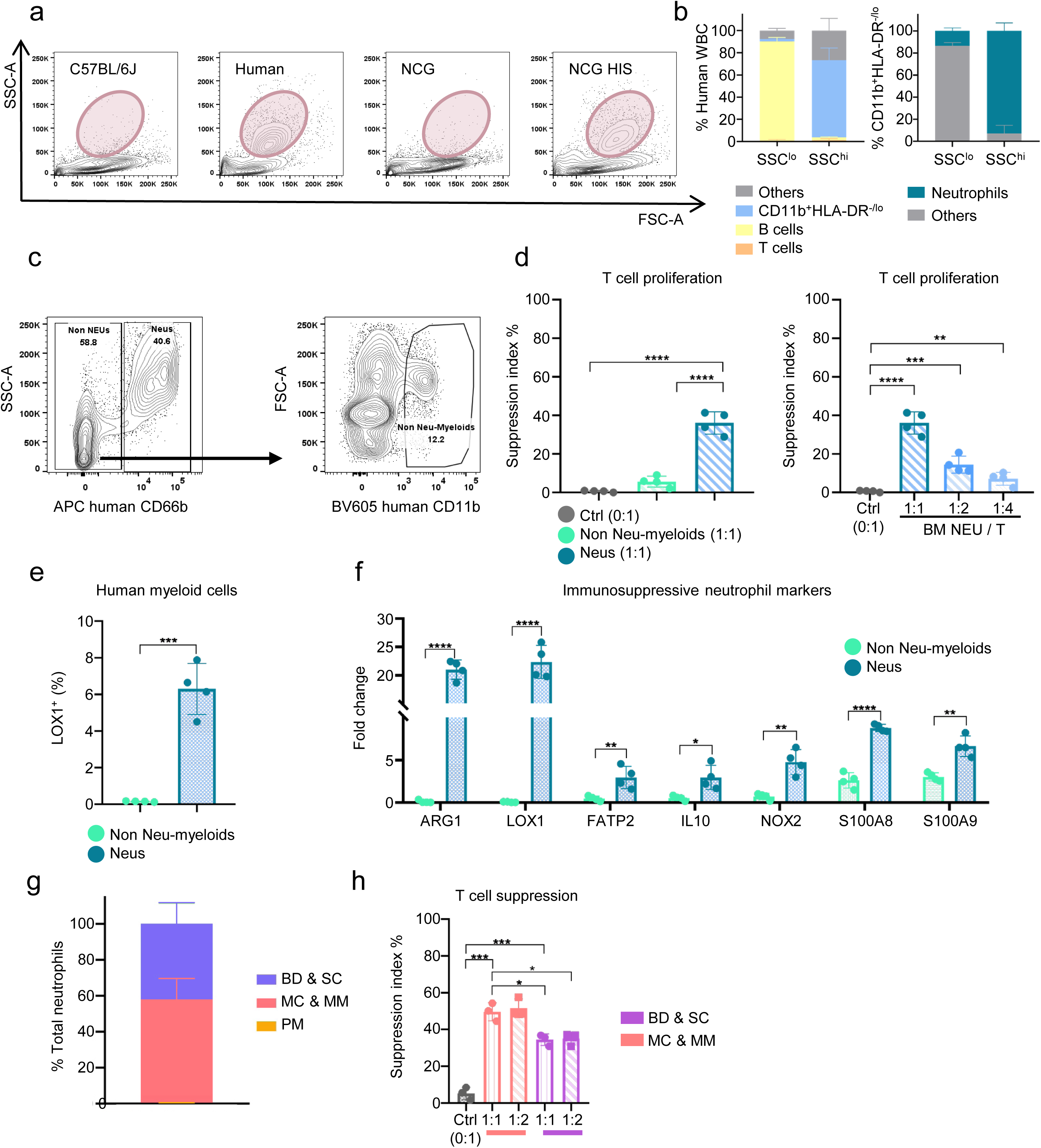
Human neutrophils in the BM of NCG HIS mice are immunosuppressive, related to Figure 2. **a** Representative flow cytometry plots of BM cells in C57BL/6J, NCG, NCG HIS mice or non-tumor donors. **b** Proportions of SSC^lo^ & SSC^hi^ cells in the BM of NCG HIS mice (*n* = 8). **c** Representative flow cytometry plots of human myeloid cell populations sorted from the BM of NCG HIS mouse. Non-neutrophilic myeloid cells: Lin^-^CD66b^-^CD11b^+^; Neutrophils: Lin^-^CD66b^+^CD11b^+^). **d** Suppression index of human T cell proliferation co-cultured with different human myeloid cell populations sorted from BM of NCG HIS mice (*n* = 3). **e** Proportions of LOX1^+^ cells among human myeloid cell populations in the BM of NCG HIS mice (*n* = 5) determined by flow cytometry. **f** mRNA expression of immunosuppressive genes in different myeloid cell populations in the BM of NCG HIS mice (*n* = 3) analyzed by q-PCR. Reference gene: human GAPDH, and endogenous control: HIS mice total BM cells. **g** Constitution of PM (CD66b^+^CD11b^-^), MC & MM (CD66b^+^CD11b^+^CD10^-^CD16^-/lo^), and BD & SC (CD66b^+^CD11b^+^CD10^lo/+^CD16^+^) neutrophils among total human neutrophils in the BM of NCG HIS mice (*n* = 14). **h** Suppression index of human T cell proliferation co-cultured with MC & MM or BD & SC neutrophils sorted from BM of NCG HIS mice (*n* = 3). All data are mean ± SD and were analyzed by two-tailed, unpaired Student’s t-test (\**P* < 0.05, \*\**P* < 0.01, \*\*\**P* < 0.001, \*\*\*\**P* < 0.0001).

**Supplementary information, Fig. S8.**
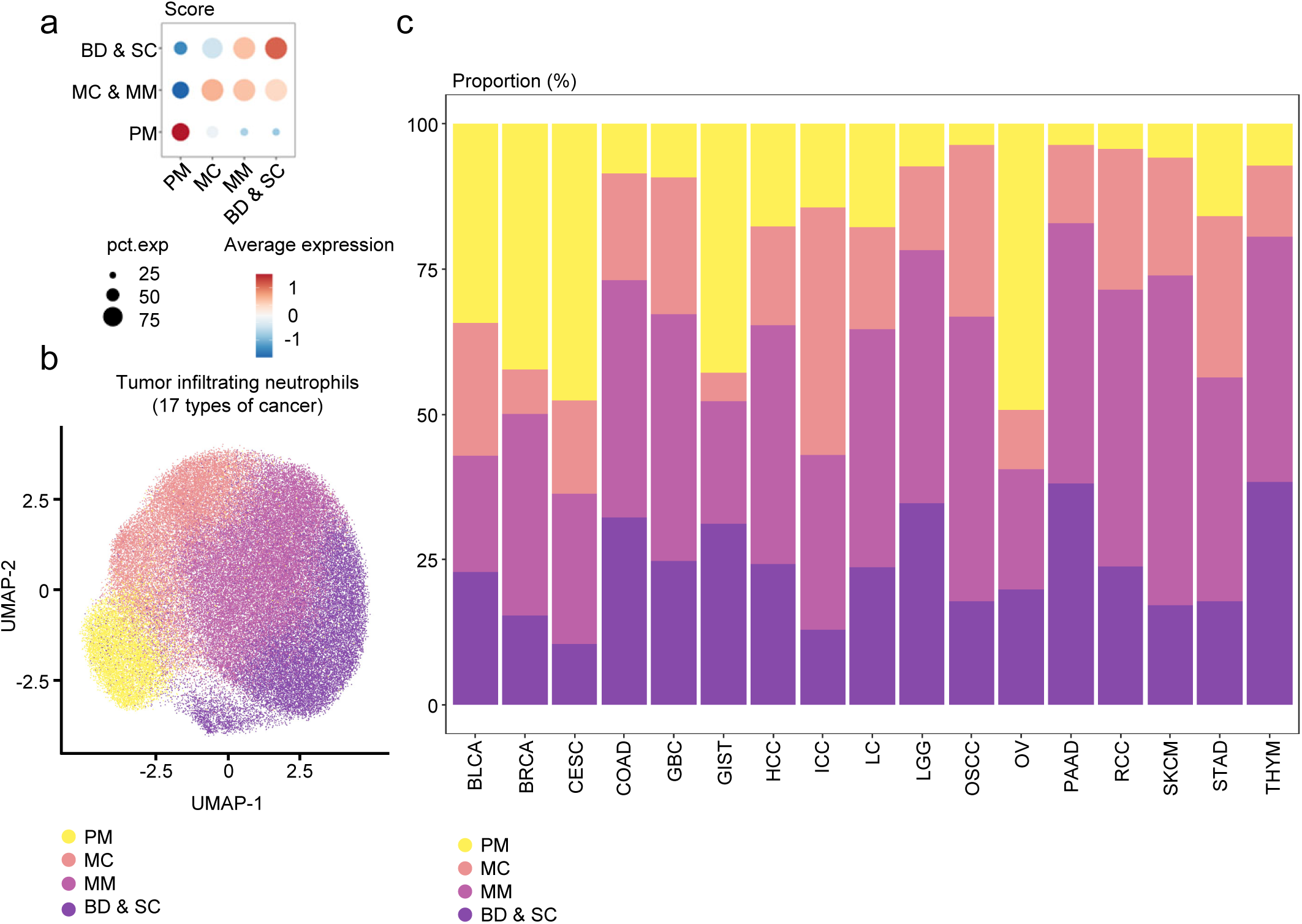
Clustering and proportion of neutrophils based on developmental stages across 17 types of cancer, related to Figure 3. **a** Analysis of neutrophil scores of TINs at different developmental stages from reported sc-RNAseq datasets containing 17 types of cancer^14^. **b** UMAP clustering of TINs at different developmental stages, data accessed from reported sc-RNAseq datasets containing 17 types of cancer^14^. **c** Proportions of TINs at different developmental stages in each cancer type. 17 cancer types include bladder urothelial carcinoma (BLCA), breast invasive carcinoma (BRCA), cervical squamous cell carcinoma and endocervical adenocarcinoma (CESC), colon adenocarcinoma (COAD), gallbladder cancer (GBC), gastrointestinal stromal tumor (GIST), hepatocellular carcinoma (HCC), intrahepatic cholangiocarcinoma (ICC), lung cancer (LC), lower grade glioma (LGG), oral squamous cell carcinoma (OSCC), ovarian serous cystadenocarcinoma (OV), pancreatic adenocarcinoma (PAAD), renal cell carcinoma (RCC), skin cutaneous melanoma (SKCM), stomach adenocarcinoma (STAD), thymoma (THYM).

**Supplementary information, Fig. S9.**
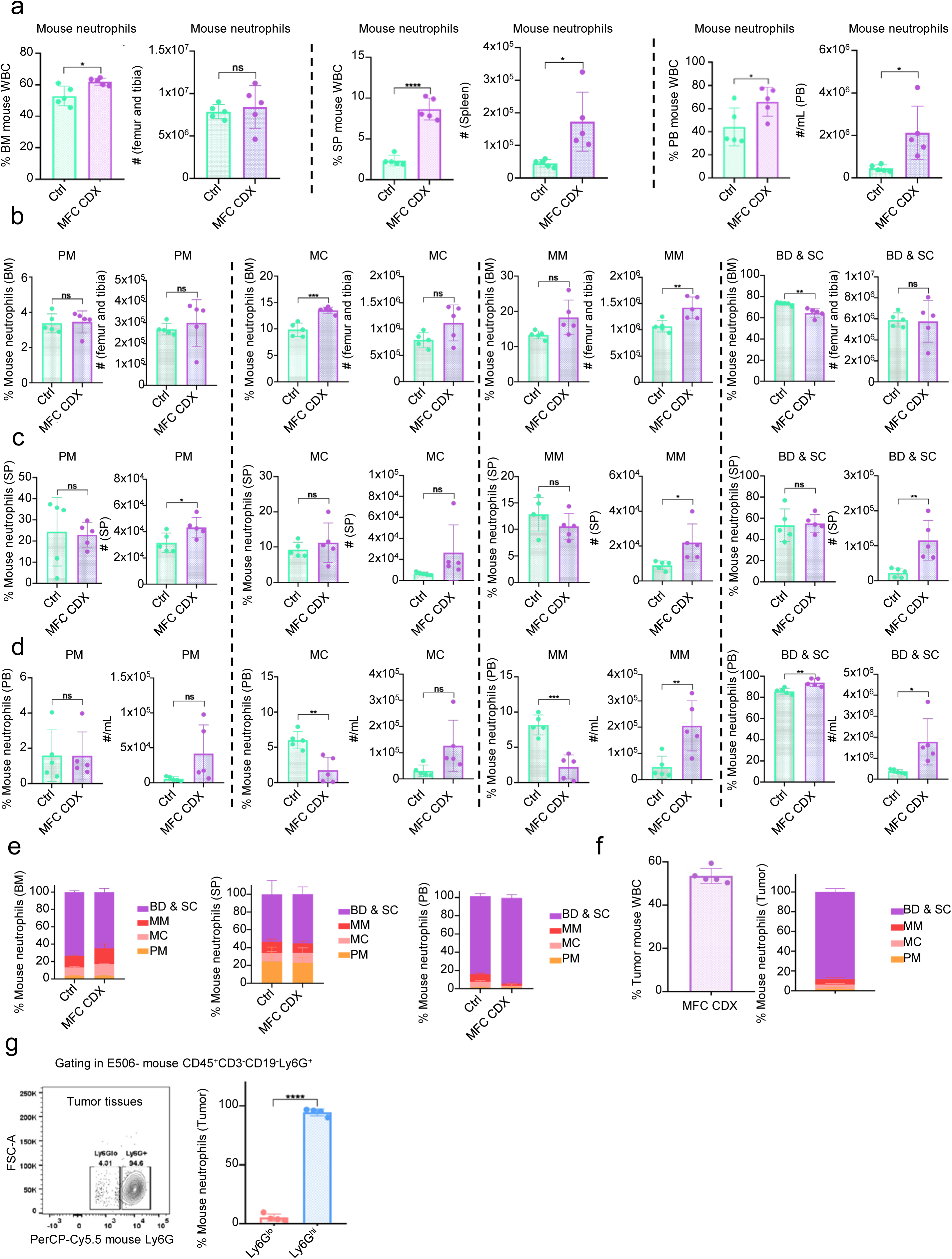
Composition of mouse neutrophils at different developmental stages in the syngeneic GC model, related to Figure 3. **a** 615-line mice were injected s.c. with 10^6^ MFC cells to establish the syngeneic GC mouse model (MFC CDX). Proportions and numbers of mouse neutrophils in BM, SP and PB of control or MFC CDX mice (*n* = 5). **b** Proportions and numbers of mouse neutrophils at different developmental stages in BM of control or MFC CDX mice (*n* = 5). **c** Proportions and numbers of mouse neutrophils at different developmental stages in SP of control or MFC CDX mice (*n* = 5). **d** Proportions and numbers of mouse neutrophils at different developmental stages in PB of control or MFC CDX mice (*n* = 5). **e** Composition of mouse neutrophils at different developmental stages in BM, SP and PB of control or MFC CDX mice (*n* = 5). **f** Proportions of mouse neutrophils at different developmental stages in tumor tissues of MFC CDX mice (*n* = 5). **g** Proportions of mouse Ly6G^+^ and Ly6G^lo^ neutrophils in prostate tumor tissues of B6/JGpt-Pten^em1Cflox^/Gpt mice (*n* = 4). All data are mean ± SD and were analyzed by two-tailed, unpaired Student’s t-test (\**P* < 0.05, \*\**P* < 0.01, \*\*\**P* < 0.001, \*\*\*\**P* < 0.0001).

**Supplementary information, Fig. S10.**
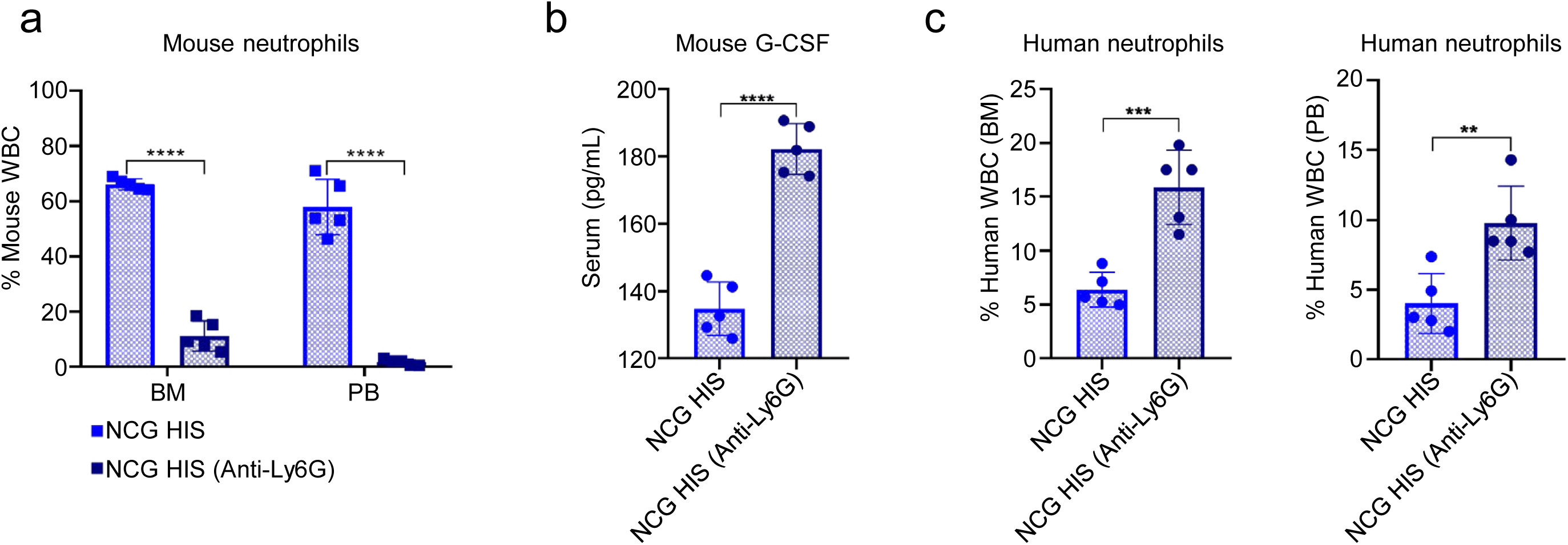
Depletion of mouse neutrophils in NCG HIS mice can promote human neutrophil reconstitution. **a** Proportions of mouse neutrophils in the BM or PB of NCG HIS (with or without anti-Ly6G treatment) mice (*n* = 5). **b** Concentration of mouse G-CSF in serum of 10 weeks old NCG HIS (with or without anti-Ly6G treatment) mice (*n* = 5). **c** Proportion of human neutrophils in the BM or PB of 10 weeks old NCG HIS (with or without anti-Ly6G treatment) mice (*n* = 5). All data are mean ± SD and were analyzed by two-tailed, unpaired Student’s t-test (\**P* < 0.05, \*\**P* < 0.01, \*\*\**P* < 0.001, \*\*\*\**P* < 0.0001).

**Supplementary information, Fig. S11.**
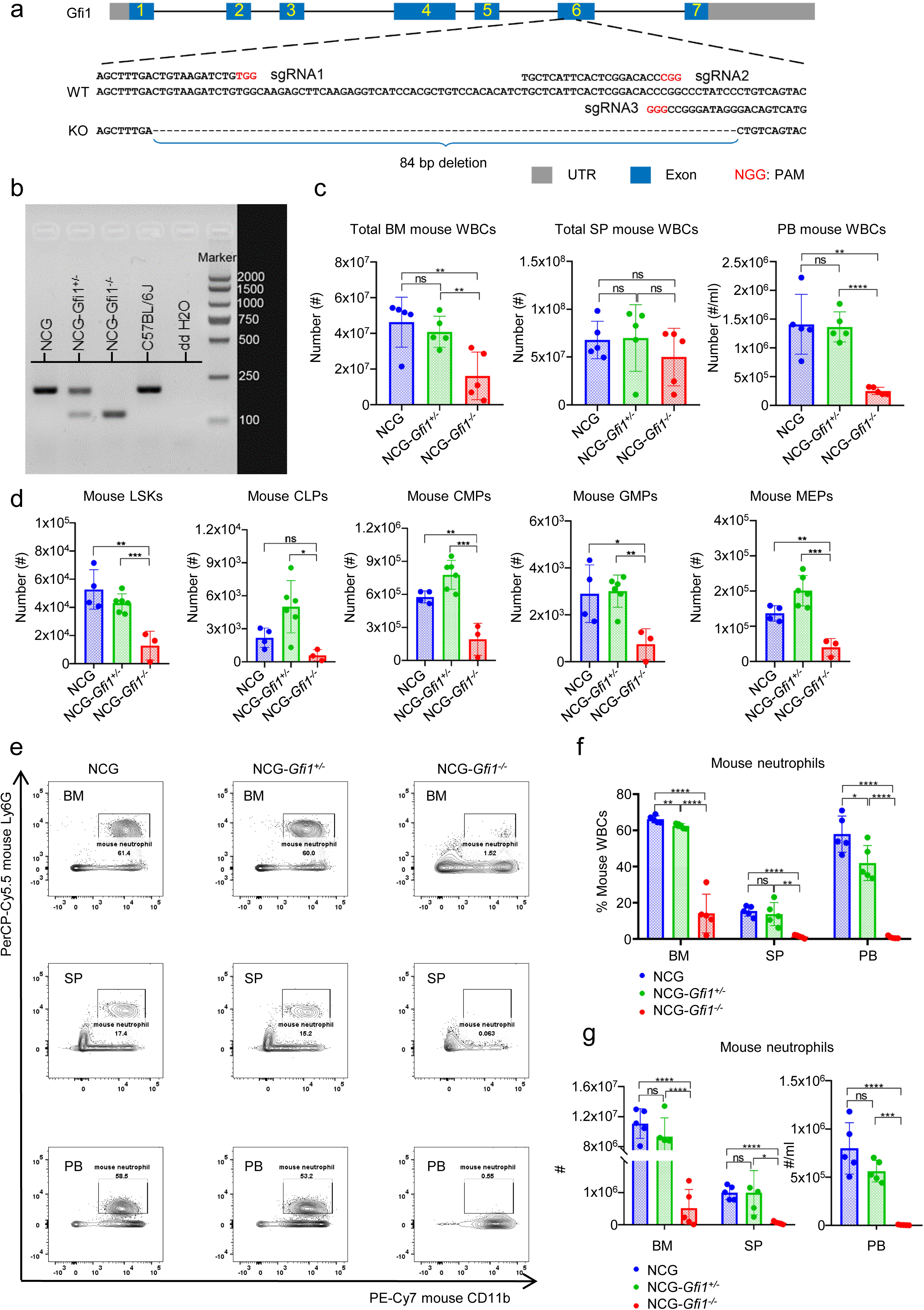
The composition of mouse WBCs in NCG-*Gfi1*^-/-^ HIS model, related to Figure 4. **a** Schematic diagram of the genetic knockout strategy for *Gfi1* using CRISPR-Cas9 in NOD Mice. **b** Representative agarose gel electrophoresis for mouse *Gfi1* gene genotyping. **c** The number of mouse WBCs in BM, SP and PB of NCG, NCG-Gif1^+/-^ and NCG-Gif1^-/-^ mice (*n* = 5). **d** The number of mouse lineage^-^Sca-1^+^c-Kit^+^ cells (LSKs), common lymphoid progenitors (CLPs), common myeloid progenitors (CMPs), granulocyte-macrophage progenitors (GMPs) and megakaryocyte-erythroid progenitors (MEPs) in BM of NCG (*n* = 4), NCG-Gif1^+/-^ (*n* = 6) and NCG-Gif1^-/-^ (*n* = 3) mice. **e** Representative flow cytometry plots of mouse neutrophils (CD11b^+^Ly6G^+^, gated on eFluor 506^-^CD45^+^) in BM, SP and PB of NCG, NCG-*Gfi1*^+/-^ and NCG-*Gfi1*^-/-^ mice (*n* = 5). **f** Proportions of mouse neutrophils in the BM, SP and PB of NCG, NCG-*Gfi1*^+/-^ and NCG-*Gfi1*^-/-^ mice (*n* = 5). **g** The number of mouse neutrophils in the BM, SP and PB of NCG, NCG-*Gfi1*^+/-^ and NCG-*Gfi1*^-/-^ mice (*n* = 5). All data are mean ± SD and were analyzed by two-tailed, unpaired Student’s t-test (\**P* < 0.05, \*\**P* < 0.01, \*\*\**P* < 0.001, \*\*\*\**P* < 0.0001).

**Supplementary information, Fig. S12.**
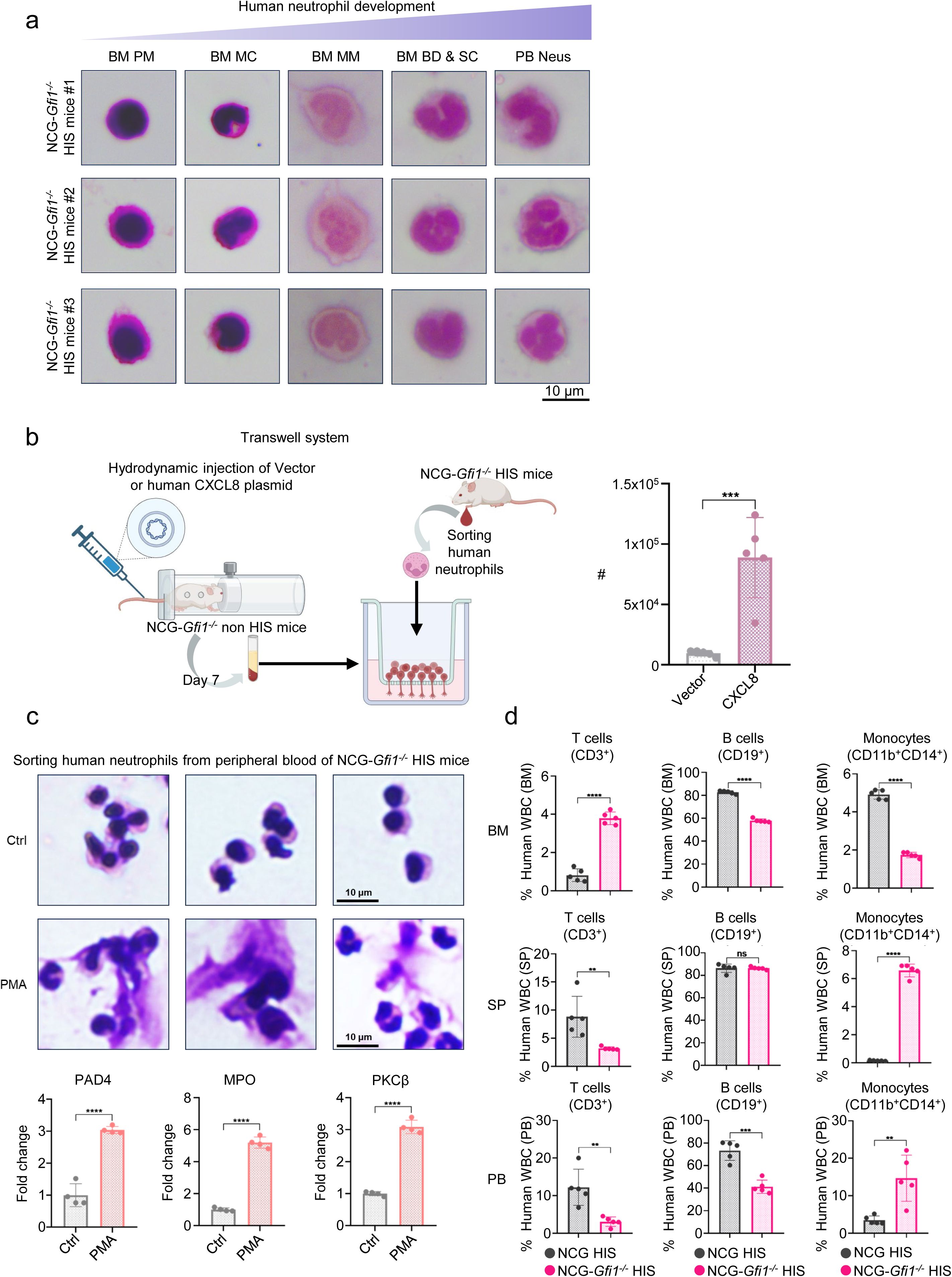
The function of human neutrophils and the composition of other immune cells in NCG-*Gfi1*^-/-^ HIS and NCG non-HIS mice, related to Figure 4. **a** Giemsa staining of BM PM∼BD & SC-stage neutrophils and PB neutrophils of NCG-*Gfi1^-/-^* HIS mice. **b** To evaluate the function of neutrophils in the peripheral blood of NCG-*Gfi1*^-/-^ HIS mice, we performed the transwell migration assay of human neutrophils sorted from NCG-*Gfi1*^-/-^ HIS mice with serum collected from NCG-*Gfi1*^-/-^ non-HIS mice hydrodynamically injected with human CXCL8 plasmid. Collected serum was incorporated it into the lower chamber of the transwell system at a 20% concentration. Human neutrophils sorted from the PB of NCG-*Gfi1*^-/-^ HIS mice were placed in the upper chamber (2 × 10⁵ cells per chamber). After 8 h, the number of neutrophils that migrated into the CXCL8-enriched lower medium was detected. **c** Giemsa staining and qPCR of human neutrophils from PB of NCG-*Gfi1*^-/-^ HIS mice after PMA treatment (*n* = 4). Reference gene: human *GAPDH*, and endogenous control: human neutrophils in control group. **d** Proportions of human T (CD3^+^), B (CD19^+^) and monocytes (CD11b^+^CD14^+^) in BM, SP and PB of 10 weeks old NCG and NCG-*Gfi1*^-/-^ HIS mice (*n* = 5). All data are mean ± SD and were analyzed by two-tailed, unpaired Student’s t-test (\**P* < 0.05, \*\**P* < 0.01, \*\*\**P* < 0.001, \*\*\*\**P* < 0.0001).

**Supplementary information, Fig. S13.**
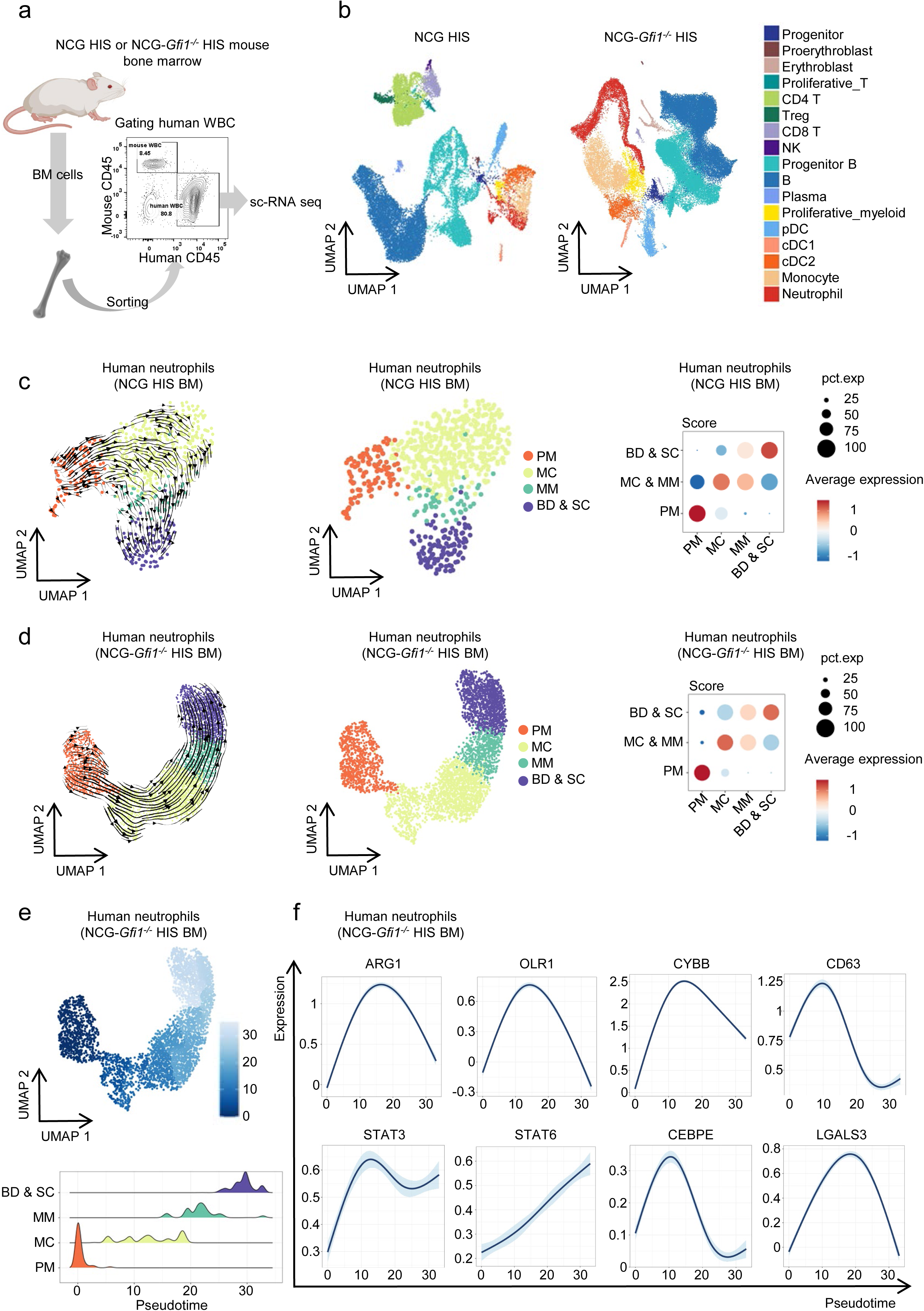
Analysis of MC & MM neutrophils from the BM of NCG HIS or NCG-*Gfi1*^-/-^ HIS mice, related to Figure 5. **a** Schematic of human immune cells (E506^-^ mCD45^-^ hCD45^+^) from BM of NCG HIS or NCG-*Gfi1*^-/-^ HIS mice were sorted for scRNA-seq analysis. **b** UMAP clustering of human immune cells from BM of NCG HIS or NCG-*Gfi1*^-/-^ HIS mice. **c** scTour analysis, UMAP clustering, and neutrophil scores at different developmental stages in the BM of NCG HIS mice. **d** scTour analysis, UMAP clustering, and neutrophil scores at different developmental stages in the BM of NCG-*Gfi1*^-/-^ HIS mice. **e** UMAP clustering and pseudotime analysis of human neutrophils from scRNA-seq dataset of the BM of NCG-*Gfi1*^-/-^ HIS mice. **f** Expression levels of immunosuppressive markers of NCG-*Gfi1*^-/-^ HIS mouse BM neutrophils at different stages.

**Supplementary information, Fig. S14.**
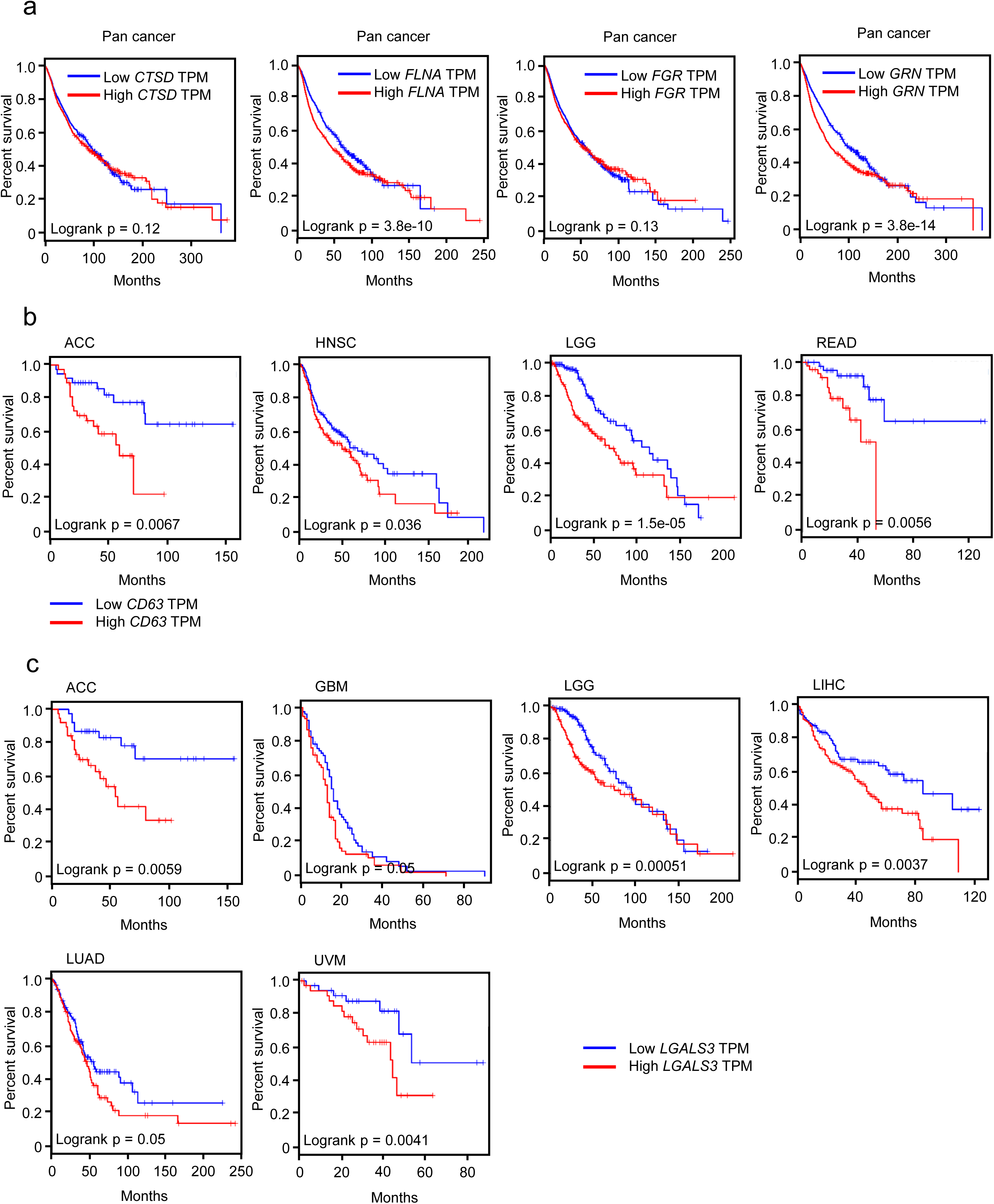
Survival analysis of DEGs of MC & MM neutrophils in various cancers from TCGA, related to Figure 5. **a** Survival analysis of patients with 32 cancer types (pan-cancer) from TCGA with low or high *CTSD*, *FLNA*, *FGR*, or *GRN* expression. **b** Survival analysis of cancer patients from TCGA with low or high *CD63* expression. ACC, Adrenocortical carcinoma; HNSC, Head and neck squamous cell carcinoma; LGG, Brain lower grade glioma; READ, Rectum adenocarcinoma. **c** Survival analysis of cancer patients from TCGA with low or high *LGALS3* expression. ACC, Adrenocortical carcinoma; GBM, Glioblastoma multiforme; LIHC, Liver hepatocellular carcinoma; LUAD, Lung adenocarcinoma; UVM, Uveal melanoma. Survival curves were analyzed by Log-rank test (\**P* < 0.05, \*\**P* < 0.01, \*\*\**P* < 0.001, \*\*\*\**P* < 0.0001).

**Supplementary information, Fig. S15.**
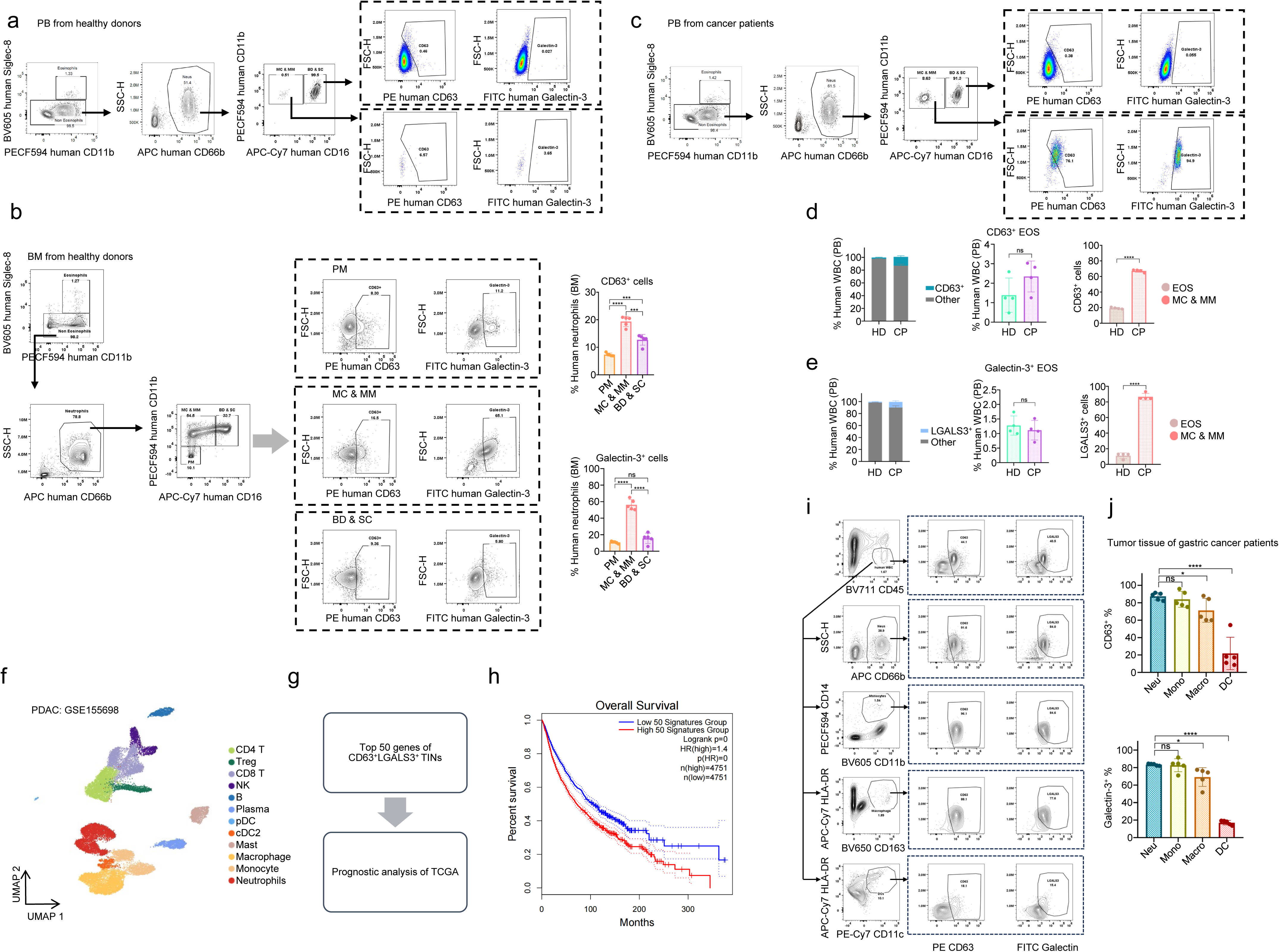
Flow cytometry gating strategies for analysis of CD63 and Galectin-3 expression in human BD & SC and MC & MM neutrophils, related to Figure 5. **a** Flow cytometry gating strategy for the analysis of CD63 and Galectin-3 expression on neutrophils in PB of healthy donors (HD). **b** Flow cytometry gating strategy for the analysis of CD63 and Galectin-3 expression on neutrophils at different developmental stages in BM (*n* = 5). **c** Flow cytometry gating strategy for the analysis of CD63 and Galectin-3 expression on neutrophils in PB of cancer patients (CP). **d** Proportions of CD63^+^ populations among eosinophils (EOS), MC & MM and BD & SC neutrophils in the PB of HD or CP (*n* = 4). **e** Proportions of Galectin-3^+^ populations among eosinophils (EOS), MC & MM and BD & SC neutrophils in the PB of HD or CP (*n* = 4). **f** UMAP plot of tumor-infiltrating immune cells (PDAC, GSE155698). **g** We utilized scRNAseq data from the PDAC tumor tissue to calculate top 50 genes that are highly expressed in CD63^+^LGALS3^+^ neutrophils and combined them with TCGA pan-cancer survival data to analyze the correlation between the gene signature (quantified as a composite score) and prognosis. **h** Survival analysis of TCGA pan-cancer patients with high or low CD63^+^LGALS3^+^ neutrophil-associated gene signature expression. **i** Flow cytometry Gating strategy for analysis of CD63 and Galectin-3 expression on neutrophils (SSC^hi^CD66b^+^), monocytes (CD11b^+^CD14^+^), macrophages (HLA-DR^+^CD163^+^) and DCs (HLA-DR^+^CD11c^+^) in tumor tissues of gastric cancer patients. **j** Proportions of CD63^+^ or Galectin-3^+^ populations among neutrophils, monocytes, macrophages and DCs in tumor tissues of gastric cancer patients (*n* = 5). All data are mean ± SD and were analyzed by two-tailed, unpaired Student’s t-test (\**P* < 0.05, \*\**P* < 0.01, \*\*\**P* < 0.001, \*\*\*\**P* < 0.0001).

**Supplementary information, Fig. S16.**
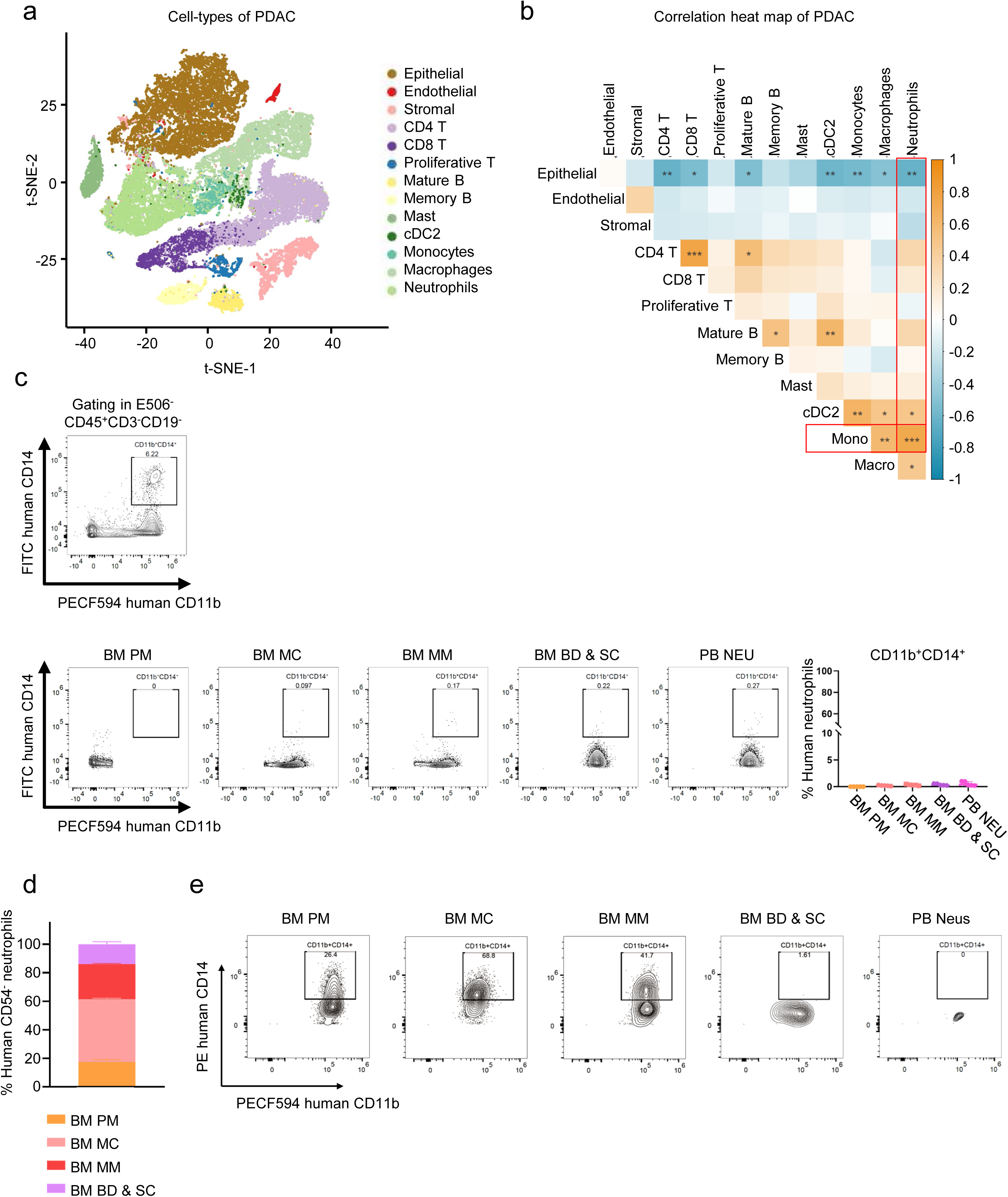
Correlation analysis of different cell types in the TIME of PDAC patients and the trans-differentiation potential of human BM neutrophils, related to Figure 6. **a** t-SNE clustering of cell types in the tumors of PDAC patients^35^. **b** Heatmap depicting the correlation of cell types in the tumors of PDAC patients. **c** Flow cytometry Gating strategy for analysis of CD11b^+^CD14^+^ expression on neutrophils in different stages of human BM and PB (*n* = 5). **d** The developmental stage distribution of human CD54^-^ neutrophils in the BM (*n* = 5). **e** After culture with Flt3L for 4 days, proportions of CD11b^+^CD14^+^ cells in each group were determined by flow cytometry (*n* = 5). All data are mean ± SD and were analyzed by two-tailed, unpaired Student’s t-test (\**P* < 0.05, \*\**P* < 0.01, \*\*\**P* < 0.001, \*\*\*\**P* < 0.0001).

**Supplementary information, Fig. S17.**
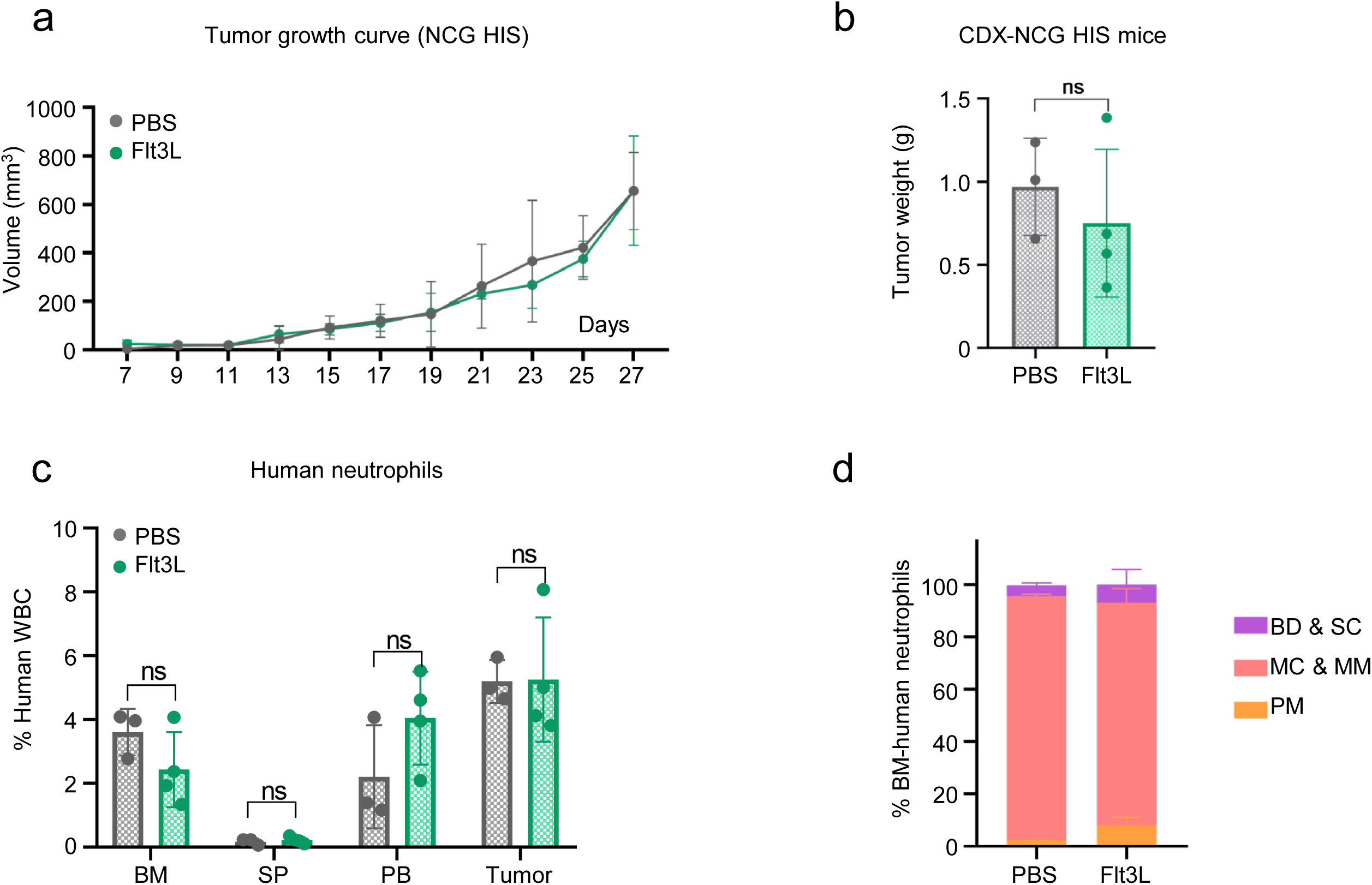
Flt3L has no tumor-control effect on CDX-NGC HIS mice, related to Figure 7. **a** A375 xenograft NCG HIS model was established, and Flt3L plasmid was injected i.v. hydrodynamically. Tumor growth of NCG HIS mice with PBS or Flt3L treatment (PBS, *n* = 3; Flt3L, *n* = 3). **b** Tumor weight of NCG HIS mice with PBS or Flt3L treatment (PBS, *n* = 3; Flt3L, *n* = 3). **c** Proportions of neutrophils (CD11b^+^CD66b^+^) in the BM, SP, PB and tumors of NCG HIS mice with or without Flt3L treatment (*n* = 3). **d** Proportions of human neutrophils at different developmental stages in the BM of NCG HIS mice with or without Flt3L treatment (*n* = 3). All data are mean ± SD and were analyzed by two-tailed, unpaired Student’s t-test (\**P* < 0.05, \*\**P* < 0.01, \*\*\**P* < 0.001, \*\*\*\**P* < 0.0001).

**Supplementary information, Fig. S18.**
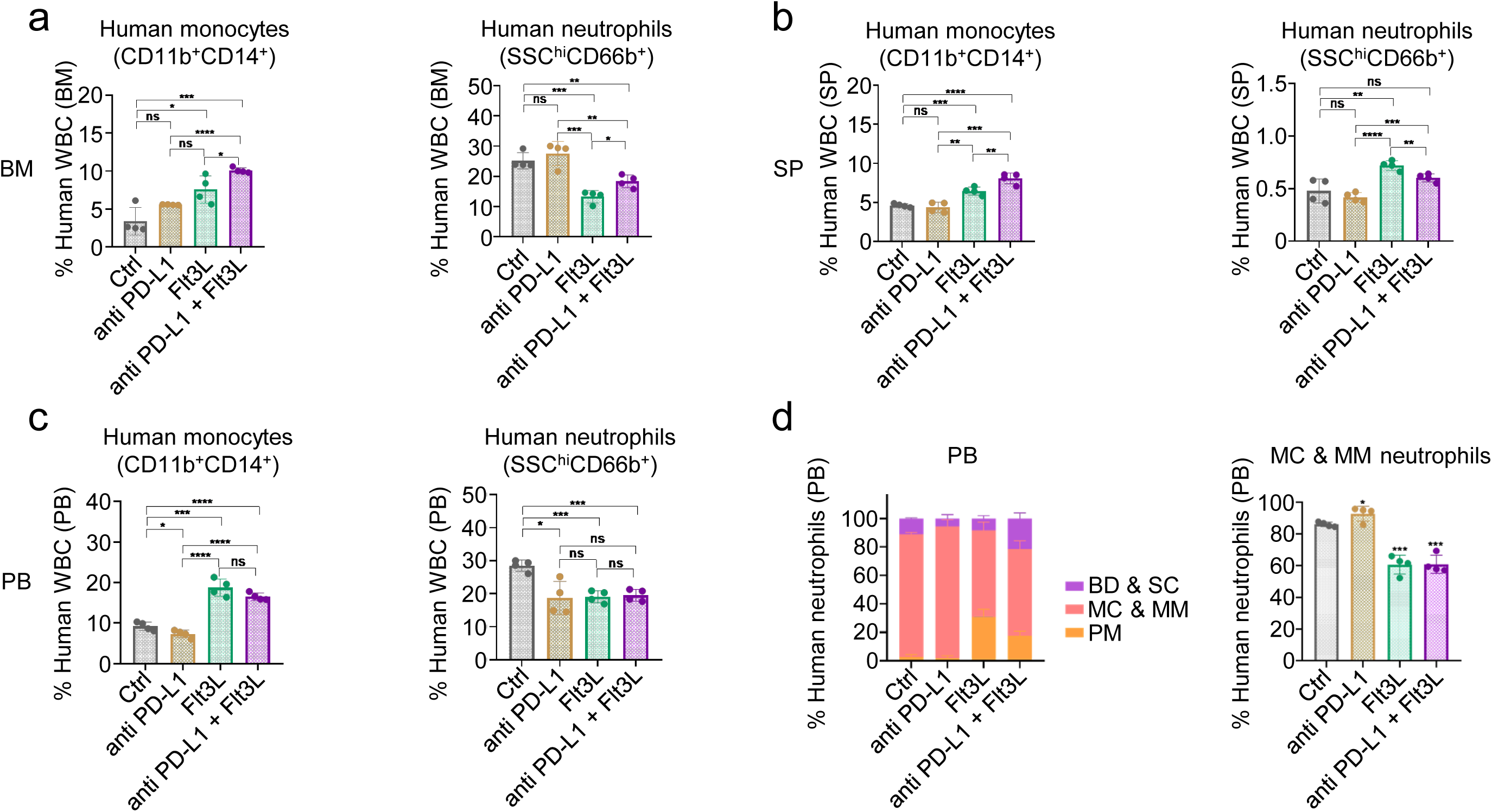
Flt3L treatment increased the proportion of human CD11b^+^CD14^+^ monocytes, related to Figure 7. **a** Proportions of human monocytes (CD11b^+^CD14^+^) and neutrophils (SSC^hi^CD66b^+^) and neutrophils at different developmental stages in the BM of CDX NCG-*Gfi1*^-/-^ HIS mice from each treatment group (*n* = 4). **b** Proportions of human monocytes (CD11b^+^CD14^+^) and neutrophils (SSC^hi^CD66b^+^) and neutrophils at different developmental stages in the spleen of CDX NCG-*Gfi1*^-/-^ HIS mice from each treatment group (*n* = 4). **c** Proportions of human monocytes (CD11b^+^CD14^+^) and neutrophils (SSC^hi^CD66b^+^) and neutrophils at different developmental stages in the PB of CDX NCG-*Gfi1*^-/-^ HIS mice from each treatment group (*n* = 4). **d** Proportions of human neutrophils at different developmental stages in PB of NCG-*Gfi1^-/-^* HIS mice with anti-PDL1 or Flt3L treatment (*n* = 4). All data are mean ± SD and were analyzed by two-tailed, unpaired Student’s t-test (\**P* < 0.05, \*\**P* < 0.01, \*\*\**P* < 0.001, \*\*\*\**P* < 0.0001).

**Supplementary information, Table S1.**
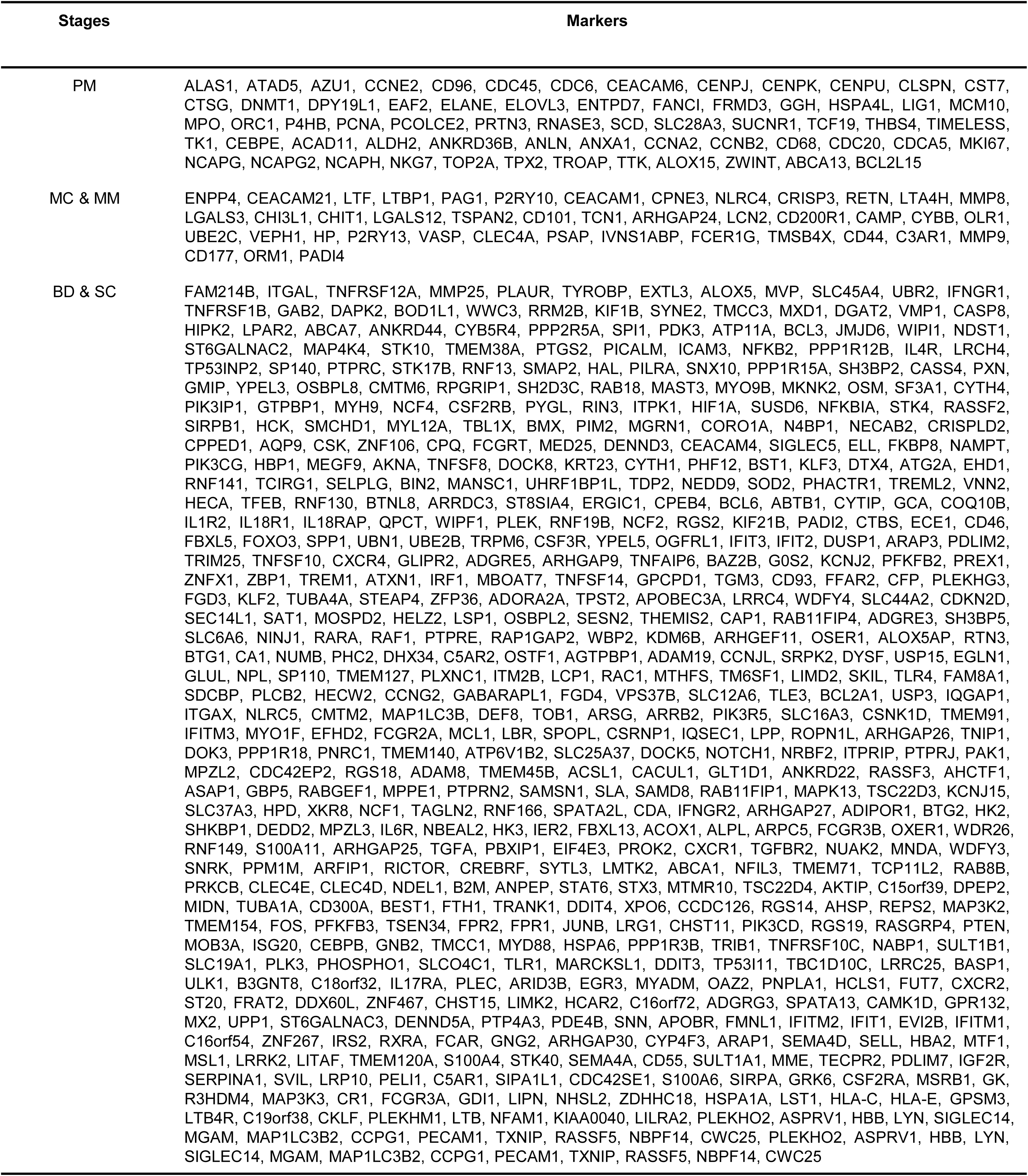
Neutrophil Score for human neutrophil stage classification.

**Supplementary information, Table S2.**
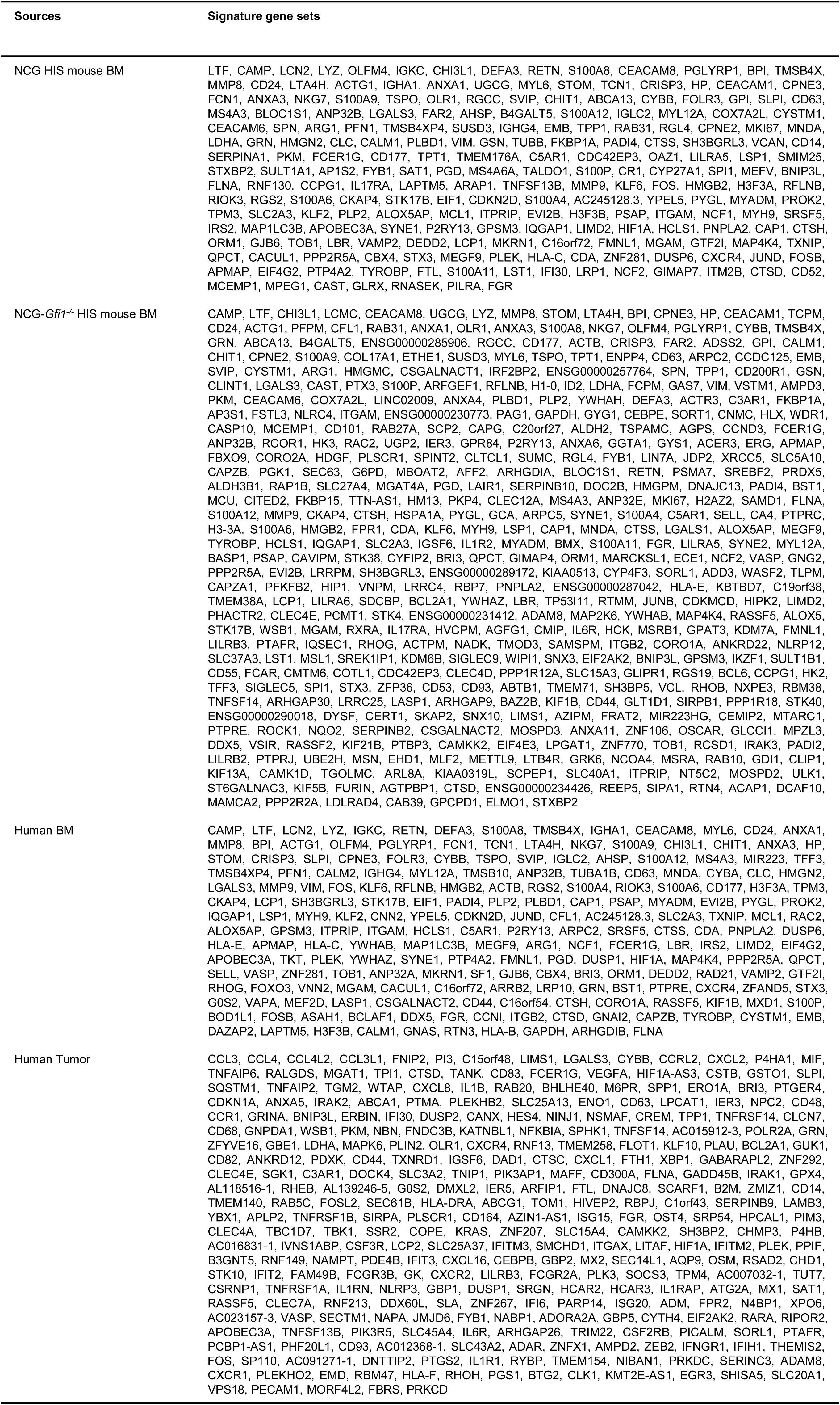
Signature gene sets of human neutrophils from multiple sources at MC & MM stages.

**Supplementary information, Table S3.** Tumor-infiltrating neutrophil (TIN) cell counts by stage in 17 tumor types.

| Cancer types | PM | MC | MM | BD & SC |
| --- | --- | --- | --- | --- |
| BLCA | 12 | 8 | 7 | 8 |
| BRCA | 11 | 2 | 9 | 4 |
| CESC | 1563 | 527 | 847 | 343 |
| COAD | 1097 | 2373 | 5328 | 4124 |
| GBC | 883 | 2242 | 4039 | 2353 |
| GIST | 131 | 15 | 64 | 95 |
| HCC | 1076 | 1029 | 2493 | 1462 |
| ICC | 485 | 1439 | 1011 | 435 |
| LC | 531 | 523 | 1218 | 700 |
| LGG | 259 | 510 | 1535 | 1221 |
| OSCC | 116 | 925 | 1533 | 556 |
| OV | 339 | 70 | 142 | 136 |
| PAAD | 334 | 1205 | 4009 | 3414 |
| RCC | 485 | 2665 | 5261 | 2617 |
| SKCM | 51 | 173 | 488 | 146 |
| STAD | 1698 | 2968 | 4120 | 1899 |
| THYM | 389 | 660 | 2276 | 2066 |

**Supplementary information, Table S4.** Flow cytometry antibody panel.

| Antibody | Source | Identifier | Dilution |
| --- | --- | --- | --- |
| Anti-human CD10 (Clone: HI10a) - FITC | Biolegend | Cat#: 312208 | 1:50 |
| Anti-human CD14 (Clone: HCD14) - FITC | Biolegend | Cat#: 325604 | 1:50 |
| Anti-human CD15 (Clone: HI98) - FITC | Ebioscience | Cat#: 11-0159-42 | 1:50 |
| Anti-mouse CD3e (Clone: BM10-37) - FITC | BD | Cat#: 553061 | 1:100 |
| Anti-mouse CD49b (Clone: DX5) - FITC | Ebioscience | Cat#: 11-5971-82 | 1:100 |
| Anti-human Galectin-3 (Clone: B2C10) - FITC | BD | Cat#: 565677 | 1:50 |
| Anti-human CD3 (Clone: UCHT1) - PerCP/Cy5.5 | Biolegend | Cat#: 300430 | 1:50 |
| Anti-human CD101 (Clone: BB27) - PerCP/Cy5.5 | Biolegend | Cat#: 331015 | 1:50 |
| Anti-human IFNγ (Clone: 4S.B3) - PerCP/Cy5.5 | Biolegend | Cat#: 502526 | 1:50 |
| Anti-mouse Ly6G (Clone: 1A8) - PerCP/Cy5.5 | BD | Cat#: 560602 | 1:100 |
| Anti-mouse CD34 (Clone: HM34) - PerCP/Cy5.5 | Biolegend | Cat#: 128607 | 1:100 |
| Anti-human Arginase I (Clone: 14D2C43) - PE | Biolegend | Cat#: 369704 | 1:50 |
| Anti-human CD19 (Clone: HIB19) - PE | Biolegend | Cat#: 302208 | 1:50 |
| Anti-human CD38 (Clone: HIT2) - PE | Biolegend | Cat#: 303506 | 1:50 |
| Anti-human CD44 (Clone: IM7) - PE | Ebioscience | Cat#: 12-0441-82 | 1:50 |
| Anti-human CD49d (Clone: 9F10) - PE | Ebioscience | Cat#: 12-0499-42 | 1:50 |
| Anti-human CD63 (Clone: H5C6) - PE | BD | Cat#: 561925 | 1:50 |
| Anti-human CD10 (Clone: HI10a) - PE/Cy7 | Biolegend | Cat#: 312214 | 1:50 |
| Anti-human/mouse CD11b (Clone: M1/70) - PE/Cy7 | Biolegend | Cat#: 101216, | 1:50 |
| Anti-human CD11c (Clone: 3.9) - PE/Cy7 | Ebioscience | Cat#: 25-0116-42 | 1:50 |
| Anti-human CD45 (Clone: 2D1) - PE/Cy7 | Biolegend | Cat#: 368532 | 1:50 |
| Anti-human CD11b (Clone: ICFR44) - PECF594 | BD | Cat#: 562399 | 1:50 |
| Anti-human/mouse CD11b (Clone: M1/70) - PECF594 | BD | Cat#: 562287 | 1:50 |
| Anti-human CD14 (Clone: MOP9) - PECF594 | BD | Cat#: 562335 | 1:50 |
| Anti-human CD33 (Clone: WM53) - PECF594 | Biolegend | Cat#: 303432 | 1:50 |
| Anti-human CD19 (Clone: HIB19) - PECF594 | BD | Cat#: 562294 | 1:50 |
| Anti-human Siglec-8 (Clone: 7C9) - PECF594 | Biolegend | Cat#: 347110 | 1:50 |
| Anti-human CD66b (Clone: G10F5) - APC | Biolegend | Cat#: 305110 | 1:50 |
| Anti-human HLA-DR (Clone: TU36) - APC | BD | Cat#: 559868 | 1:50 |
| Anti-human CD163 (Clone: GHI/61) - APC | Biolegend | Cat#: 333609 | 1:50 |
| Anti-human CD45 (Clone: HI30) - AF700 | BD | Cat#: 560566 | 1:50 |
| Anti-human CD16 (Clone: 3G8) - APC/Cy7 | BD | Cat#: 557758 | 1:50 |
| Anti-human HLA-DR (Clone: L243) - APC/Cy7 | Biolegend | Cat#: 307618 | 1:50 |
| Anti-human CD34 (Clone: 581) - BV421 | BD | Cat#: 562577 | 1:50 |
| Anti-human CD3 (Clone: HIT3a) - BV510 | BD | Cat#: 564713 | 1:50 |
| Anti-human CD19 (Clone: SJ25C1) - BV510 | BD | Cat#: 562947 | 1:50 |
| Anti-human CD56 (Clone: NCAM16.2) - BV510 | BD | Cat#: 563041 | 1:50 |
| Anti-mouse CD45 (Clone: REA737) - BV510 | Miltenyi Biotec | Cat#: 130-110-665 | 1:100 |
| Anti-human/mouse CD11b (Clone: M1/70) - BV605 | Biolegend | Cat#: 101257 | 1:50 |
| Anti-human CD14 (Clone: M5E2) - BV605 | Biolegend | Cat#: 301834 | 1:50 |
| Anti-human CD117 (Clone: 104D2) - BV605 | Biolegend | Cat#: 313218 | 1:50 |
| Anti-human CD33 (Clone: WM53) - BV650 | Biolegend | Cat#: 303430 | 1:50 |
| Anti-human CD45 (Clone: HI30) - BV711 | Biolegend | Cat#: 304050 | 1:50 |
| Anti-human CD71 (Clone: M-A712) - BV785/786 | BD | Cat#: 563768 | 1:50 |
| Anti-human CD16 (Clone: 3G8) - BUV395 | BD | Cat#: 563785 | 1:50 |
| Anti-mouse CD45 (Clone: 30-F11) - BUV395 | BD | Cat#: 564279 | 1:100 |
| Fixable viability dye eFlour506 (e506) | Invitrogen | Cat#: 65-0866-18 | 1:400 |

**Supplementary information, Table S5.**
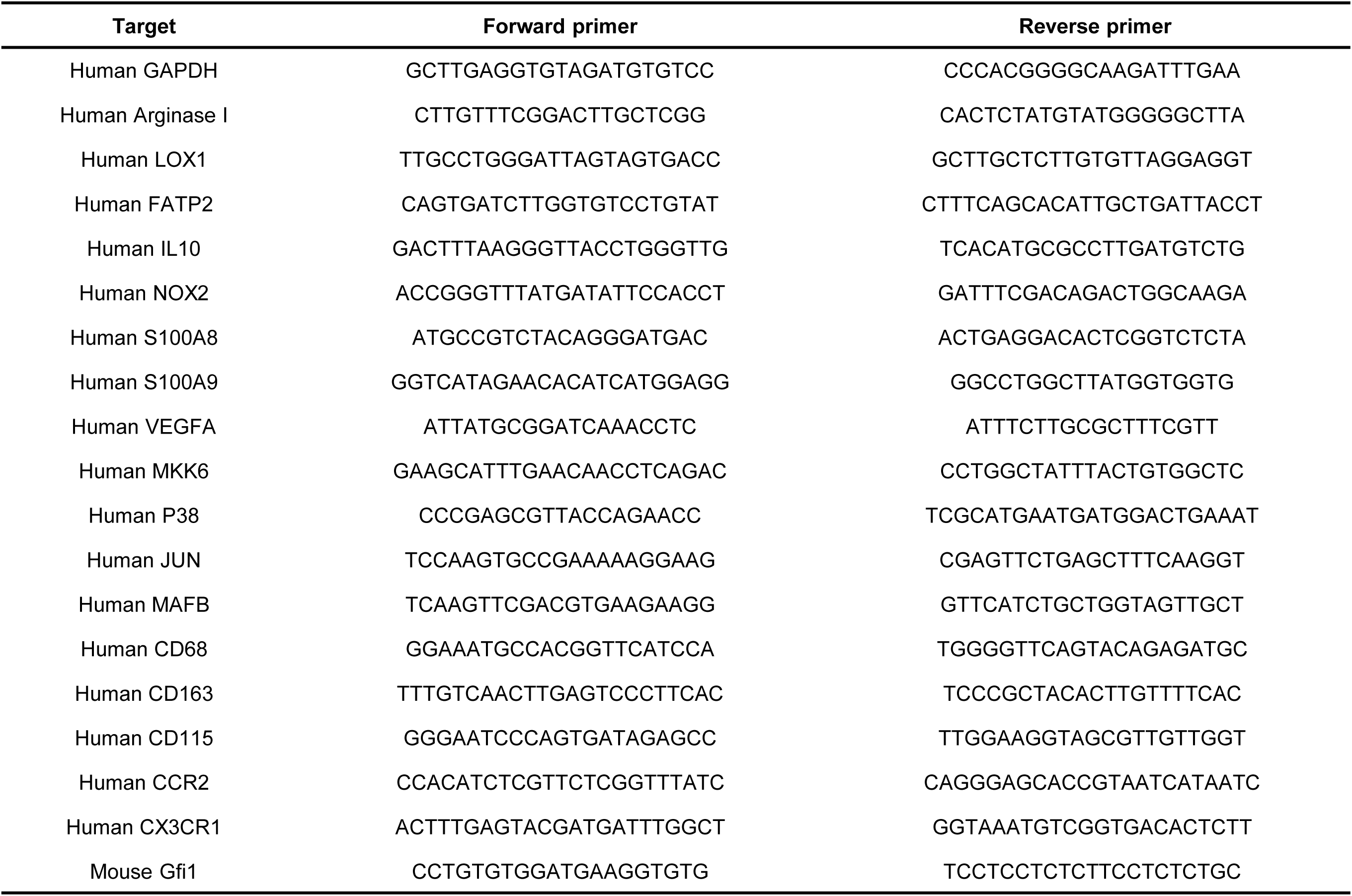
Primers used in this study.

